# A cross-species single-cell spatial atlas of kidney allografts

**DOI:** 10.64898/2026.11.19.767827

**Authors:** Jiaheng Wu, Liying Miao, Yufeng Zhao, Siyue Chen, Rui Shi, Boyang Yu, Zhengqing Lu, Chengwei Wang, Xiaoming Song, Xiaohan Yu, Ruiming Rong, Cheng Yang, Guoguo Shang, Tongyu Zhu, Dong Zhu

## Abstract

Kidney allografts with the same Banff diagnosis differ in their immune microenvironments and functional recovery after treatment, yet the tissue features distinguishing recovery from non-recovery remain unclear. We constructed SPECTRA (Spatial Etiology and Coordination of Transplant Rejection in Allografts), which comprises spatial multi-omic data from 1,011,984 cells in 83 human allograft biopsies. We also profiled 442,320 cells from 34 mouse grafts across experimental groups and time points by single-cell sequencing, and integrated a public human single-cell reference of 352,911 cells. Unsupervised analysis identified nine tissue niches, of which plasma-cell effector niche occupancy was associated with less eGFR improvement approximately 90 days after biopsy (ρ = −0.508, q = 0.0165). Plasma cells in this niche expressed IGHG1, IGKC and TXNDC5; in scarred interstitium, plasma-cell XBP1, HSPA5 and SEC61A1 expression increased with fibrosis. Tertiary lymphoid structures retained TCF7-high stem-like T cells that received CCL19–CCR7 signals from fibroblasts. Using multimodal deep-learning models to delineate anatomical regions, we separated Banff lesions into local immune-load and chronic morphological-injury axes, each with distinct compartmental responses. Peritubular-capillary endothelium increased interferon-γ-associated antigen presentation with local immune load, glomerular endothelium reduced MHC expression with glomerular inflammation, and chronic injury was accompanied by reduced metabolic programmes in specific cell types. In mice, we built a single-cell atlas of kidney transplantation covering syngeneic grafts and BALB/c, DBA/2J and DBA/2N→C57BL/6 models. Graft trajectories separated into rejection-associated and tolerance-associated paths that differed in interferon-activated myeloid states and *Hnf4a*– and *Ppara*-associated tubular metabolism. In 20 human recipients with rejection-spectrum diagnoses, predictions of functional non-response from a hierarchical causal-inference model were most sensitive to plasma-cell niche occupancy and density. Niche maps predicted from routine H&E were associated with subsequent kidney function. These findings show that kidney allograft rejection is accompanied by changes in cellular states and spatial organisation, and identify plasma-cell niches in residual and scarred interstitium as tissue features of functional non-recovery after rejection.

## Introduction

Kidney transplantation is the preferred treatment for kidney failure, yet rejection, BK polyomavirus nephropathy and calcineurin-inhibitor toxicity continue to limit graft survival. The Banff classification grades rejection by the location of inflammation and the extent of chronic damage^1–3^. Recipients who share a Banff diagnosis nevertheless follow different courses. Chronicity predicts graft loss after antibody-mediated rejection^4^, inflammation within scarred interstitium (i-IFTA) predicts poor graft outcome^5,6^, and microvascular inflammation carries risk even in the absence of donor-specific antibodies^7^. Banff scores grade how much inflammation and scarring each structure contains; the cells that occupy these structures, and their relationships with the surrounding stroma and parenchyma, are largely undescribed.

Single-cell studies of allograft biopsies have catalogued immune cell states, donor–recipient chimerism and the metabolic shutdown of structural cells during rejection^8–10^. Spatial transcriptomics has placed these responses in glomeruli, injured tubules, the interstitium and immune–fibrotic regions^11–14^, and a spatial atlas of diabetic kidney disease linked local immune environments to disease progression^15^. Imaging that measures RNA and protein at subcellular resolution^16^ allows cell identity, anatomical position and lesion grade to be read in the same biopsy section. Systematic links between in situ transcriptional signals and the structural units affected by Banff lesions, together with changes in cellular states within these structures, remain limited.

Changes in cellular states within kidney allografts are closely associated with outcomes. Plasma-cell-rich rejection responds poorly to treatment and shortens graft survival^17–19^, and CD38-directed antibodies, which target plasma cells and natural killer cells, have entered clinical study in antibody-mediated rejection^20–22^. In tubules, part of the proximal tubule population remains in a VCAM1-positive failed-repair state after acute kidney injury and sustains inflammatory programmes^23–25^. T-cell-mediated rejection in mice induces related injured epithelial states, and their abundance in human allografts correlates with graft survival^26^. Whether plasma cells occupy defined tissue environments, and whether such environments relate to functional recovery after treatment, requires further study.

Single-cell analysis of rejecting mouse kidney allografts identified a myeloid differentiation pathway associated with rejection^27^. DBA/2→C57BL/6 kidney allografts, by contrast, are accepted spontaneously, retain regulated intragraft T-cell responses and form Treg-rich organized lymphoid structures^28–30^. The key drivers of immune and parenchymal states in these grafts, and how these states relate to human rejection, remain unclear.

In this study, we profiled 83 human kidney allograft biopsies by subcellular RNA and protein imaging with registered histology, and linked them through one cell hierarchy to human and mouse single-cell data. Together these data form SPECTRA (Spatial Etiology and Coordination of Transplant Rejection in Allografts), a name that echoes the Banff 2024 view of rejection as a spectrum of phenotypes^3^. We defined nine tissue niches and asked which of them related to functional recovery, how tertiary lymphoid structures were organized, and how each compartment scored by Banff responded to acute and chronic lesions. We then followed mouse grafts along rejection-associated and tolerance-associated trajectories, related the plasma-cell niche to post-treatment function in a multilevel model, and translated niche maps to routine H&E sections.

## Results

### A cross-species spatial atlas of kidney allografts at single-cell resolution

We profiled 84 tissue cores from 83 human allograft biopsies by CosMx spatial molecular imaging, which measured 18,934 RNA targets and 64 protein features in 1,011,984 cells, and registered each section with H&E and multiplex immunofluorescence (Fig. 1a). The human spatial cohort comprised 20 biopsies with mixed rejection, 18 with antibody-mediated rejection or microvascular inflammation (AMR/MVI) and 14 with T cell-mediated rejection (TCMR). Other biopsies showed BK polyomavirus nephropathy (ten), borderline changes (nine), calcineurin-inhibitor toxicity (four) or C4d positivity without rejection (one), and seven were controls. Controls included six day-zero biopsies and one post-transplant reference biopsy from a graft with stable function. We linked the spatial data to a human single-cell reference of 352,911 cells and to 442,320 cells from 34 mouse kidney grafts. The human– mouse reference embedding in Fig. 1b displays these two single-cell datasets; the separate CosMx embedding is related through the same cell hierarchy, which resolves 26 major spatial cell identities (Fig. 1b). Spatial annotation combined eight RNA reference-transfer methods with protein-assisted hierarchical calls and anatomical masks, and divided the 26 major identities into 65 second-level and 114 third-level labels (Extended Data Fig. 1a). After integration, cells from the two CosMx slides, 97 human single-cell libraries and individual mice intermixed within their embeddings (Extended Data Fig. 1b). The mouse cohort comprised syngeneic grafts on postoperative day 7, untreated BALB/c→C57BL/6 allografts on days 7, 28 and 90, and tacrolimus-treated BALB/c grafts on day 7. It also included presensitised BALB/c and DBA/2N grafts on days 7 and 28 and DBA/2J grafts on days 7, 28 and 90.

**Fig. 1.**
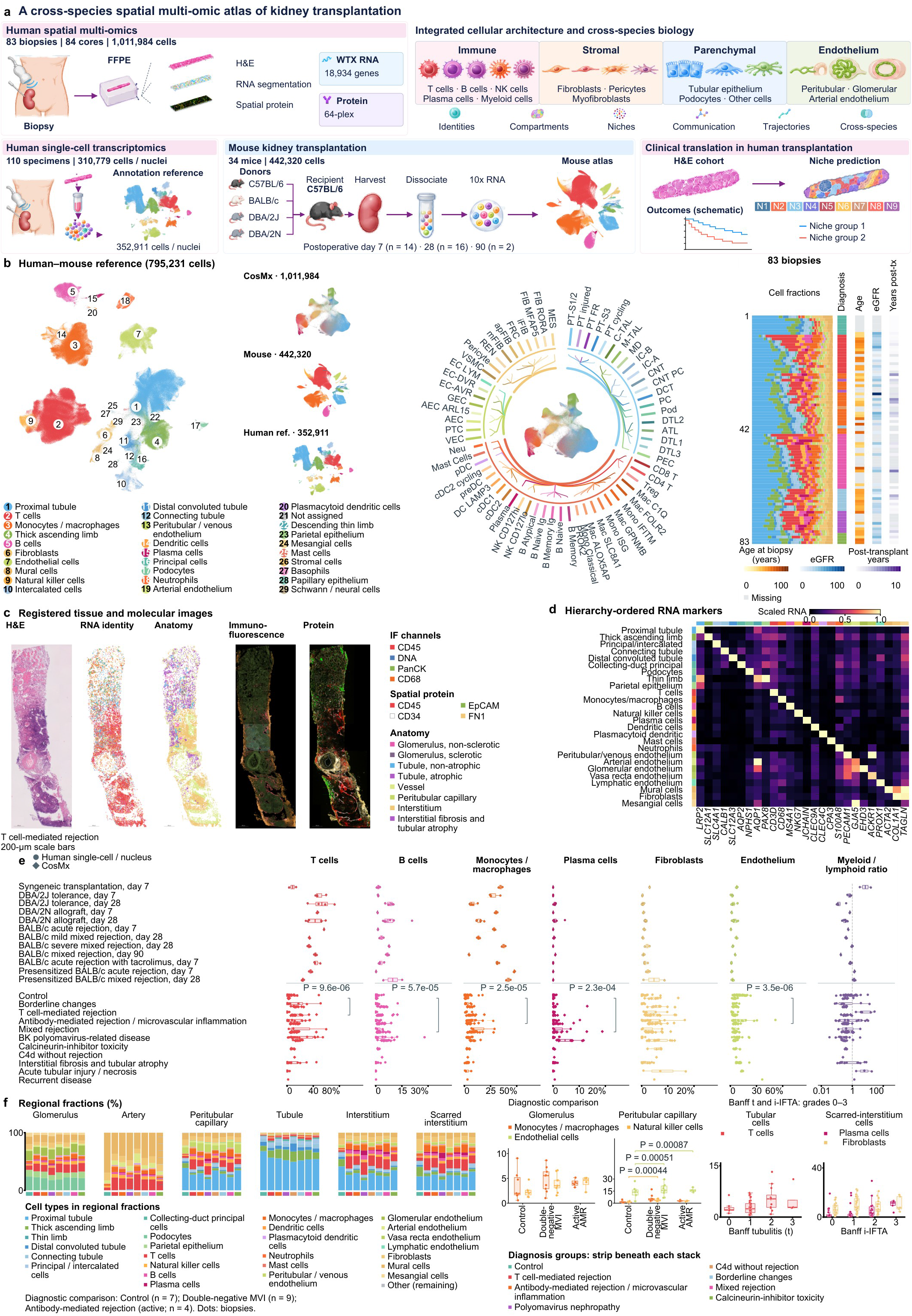
| A spatial and single-cell atlas of human and mouse kidney allografts. **a**, Study design linking the human spatial cohort (83 biopsies, 84 cores and 1,011,984 cells), human whole-tissue single-cell data (110 specimens and 310,779 cells or nuclei) and mouse kidney transplantation (34 animals and 442,320 cells). The human UMAP shows the 352,911-cell annotation reference; the mouse UMAP shows the measured mouse atlas. Spatial RNA and protein measurements are registered with histology and immunofluorescence. Survival curves illustrate the clinical translation concept. **b**, Human–mouse single-cell reference embedding (795,231 cells: 352,911 human reference cells and 442,320 cells from 34 mice), the CosMx cell hierarchy, and the separate CosMx ResolVI UMAP, mouse and human reference embeddings. Numbered identities and cell colours are keyed by name. The human–mouse embedding was fitted without public mouse data and displays only the mouse cohort of this study. The clinical sidebar shows stacked cell fractions and diagnosis, estimated age at biopsy, eGFR at biopsy and time after transplantation for all 83 spatial biopsies in a common order. Adjacent orange-brown, blue and purple scales encode age (0–100 years), eGFR (0–100 mL/min/1.73 m²) and time after transplantation (0–10 years), respectively; grey denotes missing measurements. Median [25th–75th percentiles] values are 44.3 [36.5–52.8] years (n = 47), 30 [22.2–36.8] (n = 46) and 0.5 [0.1–2.0] years (n = 47), respectively. Six implantation biopsies are placed at time zero. Diagnostic colours are defined in f. **c**, Registered H&E, RNA level-1 identities, anatomy, multiplex immunofluorescence and protein views of a representative biopsy, with channel and anatomy keys. Scale bars, 200 μm. **d**, First marker gene of each of the 26 level-1 identities in hierarchy order; rows are cell types (colour strip, identity colour), columns marker genes, and colour shows gene-wise 0–1 scaling. **e**, Fractions of T cells, B cells, monocytes/macrophages, plasma cells, fibroblasts and endothelial cells, and the myeloid-to-lymphoid ratio. The myeloid numerator includes monocytes/macrophages, dendritic cells, plasmacytoid dendritic cells, neutrophils and mast cells; the denominator includes T, natural killer, B and plasma cells. Mouse fractions use the corresponding available myeloid identities, including basophils. Ratios use no pseudocount and a logarithmic axis. Each point represents one mouse, one of 94 human sampling units or one of 83 CosMx biopsies. Human single-cell and single-nucleus units share diagnosis boxes and circles; diamonds denote CosMx. Human Control contains 13 post-transplant references and 22 native-kidney units; CosMx Control contains six designated implantation biopsies and one post-transplant reference. Human single-cell distributions are shown by sampling unit; group inference uses CosMx biopsies. Brackets give unadjusted P values from year/depth-adjusted CosMx models only for comparisons with complete-family BH q < 0.05; complete effect estimates, P values and BH q values are in Source Data. Boxes show the median and interquartile range with 1.5-IQR whiskers. One human ratio is undefined because its lymphoid denominator is zero; its other fractions are shown. **f**, Biopsy-mean cellular composition by diagnosis in glomeruli, arteries, peritubular capillaries, tubules, interstitium and scarred interstitium. Strips beneath the stacks identify diagnosis; local keys define 26 level-1 identities plus remaining cells and eight diagnostic groups. Control combines six implantation biopsies and one post-transplant reference. Boxplots compare glomerular monocyte/macrophage and natural killer fractions and peritubular-capillary monocyte/macrophage, natural killer and endothelial fractions among controls (seven biopsies), double-negative MVI (nine biopsies) and active AMR (four biopsies). Double-negative MVI requires a diagnosis of MVI with negative donor-specific antibodies and C4d; MVI with TCMR or borderline changes, BK polyomavirus nephropathy and probable AMR are excluded from this three-group comparison. Tubular T-cell and scarred-interstitial plasma-cell and fibroblast fractions are shown by Banff t and i-IFTA grades 0–3. Peritubular-capillary endothelial fractions sum the EC-Art, EC-LYM, EC– PTC_VEC and EC-VR identities. Boxplot colours identify cell types, and denominators contain all cells in the corresponding biopsy compartment. Boxes show medians and interquartile ranges, whiskers extend to the most distant observation within 1.5 interquartile ranges, and dots represent biopsies. Three-group models adjust for sampling year and capture depth; BH correction covers three pairwise contrasts across 44 compartment–cell fractions (132 tests). Brackets show raw P values for comparisons with complete-family q < 0.05. All effects, P values and q values are in Source Data.

Registered H&E, RNA identities, anatomical masks and protein images place each cell in the structure it occupies (Fig. 1c). Lineage markers such as LRP2, CD3D, COL1A1, GJA5 and EHD3 distinguish proximal tubule, lymphocyte, fibroblast and endothelial identities (Fig. 1d). We aligned patient diagnoses, age at biopsy, eGFR at biopsy and time after transplantation with cell composition, showing wide variation in immune, epithelial and stromal content among biopsies that share a diagnosis (Fig. 1b). In sample-level RNA principal components, human samples with the same diagnosis did not form separate groups (Extended Data Fig. 1c).

Biopsies with T-cell-mediated rejection (TCMR), antibody-mediated rejection or microvascular inflammation (AMR/MVI) and mixed rejection had higher median proportions of T cells, B cells, monocytes/macrophages and plasma cells than reference biopsies. In human single-cell data, AMR/MVI also carried more monocytes/macrophages (Fig. 1e). The myeloid-to-lymphoid ratio rose from borderline changes (median 0.30) and TCMR (0.38) to mixed rejection (0.48). Rejection severity also varied in mice: the ratio was 0.97 in severe and 0.40 in mild mixed rejection on day 28 and 1.34–1.39 in presensitised grafts. Severe mixed rejection also had fewer endothelial cells than mild rejection, and DBA/2J grafts retained abundant T cells on both days 7 and 28 (Fig. 1e). Plasma cells accumulated with time after transplantation. In BALB/c grafts they rose from 0.03% of cells on day 7 to 0.87% on day 90. In human single-cell data they were most frequent in interstitial fibrosis and tubular atrophy (IFTA) and in mixed rejection (0.35% and 0.41%; Fig. 1e).

Anatomical position resolved where these infiltrates accumulate. The median tubular T-cell fraction rose from 2.1% of tubular-compartment cells at Banff t0 to 5.3% at t2 and was 2.8% in the four biopsies at t3. The median plasma-cell fraction of scarred residual interstitium rose from 0.9% at i-IFTA 0 to 3.3% at grade 2 and 11.5% in the five biopsies at grade 3 (Fig. 1f). Peritubular-capillary composition also changed with microvascular disease. After adjustment for sampling year and capture depth, endothelial fractions were higher than in controls in microvascular inflammation negative for donor-specific antibodies and C4d (double-negative MVI; q = 0.0132) and in active AMR (q = 0.0144). Double-negative MVI also had a higher natural killer-cell fraction (q = 0.0132), and its monocyte/macrophage fraction showed an upward trend (q = 0.070; Fig. 1f). Because immune cells differed in abundance from one structure to the next, we next asked whether immune and tissue cells assemble into recurrent local environments that cross anatomical boundaries.

### Spatial niches in kidney allografts

We used CellCharter^31^, an unsupervised method to partition cell neighbourhoods. It concatenates the expression latent variables of each cell with the aggregated latent variables of its first– to third-order spatial neighbours and clusters the result with a Gaussian mixture model. Regions with similar cell composition and expression context are therefore grouped together across samples. After comparing cluster stability and correspondence with pathological structures for 3 to 15 clusters, we identified nine niches that recurred across samples (Fig. 2a and Extended Data Fig. 2a). These niches matched known kidney structures. Four were dominated by tubular epithelium: proximal tubular parenchyma (N3), a mixed proximal-tubule niche (N4), distal nephron and collecting duct (N6), and thick ascending limb and medulla (N7). The other five were a glomerular and vascular scaffold (N1), a T-cell and B-cell lymphoid environment (N2), tubulointerstitial remodelling (N5), a vascular, stromal and immune interface (N8) and a plasma-cell effector environment (N9). B and T cells dominate N2, fibroblasts and myeloid cells gather in N5, and plasma cells concentrate in N9. N4 combines a predominant proximal-tubule population with T cells, mural cells and monocytes/macrophages (Fig. 2b).

**Fig. 2.**
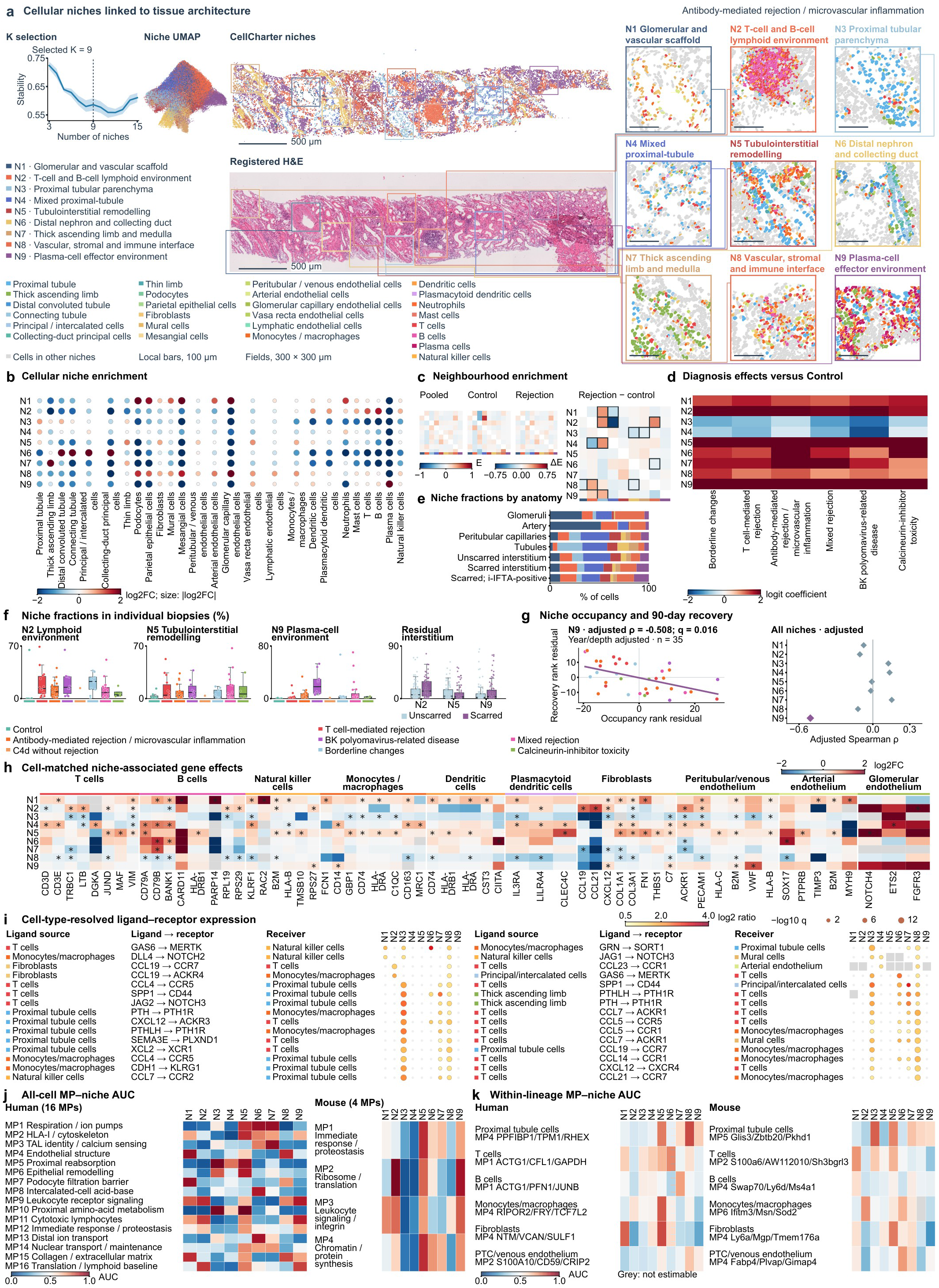
| Tissue niches and cell-intrinsic programmes. **a**, Cell coordinates and level-1 segmentation of a representative biopsy, with the CellCharter stability curve and UMAP for the nine-niche solution. Scale bars show physical distances. **b**, Enrichment of 26 cell classes across nine niches. Colour indicates log2 fold change; dot area scales with its magnitude. **c**, Pooled, Control, rejection and rejection-minus-Control neighbourhood enrichment. E is observed-minus-expected adjacency and ΔE the group difference. Outlines mark directed pairs with permutation BH q<0.05. Permutation testing used 100 sample-label permutations; BH correction covers 72 directed pairs across three contrasts. **d**, Year– and depth-adjusted logit-limma coefficients for six diagnoses relative to Control. Colours saturate at −2 and +2; complete coefficients, model P values and BH q values across nine niches are in Source Data. **e**, Biopsy-weighted mean niche fractions by anatomical compartment. The last bar represents scarred interstitium in 54 biopsies with Banff i-IFTA>0. **f**, Niche fractions in 83 biopsies and in unscarred and scarred interstitium from 82 and 81 biopsies, respectively. Points denote biopsies. Boxes span the 25th–75th percentiles, centre lines are medians, and whiskers reach the most distant observation within 1.5 interquartile ranges. The key below the fourth subpanel identifies unscarred and scarred regions. **g**, Adjusted association between N9 grid occupancy and relative eGFR change approximately 90 days after biopsy. Points are year– and sequencing-depth-adjusted rank residuals for 35 patients; the line is a descriptive least-squares fit in these coordinates. Spearman ρ = −0.508123, P = 0.00183134 and BH q = 0.0164821 across nine niches. The adjacent plot shows adjusted correlations for all nine niches. **h**, Sixty niche-associated genes in ten cell classes: T cells, B cells, natural killer cells, monocytes/macrophages, dendritic cells, plasmacytoid dendritic cells, fibroblasts, and peritubular/venous, arterial and glomerular endothelium. Colour shows the smiDE/metafor log2 fold change between each niche and the remaining niches, saturating at ±2. Stars indicate BH q<0.05; grey denotes missing estimates. Each displayed gene has at least one positive, q<0.05 niche contrast within its cell class. **i**, Thirty cell-type-specific ligand–receptor expression associations. Each row identifies the ligand-expressing cell, ligand, receptor and receptor-expressing cell; at least one partner is an immune cell. Colour shows the mean paired log2 ratio of expression products between a niche and the remainder of the same core. Dot area increases with −log10 BH q and is capped at 12. Coloured dots require q<0.05, a mean log2 ratio≥0.5 and at least 12 paired cores; colour saturates at 4. Pale dots denote tested combinations outside these display criteria, and grey squares denote unavailable comparisons. Ligand–receptor orientation agrees with CellChatDB. **j**, All 16 human and four mouse all-cell metaprogrammes compared with nine human CosMx niches. AUC is the probability that a programme member has a higher niche identity score than a background gene, counting ties as one half. Each niche uses all 18,918 ranked genes. Scores derive from depth-adjusted niche pseudobulk edgeR specificity, with mouse genes mapped to one-to-one human orthologues. BH correction covers 144 human and 36 mouse programme–niche comparisons separately. **k**, Cell-matched human and mouse programme–niche stratified AUC for proximal tubule, T, B, monocyte/macrophage, fibroblast and peritubular/venous endothelial cells. For each class, the programme with the largest AUC in any tested niche is held fixed across all nine niches. Names summarize programme gene content; grey denotes missing estimates. Complete matrices, names and cell-subtype correspondences are provided in Extended Data Fig. 2d–g and its Source Data atlas.

Rejection rearranged niche abundance and adjacency. N2 was less often adjacent to N4 in rejection than in Control (ΔE = −0.71) and more often adjacent to N8 (ΔE = +0.36; permutation q = 0.046 for both; Fig. 2c). After adjustment for sampling year and sequencing depth, N2, N5 and N9 were all more abundant in mixed rejection than in Control (q ≤ 0.024; Fig. 2d and Extended Data Fig. 2b). Niche fractions for the 15 recorded diagnoses within rejection, resolved by Banff grade and coexisting lesions, are shown in Extended Data Fig. 2c. N9 extended into the residual interstitium in which i-IFTA is graded. It occupied a mean of 8.8% of unscarred and 12.4% of scarred residual interstitium, and 15.0% of scarred interstitium in the 54 biopsies with i-IFTA > 0 (Fig. 2e,f).

We then asked whether any niche related to recovery of kidney function. Among 35 patients with complete follow-up records, greater N9 occupancy was associated with a smaller relative improvement in eGFR about 90 days after biopsy (Spearman ρ = −0.508, q = 0.0165; Fig. 2g). This association was adjusted for sampling year and sequencing depth.

Cells of the same class adopted different states in different niches (Fig. 2h). In N2, fibroblasts expressed more CCL19, CCL21 and CXCL12, peritubular capillary and venular endothelium more ACKR1 and T cells more LTB. In N5, monocytes/macrophages expressed more GBP1, CD74, HLA-DRA and C1QC, and fibroblasts more COL1A1, COL3A1, THBS1, FN1 and C7. In N9, fibroblasts expressed more CXCL12 and C7, monocytes/macrophages more CD14, and peritubular capillary and venous endothelium more VWF. CXCL12 supports plasma-cell residence and survival in bone marrow^32,33^, so its stromal expression in N9 suggests a candidate retention signal for plasma cells in the graft. Communication by the same cell class also differed between niches. Among cell-type-specific ligand–receptor expression products oriented by CellChatDB^34^, fibroblast CCL19 paired with T-cell CCR7 (q = 2.8 × 10⁻⁸) and with macrophage ACKR4 more strongly in N2 than in the rest of the same cores (Fig. 2i). This points to a distinct stroma–immune interaction in N2.

We next asked whether these niche-specific cellular programmes could be identified in mouse and human single-cell data. The 366 sample-derived NMF programmes from 118 human libraries and the 120 mouse programmes formed 16 and 4 all-cell metaprogrammes, respectively (Extended Data Fig. 2d). To relate programmes to niches, we ranked 18,918 genes in each niche by pseudobulk specificity relative to the other niches and computed the rank AUC of the member genes of each metaprogramme in this ranking. An AUC of 0.5 indicates randomly distributed member genes, and values near 1 indicate member genes concentrated among the most niche-specific genes. Among all-cell metaprogrammes, the human leukocyte receptor-signalling programme and the mouse leukocyte signalling and integrin programme both matched N9 most closely (AUC = 0.90 and 0.80; Fig. 2j and Extended Data Fig. 2e). N9 therefore carries immune signalling features beyond those of plasma cells. Within cell classes, human and mouse fibroblast matrix programmes both matched N5. Proximal-tubule programmes matched different niches in the two species: the human PPFIBP1/TPM1/RHEX programme matched N8, and the mouse Glis3/Zbtb20 programme matched N3 (Fig. 2k and Extended Data Fig. 2f). Assigned to their highest-AUC niches, the 79 human and 64 mouse within-lineage metaprogrammes spanned all nine niches (Extended Data Fig. 2g). Because N2 joined B cells and T cells with stromal chemokines, which provide the conditions for tertiary lymphoid structure formation, we next examined the cell states of these structures in N2.

### Stem-like T cells and immune–stromal interaction programmes in tertiary lymphoid structures

We identified 167 tertiary lymphoid structures (TLS) in 40 cores by lymphocyte-density clustering and pathologists’ annotation and classified 102 as Stage I and 65 as Stage II by depth-adjusted CR2 and FCER2 expression (Fig. 3a). In the 17 cores that contained both stages, follicular dendritic cell scores were higher in Stage II structures (Fig. 3a and Extended Data Fig. 3a). A homing programme was higher inside both stages than in their matched exteriors (Stage I, q = 0.0037; Stage II, q = 0.014; Fig. 3a). Stage II structures predominated only in AMR/MVI cores. Most structures lay beside vessels or tubules, and veins accounted for 64 of the 70 vessel-adjacent structures (Extended Data Fig. 3i).

**Fig. 3.**
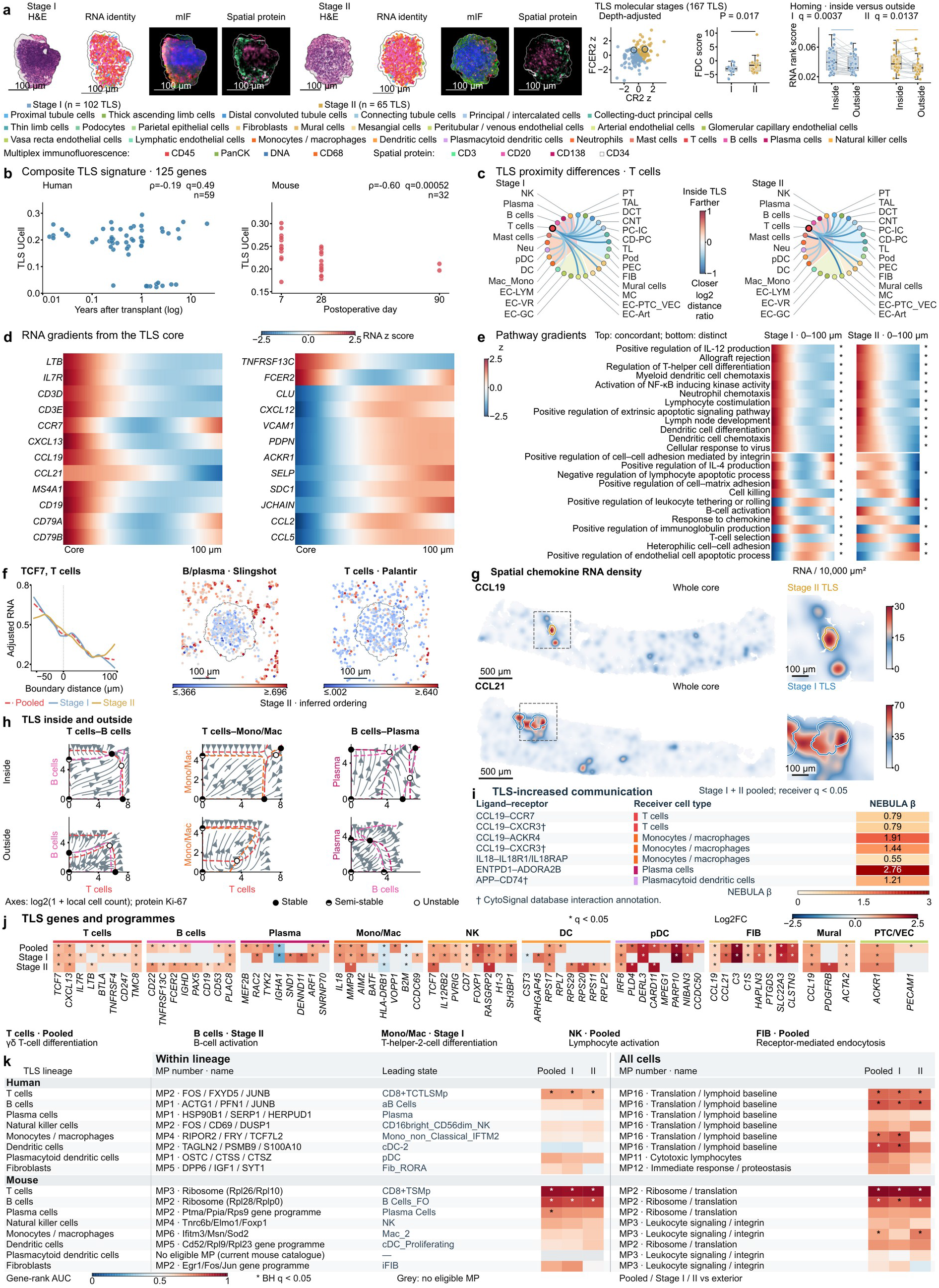
| Spatial organization and cellular states of tertiary lymphoid structures. **a**, Registered H&E, RNA cell identities, multiplex immunofluorescence and spatial protein measurements of representative Stage I and Stage II lymphoid aggregates. Scale bars, 100 µm. The scatter displays 167 aggregates classified from depth-adjusted CR2 and FCER2 signals (Stage I, 102; Stage II, 65). FDC scores are shown for the 17 cores containing both stages; each point is a stage-specific core mean (two-sided paired Wilcoxon P = 0.01743; BH q = 0.05228 across the FDC, germinal-centre and high-endothelial-venule scores). The Homing programme is compared inside TLS and in its protected exterior, separately for Stage I (33 paired cores; P = 0.0016955372; q = 0.003673664) and Stage II (24 paired cores; P = 0.0079205036; q = 0.013728873). Lines join paired core means. Homing q values are corrected across all 13 programmes and both stages (26 tests). **b**, Composite TLS-signature scores in relation to transplant time. The signature contains 125 genes with positive optimal/metafor effects and FDR < 0.05 in at least one level-1 lineage; 119 one-to-one orthologues were scored in mice. Each point is the mean pyUCell score across all cells from one human tissue sample or mouse animal. Human data comprise 59 whole-tissue post-transplant samples with recorded transplant intervals, shown in years on a logarithmic axis. Mouse data comprise individual animals at their recorded postoperative days. Spearman ρ and BH q values are shown; correction covers all cells and eight lineages across three TLS signatures within each species. Lineage-specific, diagnostic and mouse– model summaries are provided in Source Data. **c**, Stage-specific differences in BallTree proximity to T cells. Lineages are arranged on a circle and joined by Bézier curves. Only paired inside-versus-outside comparisons with BH q < 0.05 are drawn. Edge colour is the median log2 ratio of density-normalized partner-to-hub distance inside versus outside TLS; blue indicates closer proximity inside. Node colours identify level-1 lineages and the filled sectors identify broad lineages; cell-type abbreviations follow the full-name key in a. All five hub classes and the complete test families are shown in Source Data. **d**, Twenty-four transcript gradients from the TLS core to approximately 100 µm, expressed as within-gene z scores of the depth– and detection-adjusted fitted values. CCL19 has a significant distance association (q = 1.96 × 10⁻¹⁸); the adjacent CCL21 profile has q = 0.106. The other displayed genes have q < 0.05. **e**, Twenty-four pathway-score profiles across Stage I and Stage II aggregates. The upper block contains 12 concordant profiles with significant distance associations in both stages. The lower block contains 12 profiles with lower correspondence between the fitted stage curves; each pathway is significant in at least one stage. Asterisks mark within-stage BH q < 0.05. Curves are standardized within each pathway and stage. Gene and pathway models account for RNA depth and detection and use aggregate-clustered standard errors. Complete feature families, slope estimates, P and q values accompany the figure. **f**, T-cell TCF7 expression across the TLS boundary. Negative distances lie inside TLS. Curves show pooled, Stage I and Stage II profiles; the pooled data contain 49 TLS, 27 cores and 11,998 T cells. Shape-preserving cubic interpolation connects 16 LOWESS estimates per profile (LOWESS span 0.4). The plotting area is square. **Spatial inferred ordering in f.** Slingshot ordering for the joint B/plasma-cell population and Palantir ordering for T cells are mapped to cell coordinates in the same Stage II TLS. The dashed line marks the TLS boundary. Both panels use equal coordinate scaling and a 100– µm scale bar. Slingshot colours span 0.366–0.696 on a linear scale; Palantir colours span 0.002–0.640 on a logarithmic scale, with values beyond these limits assigned the endpoint colours. Scores describe relative transcriptional ordering within the input populations. Radial means and smoothed curves are provided in Source Data. **g**, Whole-core CCL19 and CCL21 RNA densities in two representative cores, with 570 × 570 µm TLS-centred views. Representative cores were selected from 27 cores containing 49 TLS by requiring the whole-core density maximum of the corresponding gene to lie within an annotated TLS. All 12,649 and 18,634 measured cells, respectively, were included. Maps show transcript density per 10,000 µm² (5-µm grid; Gaussian σ = 35 µm; kernel truncated at 3σ). Smoothed counts were divided by 25 µm² and multiplied by 10,000. Whole-core and zoom views for each gene share the same unclipped linear colour scale. Maps are displayed within 25 µm of measured cells. Gold and blue outlines mark Stage II and Stage I TLS boundaries; dashed boxes indicate the zoom regions. Scale bars, 500 µm for whole cores and 100 µm for zoom views. Both cores and their TLS outlines were rigidly rotated by 90°. The two genes are shown in different samples. Density maps and per-cell means for all candidates are provided in Source Data. **h, Pairwise population dynamics inside and outside TLS.** Columns use local densities of T and B cells, T cells and monocytes/macrophages, or B and plasma cells as the OSDR model axes. Rows represent the interior and exterior of the same pooled TLS definition. Models use protein Ki-67 division labels from 40 tissue cores. The B–plasma-cell models include 11,103 B cells and 1,179 plasma cells inside, and 3,601 and 1,126 outside. Axes show log2(1 + cells within 80 µm), and arrows show fitted changes in these log coordinates. Both compartments share display limits within each pair; vector fields are restricted to each model’s observed 99.5th-percentile local-density range. Dashed lines are cell-type-coloured zero-change contours. Filled, half-filled and open points denote stable, semi-stable and unstable fixed points, respectively. Stage-specific and RNA MKI67 models are provided in Source Data. **i, Increased receiver signalling inside TLS.** Colours and numbers show all seven positive receiver-level CytoSignal NEBULA β coefficients with BH q < 0.05 for TLS interiors versus exterior tissue, pooling Stage I and Stage II. The six ligand–receptor pairs span T cells, monocytes/macrophages, plasma cells and plasmacytoid dendritic cells. Coloured strips identify the receiver cell type. The models used tissue core as the random effect and total RNA counts as the offset. BH correction was applied separately within each receiver, diffusion/contact score class and fitted covariate contrast. The displayed receiver fits include 40, 39, 24 and 30 cores, respectively; cell counts and P and q values are provided in Source Data. †CCL19–CXCR3 and APP–CD74 follow the CytoSignal database interaction annotations. **j, Cell-matched TLS gene programmes.** Optimal-smiDE/metafor estimates are shown for 69 class–gene pairs across ten cell classes in Pooled, Stage I and Stage II comparisons. Colour denotes log2 fold change; a single asterisk denotes adjusted q < 0.05. Grey denotes a class–gene comparison without an available fitted estimate. Biological examples with q < 0.05 in at least one comparison were prioritised, followed by non-ribosomal genes ranked by minimum q and then the remaining genes in the same ranking. LEF1 was excluded from the display. Positive-list GO biological-process terms are labelled by their comparison; cell classes without significant enrichment have no GO label. Full effects, uncertainty estimates, q values and GO terms are provided in Source Data. **k**, Correspondence between lineage-resolved TLS gene effects and human or mouse metaprogrammes (MPs). Within lineage and All cells denote the scope in which MPs were derived. Both displays compare MPs with the TLS effect vector for the named lineage. Human and mouse are shown separately; each contains T cells, B cells, plasma cells, natural killer cells, monocytes/macrophages, dendritic cells, plasmacytoid dendritic cells and fibroblasts. For each species, lineage and catalogue scope, the MP with the highest pooled TLS-versus-exterior AUC is shown for pooled TLS, Stage I and Stage II versus exterior. Labels give the MP number and its catalogue name. Within-lineage MP labels also give the highest-scoring cell state. The leading cell state for human T-cell MP2 is labelled CD8+TCTLSMp. Colour denotes competitive gene-rank AUC, with 0.5 indicating no rank preference; asterisks denote BH q < 0.05 across the complete MP-by-cell-type test family within species and contrast. Grey denotes an unestimated comparison. The mouse within-pDC comparison was unestimated. AUCs, P and q values, gene membership and the complete test family are provided in Source Data.

TLS had a layered internal organization. After adjustment for local cell spacing, T cells and B cells were closer to each other inside TLS, and dendritic cells, monocytes/macrophages and fibroblasts showed stage-specific proximity patterns (Fig. 3c). CCL19, LTB, IL7R, MS4A1 and TNFRSF13C formed an expression gradient that declined from the TLS core towards the periphery. CXCL12, VCAM1 and several secretion-associated transcripts instead rose towards the exterior (Fig. 3d and Extended Data Fig. 3b–d,j). Both stages shared core-enriched programmes for lymphocyte costimulation, helper T-cell differentiation and dendritic-cell chemotaxis, and differed in the spatial profiles of B-cell activation, immunoglobulin production and adhesion (Fig. 3e). In T cells, the stemness– and homing-associated genes TCF7 and CCR7 declined from the TLS interior towards the exterior. IL7R, SELL and MAL changed in the same direction with smaller magnitude (Fig. 3f and Extended Data Fig. 3e,f). We further ordered T cells, B cells and plasma cells in pseudotime. Pseudotime of B and plasma cells (Slingshot^35^) rose slightly from the core outwards, suggesting a possible inside-out maturation gradient, whereas pseudotime of T cells (Palantir^36^) showed a weaker gradient in the same direction (Fig. 3f). Cell proportions supported this organization: CD4+ and CD8+ TSN T cells and naive B cells were most frequent at the TLS centre, and fibroblastic reticular cells peaked about 25 µm from it (Extended Data Fig. 3g). Hotspots of lymphoid-organizer chemokines such as CCL19 and CCL21 also lay within TLS (Fig. 3g).

The CCL19 gradient led us to test whether the corresponding receptor signals were stronger inside TLS. Pooling both stages, seven receiver-level CytoSignal coefficients were higher inside TLS than outside, including CCL19–CCR7 reception by T cells (NEBULA^37^ β = 0.789, q = 8.05 × 10⁻⁶; Fig. 3i). Expression measured at both molecular ends supported this result. Across paired cores, the product of fibroblast CCL19 and T-cell CCR7 was higher inside TLS (q = 2.68 × 10⁻⁸). Forty-five concordant source–target relationships from 11 source classes supported five of the seven receiver effects (Extended Data Fig. 3h). Within cell classes, T cells of both stages expressed more TCF7 and CXCL13 inside TLS, Stage II B cells more CD22 and TNFRSF13C, and Stage I fibroblasts more CCL21 and C3 (Fig. 3j). These results suggest one possible organization of TLS in kidney allografts, in which activated fibroblasts produce chemokines that recruit T and B cells and maintain a local environment where TCF7-high T cells coexist with B cells.

We next modelled how pairs of populations change with local density, using protein Ki-67 division labels in OSDR. Inside TLS, the T–B, T–monocyte/macrophage and B–plasma-cell models all predicted stable coexistence. Outside TLS, the T–B model instead had two stable states, each dominated by one cell type, and the B–plasma-cell model again predicted coexistence (Fig. 3h and Extended Data Fig. 3n). Stage-resolved B–plasma-cell models predicted coexistence in Stage I interiors and exteriors and bistability in Stage II. These dynamics suggest that TLS, through organized stroma and chemokine signals, may keep lymphocytes in balance across a wide range of densities. Outside TLS, T and B cells may compete for local space or survival signals, so that local tissue settles into a state dominated by one cell type. The B–plasma-cell bistability in Stage II suggests that mature TLS may switch between B-cell-dominated and plasma-cell-output states. OSDR estimates how the balance of proliferation and clearance varies with local cell density. To examine how TLS states change, we aggregated the cell composition of each structure and ordered the 167 structures accordingly. The order extended from B-cell-rich aggregates towards aggregates with more monocytes/macrophages, natural killer cells and dendritic cells. Along this order, FOXP1, KLF13, IRF8, TCF7, NFATC1 and KLF2 declined and MEF2B rose (Extended Data Fig. 3k).

To place TLS states in time and across species, we scored a composite signature of 125 genes that were upregulated inside TLS in at least one cell class. Its all-cell score declined with postoperative day in mice (ρ = −0.602, q = 0.000517). In human samples it showed no association with time after transplantation (ρ = −0.19, q = 0.49, n = 59; Fig. 3b). In BALB/c grafts, CD8+ TSMp stem-like memory T cells declined after day 7, whereas CD4+ TH17DA cells increased through day 90 (Extended Data Fig. 3m). The CCL19–CCR7 expression product in single-cell data was higher in human AMR/MVI than in Control and rose from day 7 to day 28 in DBA/2J and DBA/2N grafts. In BALB/c grafts it was lower on day 28 and rose again on day 90 (Extended Data Fig. 3l). Metaprogramme correspondence linked TLS T cells to a FOS/FXYD5/JUNB programme led by CD8+ TCTLSMp cells in humans and to a ribosomal programme in mice (AUC = 0.7125 and 0.9386; Fig. 3k). Accepted mouse kidney grafts form organized regulatory lymphoid structures^28^, yet the single-cell score declined with time after transplantation. TLS transcriptional features may therefore be diluted by signals from other cells in dissociated whole-tissue single-cell data, which underlines the need for spatial analysis. Most graft-infiltrating immune cells lie outside such structures, in the compartments that Banff scores for acute rejection, and we next asked how each compartment responds to the immune burden it carries.

### The landscape of acute lesions reveals compartment-specific responses

To measure lesions in individual structural units, pathologists first manually annotated glomeruli, tubules, arteries, peritubular capillaries and interstitium. We trained in-house deep-learning segmentation models on these annotations and ran inference on all fields, and pathologists then reviewed every segmentation mask in QuPath^38^. Because each cell carries a position and identity at single-cell resolution, we could assign cells to the structural units they occupy and measure local inflammatory burden separately in the five compartments scored by Banff (Fig. 4a). Unlike semi-quantitative Banff scores, these standardized quantitative measures describe the local inflammatory burden of each structure. Interstitial burden counts mononuclear cells per residual interstitial area, and tubulitis burden counts mononuclear cells per ten tubular epithelial cells in the wall. Arterial burden counts mononuclear cells per intimal area, glomerular burden immune cells per glomerulus and capillary burden immune cells per peritubular capillary area. Across burden strata, cell composition within lesioned structures showed an overall increase in immune cells and loss of parenchymal cells. T-cell and monocyte/macrophage fractions rose with burden in all five compartments. Proximal tubule cells declined across tubulitis tiers, and peritubular endothelium declined as capillary burden rose (Fig. 4b).

**Fig. 4.**
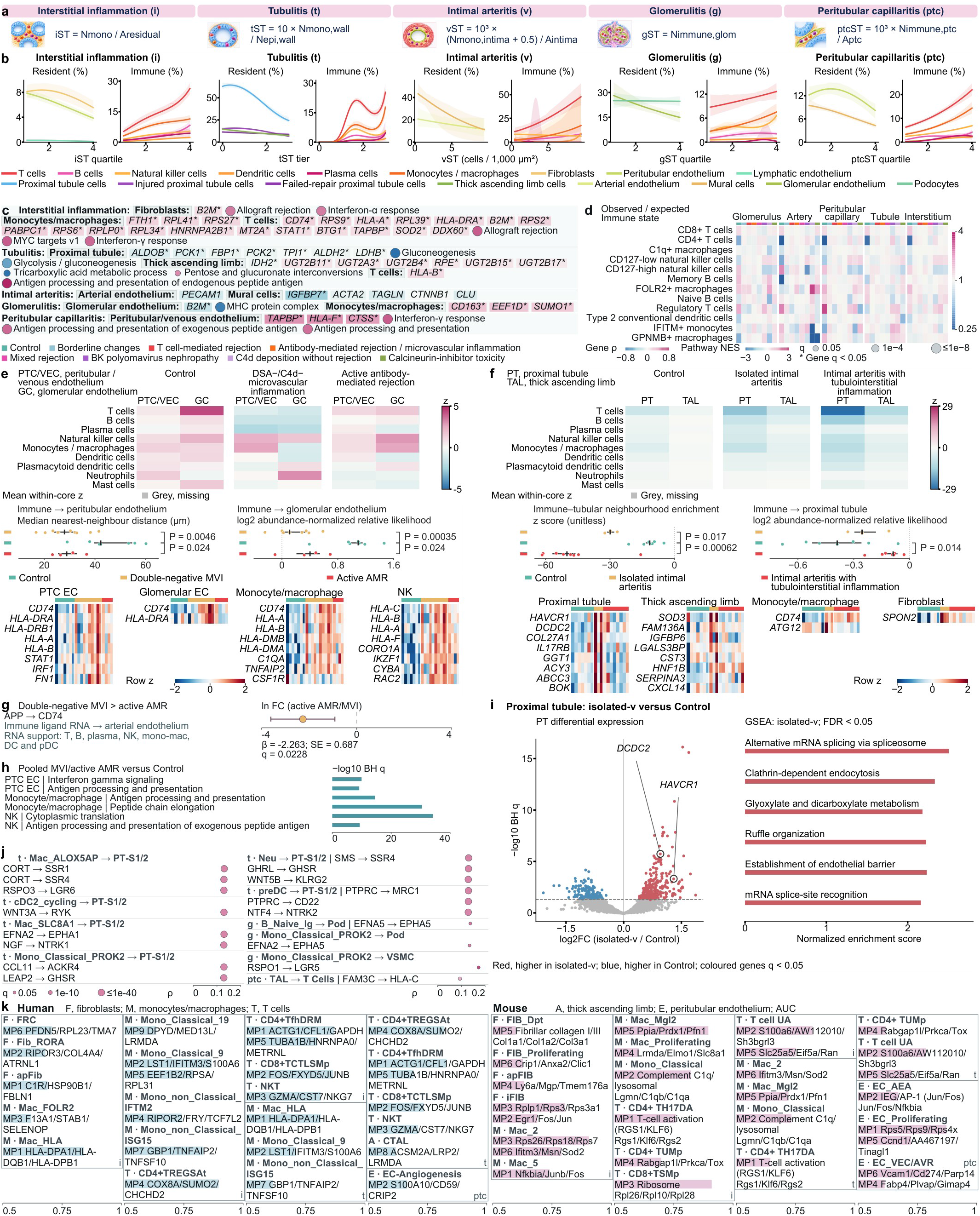
| Acute lesions elicit compartment-specific endothelial and tubular responses. **a**, Anatomical definitions of interstitial, tubulitis, arterial, glomerular and peritubular-capillary inflammatory burden. **b**, Compartment cell fractions across burden tiers; bands show pointwise 95% bootstrap intervals with biopsies as resampling units. **c**, Gene–burden correlations and pathway enrichment in 11 cellular contexts; stars denote gene-family q<0.05. **d**, Observed/expected ratios (Ro/e) of immune states within their parental classes. **e,f**, Core-level spatial organization and row-z-scored expression. Points denote cores; marks show medians and interquartile ranges. Displayed two-sided Mann–Whitney P values satisfy within-metric BH q≤0.05. In e, distance and relative-likelihood comparisons use 7/10/4 and 7/9/4 Control/MVI/AMR cores, and glomerular expression uses 5/8/4. In f, Control, isolated v and arteritis with tubulointerstitial inflammation include 7, 3 and 8 cores. **g**, Arterial-endothelial APP–CD74 receiver scores are higher in double-negative microvascular inflammation (MVI) than in active antibody-mediated rejection (AMR; 10/4 cores). The point gives the active-AMR/MVI natural-log coefficient, and the whisker an unadjusted 95% Wald interval. Ligand-source labels indicate RNA expression. **h**, Pathways over-represented among genes higher in pooled MVI and active AMR than in Control. **i**, Proximal-tubule isolated-v-versus-Control volcano plot (3 and 7 cores; 3,702 and 13,978 cells) from an NB2 smiDE mixed model with a Matérn spatial term across 17,361 genes; coloured genes have whole-family BH q<0.05. Right, pathways with positive normalized enrichment scores (NES) at GSEA FDR<0.05 (1,000 permutations; gene sets of 10–500 genes). **j**, Lesion–receiver-score Spearman associations across anatomical units. **k**, Human and mouse programme gene-rank AUC, the probability that a programme member ranks above a background gene (null, 0.5). Displayed enrichments, routes and programmes meet their family q<0.05. Units, complete counts and test families are detailed in Methods and Source Data.

We then related the transcriptional programmes of cells in each compartment to inflammatory burden and found that each compartment responded with its own programme (Fig. 4c). With interstitial burden, monocytes/macrophages increased HLA-DRA, B2M, STAT1, TAPBP, MT2A and SOD2, with enrichment of allograft-rejection, interferon-γ-response and MYC-target pathways, and T cells increased CD74, HLA-A, HLA-DRA and B2M. Tubular segments responded differently to tubulitis. Proximal tubules reduced ALDOB, PCK1, FBP1, PCK2, TPI1, ALDH2 and LDHB, with loss of gluconeogenesis, and thick ascending limb cells reduced IDH2 and UDP-glucuronosyltransferase genes. Mural cells reduced IGFBP7 with arterial burden. The two capillary beds responded in opposite directions. As capillary burden rose, peritubular endothelium increased TAPBP, HLA-F and CTSS, with enrichment of interferon-γ response and antigen processing. Glomerular endothelium instead reduced B2M and MHC protein complex genes with glomerular burden, as glomerular monocytes/macrophages increased CD163, EEF1D and SUMO1 (Fig. 4c). Endothelial HLA-F marks poor prognosis in antibody-mediated rejection^39^, and these data place its induction in the capillary bed in which inflammation accumulates. Immune subtypes within lesions also differed by diagnosis. Within the same lineage and compartment, observed-to-expected ratios (Ro/e)^40^ calculated by diagnosis resolved enriched subtypes. CD4 T cells and GPNMB-positive macrophages were enriched in tubules with BK polyomavirus nephropathy (Ro/e = 1.41 and 1.34), and IFITM-positive monocytes in AMR/MVI glomeruli (Ro/e = 1.28; Fig. 4d). Infiltrate immunophenotypes overlap between BK polyomavirus nephropathy and TCMR^41^, and these tubular states are candidates for distinguishing the two.

Double-negative MVI is an incomplete phenotype within the AMR/MVI spectrum whose clinical significance was highlighted separately in the Banff 2022 and Banff 2024 reports^1,3^. Spatially resolved transcriptomics allowed us to examine the spatial relationships between immune cells and endothelium in double-negative MVI and active AMR. In both groups of cores, immune cells lay closer to peritubular endothelium than in Control (median 28.0 and 28.9 µm versus 42.3 µm; P = 0.0046 and 0.024; Fig. 4e). With microvascular inflammation, peritubular endothelium expressed more CD74, HLA-DRA, HLA-DRB1, HLA-A, HLA-B, STAT1, IRF1 and FN1, and glomerular endothelium more CD74 and HLA-DRA. Monocytes/macrophages and natural killer cells also expressed more HLA class I genes (Fig. 4e). The two microvascular conditions differed in arterial signalling. APP–CD74 reception by arterial endothelium, with APP expressed by T, B, plasma, natural killer, monocyte/macrophage and dendritic cells, was higher in double-negative MVI than in active AMR (β = −2.263, q = 0.0228; Fig. 4g). Genes higher in the pooled microvascular conditions than in Control were enriched for interferon-γ signalling and antigen processing in peritubular endothelium (Fig. 4h), as in the peritubular response to capillary burden (Fig. 4c). Double-negative MVI carries a graft-loss risk similar to that of AMR^7^, and in this cohort both conditions brought immune cells closer to peritubular endothelium. Although it is an incomplete phenotype, double-negative MVI therefore resembled AMR more than rejection-free controls at the molecular level. Whether it warrants AMR-like intervention deserves further evaluation.

Isolated intimal arteritis (isolated v), which lacks tubulointerstitial inflammation above threshold, is another distinctive phenotype whose status as rejection has long been debated^3,42–44^. In isolated-v cores, the neighbourhood enrichment z-score of immune cells with tubules was lower than in Control (−29.9 versus −11.5; P = 0.017). A negative z-score means that the two cell classes are adjacent less often than expected under random permutation, and lower values indicate stronger avoidance of tubules by immune cells. The abundance-normalized likelihood of immune cells neighbouring proximal tubules did not differ from Control (P = 0.38; Fig. 4f). Cores with intimal arteritis and tubulointerstitial inflammation also had a lower z-score (P = 0.00062) but a higher likelihood of immune– proximal-tubule contact (P = 0.014). Although immune cells did not approach proximal tubules in isolated v, the tubules still responded. Among the three groups, proximal tubules expressed HAVCR1, DCDC2, COL27A1, IL17RB, GGT1, ACY3 and ABCC3 most highly in isolated-v cores (Fig. 4f). A cell-level mixed model of 3,702 proximal tubule cells from three isolated-v cores and 13,978 from seven Control cores identified 562 genes at q < 0.05. These included HAVCR1 (log2FC = 1.30, q = 4.8 × 10⁻⁴) and DCDC2 (log2FC = 0.95, q = 1.5 × 10⁻⁶). Positively enriched pathways included alternative mRNA splicing (NES = 2.50), clathrin-dependent endocytosis, glyoxylate and dicarboxylate metabolism, ruffle organization and establishment of endothelial barrier (Fig. 4i). DCDC2 marks the adaptive or failed-repair proximal tubule state in human and mouse kidneys^23,24^, and HAVCR1 encodes the injury marker KIM-1. Isolated arteritis is therefore accompanied by a failed-repair-associated tubular programme while immune cells remain apart from tubules.

Communication and single-cell programmes linked these lesion responses across compartments and species. In interactions between immune cells and PT S1/2 cells, scores of several ligand–receptor pairs rose with tubulitis burden (Fig. 4j). They included CORT– SSR1, CORT–SSR4 and RSPO3–LGR6 from ALOX5AP-positive macrophages, WNT3A– RYK from cycling cDC2, CCL11–ACKR4 and LEAP2–GHSR from PROK2-positive classical monocytes, and GHRL–GHSR and WNT5B–KLRG2 from neutrophils. In glomeruli, naive B cells and PROK2-positive classical monocytes signalled to podocytes through EFNA5– EPHA5 and EFNA2–EPHA5, and with capillaritis the thick ascending limb engaged T cells through FAM3C–HLA-C. Lesion-associated genes matched human and mouse programmes in the corresponding cell classes, including an HLA class II macrophage programme in human interstitium and a Vcam1/Cd274/Parp14 endothelial programme for mouse capillaritis (Fig. 4k). Scored in mouse single cells, the spatially derived signatures followed the mouse Banff grades. Interstitial monocyte/macrophage and proximal-tubule inflammation scores rose with i, the thick ascending limb tubulitis-down score fell with t, and the monocyte/macrophage capillaritis score rose with ptc (all q < 0.05; Extended Data Fig. 4a). In an independent RNA-seq cohort of 28 biopsies, the glomerulitis-up signature rose with g and the tubulitis-down signature fell with t (ρ = 0.715 and −0.735; q = 0.000225 for both). The interstitial monocyte/macrophage and peritubular endothelial signatures rose with i and ptc (Extended Data Fig. 4b).

Lesion-associated states also co-occurred across mouse grafts. Among cell states that co-varied across individual mice, interferon-stimulated macrophages correlated with Fut9-positive fibroblasts (Pearson r = 0.820) and with adaptive thick ascending limb cells (r = 0.788). These three states formed the leading combination among 216 three-state combinations that spanned three parent lineages (Extended Data Fig. 5a–c). Acute lesions thus engage immune cells, endothelium and tubular epithelium in compartment-specific ways, and the co-occurrence of macrophage, fibroblast and injured epithelial states in mouse grafts points to their persistence into remodelling. We next examined the chronic lesions in which scarring and inflammation coexist.

### The landscape of chronic lesions reveals sustained immune secretory programmes and loss of metabolic programmes

We mapped interstitial fibrosis, tubular atrophy and arterial intimal thickening onto measured structures (Fig. 5a,b). As for acute lesions, pathologists manually annotated these structures, which were then segmented by our trained in-house deep-learning models and reviewed by pathologists. Structural fibrosis (ciST) is the scarred fraction of residual interstitial area, and atrophy (ctST) the loss of tubular diameter relative to non-atrophic reference tubules in the same core. Intimal thickening (cvST) is the intimal share of the combined intimal and luminal area. Yi et al. used deep learning to quantify interstitial, tubular and mononuclear-leukocyte lesions on whole-slide images and built composite scores from them^45^. Here, all three measures are computed within the same core with structure-specific denominators and are therefore relative fractions, so no further cross-slide standardization was applied. In i-IFTA regions, plasma cells as a fraction of all cells rose with clinical i-IFTA grade (ρ = 0.391, P = 0.0029; Fig. 5c), consistent with their increase across grades in scarred interstitium (Fig. 1f). ciST measures scar extent and clinical i-IFTA grades inflammation within scar, and both describe the microenvironment in which plasma cells and other immune cells reside.

**Fig. 5.**
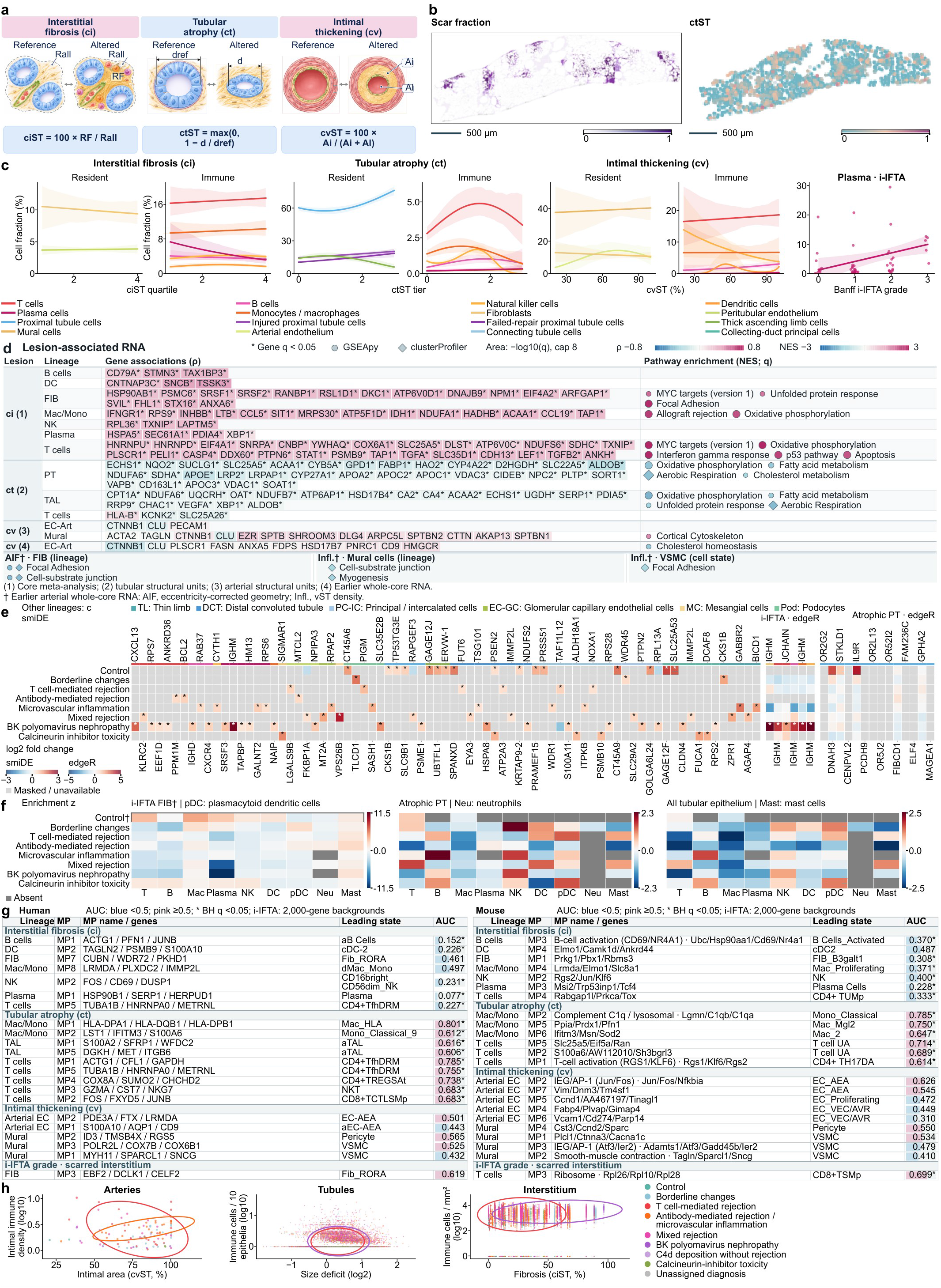
| Chronic lesions combine sustained immune secretory programmes with loss of mature tubular function. **a**, Structural definitions: ciST=100×RF/Rall, ctST=max(0,1−d/dref), and cvST=100×Ai/(Ai+Al). RF and Rall denote scarred and total residual-interstitial areas; d and dref denote minimum Feret tubular diameter and matched reference diameter; Ai and Al denote intimal and luminal areas. **b**, Fibrosis and tubular atrophy in a representative biopsy. Both maps use equal physical x/y scale; each coloured point on the tubular map is a ctST-scored structure. Pale points are unscored. Scale bars, 500 µm. **c**, Cell fractions along the chronic-lesion axes. Solid curves are descriptive logistic-polynomial estimates; shading denotes pointwise 95% intervals from 200 bootstrap samples of complete sampling units. Polynomial degree 1–3 was selected by leaving one unit out at a time, using fractional-binomial log loss and equal total fitting weight per unit. Unit identities follow each compartment analysis; physical cores from the same biopsy remain separate where defined in that analysis. Fractions use all assigned compartment cells. Tubular-wall denominators include immune and epithelial cells; parent proximal-tubule, injured and failed-repair populations overlap. The i-IFTA facet shows 56 biopsy observations and the 80-point weighted least-squares mean and 95% interval for the plasma-cell fraction. This facet uses clinical i-IFTA grade; the other facets use measured structural scores. The shared legend below identifies curve cell types; e uses these colours with six additional cell-type entries. **d**, Cell-specific RNA responses to chronic lesions. Lesion, cell class, genes and pathways occupy aligned columns. The 150 gene tiles show Spearman correlations with lesion loading; asterisks denote gene-family q<0.05. The 27 pathway readouts are arranged under 15 full biological pathway names. Colour denotes signed normalized enrichment score (NES), area denotes capped −log10(q), and circles and diamonds distinguish GSEApy and clusterProfiler results. All displayed pathways have their enrichment-test-family q<0.05. Cholesterol homeostasis in whole-core arterial endothelium has NES=−1.710, q=0.00713; proximal-tubule cholesterol metabolism along ct has NES=−1.456, q=0.04743. Mural-cell cortical cytoskeleton enrichment along cv has NES=1.855, q=0.03765. The seven readouts marked with a dagger use whole-core RNA from the joint arterial analysis: fibroblasts were related to the eccentricity-corrected geometry axis, and mural or vascular smooth-muscle cells to the inflammation-density axis. Myogenesis has NES=−1.542, q=0.00207 in that whole-core analysis. The plasma-cell entries include HSPA5, SEC61A1, PDIA4 and XBP1 RNA across 32 pathology cores. Source Data provide gene sets, leading edges, software/library-specific statistics and observation units. Gene estimates without an asterisk are nonsignificant; finite-permutation q values reported as zero use the bubble-size cap. **e**, Diagnosis-versus-rest RNA effects. The smiDE block uses whole-core immune cells, scarred-interstitial fibroblasts, and tubular epithelial, peritubular endothelial or glomerular compartments (left). Local edgeR results show six cell-type/gene entries in i-IFTA residual interstitium (54 biopsies, middle) and 14 PT genes in atrophic outer walls (64 eligible biopsies, right), with biopsies as replicates. Control RnonF is excluded from i-IFTA comparisons. Local IGHM/JCHAIN estimates describe captured RNA associations within annotated level-1 lineages. In the smiDE block, coloured cells show significant log2 fold changes; local blocks show all estimable effects. Scales are capped at ±3 and ±5, respectively. Stars indicate positive effects with the corresponding model-family BH FDR<0.05; PT14 has no positive FDR result. Grey denotes nonsignificant or missing smiDE values, or untested/unestimated local values. Lineage colours follow the shared key in c and six additional full names above. Complete testing families and biopsy counts per contrast are supplied in Source Data. **f**, Squidpy neighbourhood enrichment in three cellular scopes: i-IFTA residual-interstitial fibroblast–immune pairs (left), atrophic-wall PT–immune pairs (middle), and all seven pooled tubular epithelial classes in the same walls (right). Case interstitium and atrophic walls contain 54 and 79 physical cores, respectively. The pooled target contains 10,530 cells within an 11,971-cell mask. The separately framed Control-fibroblast row (†) uses unscarred residual interstitium (RnonF) as an anatomical reference: 9,888 cells including 923 fibroblasts from seven biopsies. Tubular Control rows use the atrophic-wall mask. Masks induce local subgraphs of full-core 30-µm Delaunay graphs, with 1,000 within-core label permutations and seed 42. Views use separate symmetric z scales. Each contains 72 tiles, giving 216 in total; 70, 56 and 56 values are finite. Grey denotes cell classes absent from the mask (2, 16 and 16 tiles). T, T cells; B, B cells; Mac, monocytes/macrophages; Plasma, plasma cells; NK, natural killer cells; DC, dendritic cells; pDC, plasmacytoid dendritic cells; Neu, neutrophils; Mast, mast cells. Z scores describe within-mask neighbourhood organization; between-diagnosis tests were not performed. Complete matrices, cell counts and absence causes are supplied in Source Data. **g**, Rank correspondence of human and mouse metaprogrammes to genes associated with interstitial fibrosis (ci), tubular atrophy (ct), intimal thickening (cv) and clinical i-IFTA grade. The 45 estimates comprise the highest ci AUC within each measured lineage and species (14 results), 15 significant positive ct results, all 14 estimable cv results, and the highest i-IFTA AUC in each species. Human and mouse results occupy separate blocks, with thin rules separating lesions. Aligned columns show the lineage, MP identifier, programme name with representative and explicitly named programme genes, leading cell state and AUC. The leading cell state is the annotated subtype with the highest mean programme expression. AUC=U/(nMP×nbackground), using all finite nonmembers in the same lesion/cell ranking. The numerical tracks span 0–1; blue values below 0.5 and pink values at or above 0.5 indicate lower and higher lesion-correlation ranks among programme members, respectively. Asterisks denote BH q<0.05 in each complete testing family. The 264 lesion/cell/MP tests use separate species-wide BH families of 142 human and 122 mouse comparisons. The 38 clinical i-IFTA tests use separate BH families of 21 human and 17 mouse comparisons. Each test ranks year/depth-residualized expression against clinical grade by Spearman correlation in scarred-interstitial fibroblasts, monocytes/macrophages and T cells, on 2,000 high-variance genes per lineage. Human i-IFTA and all cv estimates are nonsignificant. Complete lesion/MP results, independent i-IFTA families, all directions and nonsignificant estimates, and separate sample-score–metaprogramme correlations are supplied in Source Data. **h**, Joint inflammation–morphology distributions of matched anatomical structures, coloured by diagnosis: 99 arterial, 11,120 tubular and 10,000 interstitial observations. Contours denote descriptive 80% bivariate-normal covariance regions. Arterial contours compare T cell-mediated rejection with the combined antibody-mediated rejection/microvascular-inflammation category; tubular and interstitial contours show T cell-mediated rejection and BK polyomavirus nephropathy. n denotes structures. The adjacent legend identifies diagnosis colours.

Cell-specific RNA programmes followed each lesion axis (Fig. 5d). As ciST rose, plasma cells increased HSPA5, SEC61A1, PDIA4 and XBP1. These endoplasmic-reticulum chaperone, translocon and transcription-factor genes expand the secretory apparatus during plasma-cell differentiation^46^. In the same scarred interstitium, macrophages increased IFNGR1, LTB, CCL5, CCL19 and TAP1. T cells increased STAT1, PSMB9, TAP1, LEF1 and TGFB2, with enrichment of interferon-γ, p53 and apoptosis pathways, and fibroblasts showed unfolded-protein-response and focal-adhesion programmes. As ctST rose, proximal tubules reduced FABP1, ACAA1, HAO2 and ALDOB, with loss of fatty-acid oxidation, oxidative phosphorylation and cholesterol metabolism. Thick ascending limb cells reduced CPT1A together with oxidative phosphorylation and unfolded-protein-response programmes, including XBP1. This is consistent with the known promotion of kidney fibrosis by defective fatty-acid oxidation in tubular epithelium^47^. Along cvST, mural cells increased cortical cytoskeleton genes and arterial endothelium reduced cholesterol homeostasis, suggesting broad transcriptional remodelling of the arterial wall. Chronic lesions thus combine plasma-cell secretory adaptation, persistent interferon-associated T-cell and macrophage programmes and loss of tubular fatty-acid oxidation.

The same lesions also carried distinct transcriptional signals in different diagnostic categories (Fig. 5e). BK polyomavirus nephropathy and TCMR can both cause tubulointerstitial inflammation and can be difficult to distinguish histologically. Molecularly, however, BK polyomavirus nephropathy combined higher T-cell CXCL13 (log2FC = 1.75, q = 3.6 × 10⁻¹⁰), B-cell CXCR4 and IGHD, and plasma-cell IGHM (log2FC = 2.83, q = 1.9 × 10⁻⁶), SRSF3, TAPBP and HM13. Mixed rejection was associated with higher T-cell KLRC2 (log2FC = 0.69, q = 0.0023), glomerular endothelial MT2A (1.19, q = 0.0015) and mesangial VPS26B (2.38, q = 0.0099). Microvascular inflammation was associated with higher B-cell RAB37, plasma-cell GALNT2, natural killer CYTH1 and glomerular endothelial RPAP2. AMR carried higher T-cell BCL2 and PPM1M and peritubular endothelial MTCL2, and TCMR higher peritubular endothelial LGALS9B. Calcineurin-inhibitor toxicity was associated with higher fibroblast SIGMAR1 in scarred interstitium (log2FC = 1.44, q = 0.017). Proximal tubules in atrophic walls, by contrast, showed no diagnosis-specific increase, suggesting that tubular atrophy may share pathways across categories. IgM-expressing plasma cells and CXCL13-expressing T cells, which also mark TLS interiors (Fig. 3j), together characterize BK polyomavirus nephropathy, whose infiltrates are difficult to distinguish from TCMR by immunophenotyping alone^41^. Neighbourhood maps placed immune classes relative to fibroblasts in scarred residual interstitium and to proximal tubules and pooled tubular epithelium in atrophic walls, with unscarred Control interstitium as an anatomical reference (Fig. 5f).

These lesion-specific programmes were also reflected in single-cell data from both species. The atrophy-associated monocyte/macrophage signal was linked to a human HLA-DPA1/HLA-DQB1/HLA-DPB1 programme and a mouse complement C1q and lysosomal programme (AUC = 0.801 and 0.785; q < 0.001). T-cell programmes were also positively associated in both species (Fig. 5g). Atrophy-down signatures of proximal tubule and thick ascending limb cells fell with Banff ct in human single-cell data (ρ = −0.483 and −0.544). The same signatures fell with ct grade in mouse grafts (ρ = −0.742 and −0.681; all q < 0.05; Extended Data Fig. 4c). In an independent biopsy RNA-seq cohort, the interstitial B-cell fibrosis-up signature rose with ci (ρ = 0.504, q = 0.0229). The proximal-tubule atrophy-down signature, which contains ALDOB, SLC22A6 and PCK1, fell with ct (ρ = −0.517, q = 0.0191; Extended Data Fig. 4d).

Unlike these cell-specific RNA states, joint inflammation–morphology distributions overlapped extensively between TCMR and BK polyomavirus nephropathy in tubules and interstitium, and between TCMR and AMR/MVI in arteries (Fig. 5h). Inflammation and morphology alone therefore did not distinguish these diagnoses. Human biopsies capture these states at one time point, so we turned to the mouse cohort to follow how immune and tubular states change as grafts move towards rejection or acceptance.

### Rejection and tolerance trajectories diverge in immune states and parenchymal recovery

We used PILOT^48^ to model sample-level trajectories of single-cell profiles from 34 mouse grafts across transplant models and sampling times. PILOT compares the cell-composition distributions of animals by optimal transport (Wasserstein distance), builds a sample tree from these distances and infers sample-level pseudotime; we rooted the tree in day-7 syngeneic grafts. The tree resolved a rejection-associated path of 21 animals and a tolerance-associated path of 15 animals, which shared two syngeneic root animals (Fig. 6a).

**Fig. 6.**
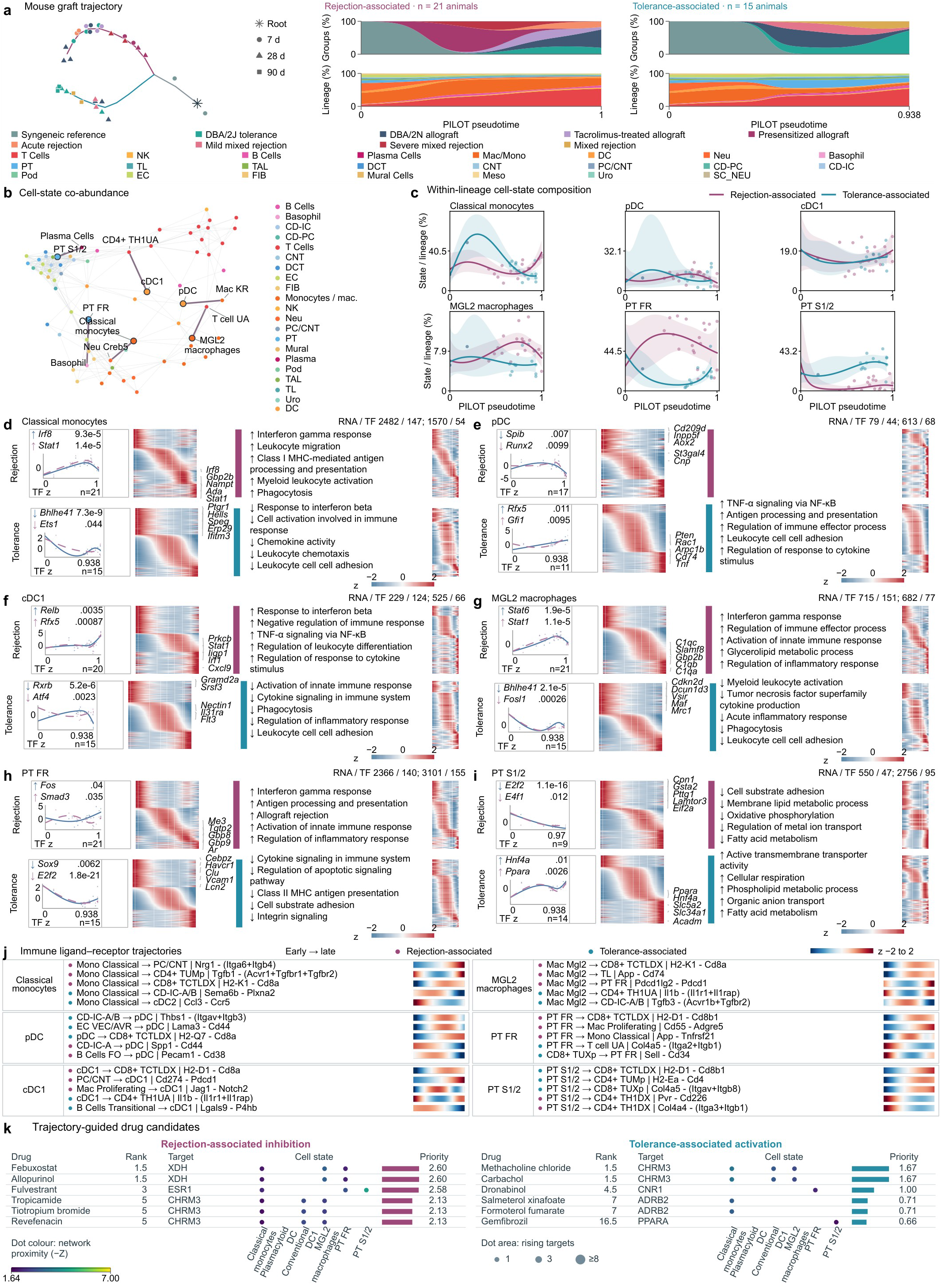
| Immune states, tubular programmes and candidate intervention targets along graft-level trajectories. **a**, PILOT trajectories rooted in day-7 syngeneic grafts, with diagnostic and lineage composition. Points denote animals; circles, triangles and squares indicate days 7, 28 and 90. Magenta and cyan denote rejection-associated and tolerance-associated paths. The paths contain 21 and 15 animals, including two shared-trunk animals, for 34 independent animals in total. **b**, Cell-state co-abundance network with 70 nodes and 359 positive edges (Pearson r>0.3 and BH q<0.05 across 2,415 pairs). Highlighted edges connect the six selected states to their strongest cross-lineage partners involving an immune state. **c**, Within-lineage proportions of classical monocytes, pDC, cDC1, MGL2 macrophages, PT FR and PT S1/2. Points denote animals; lines and shading show spline means and pointwise 95% intervals over each path’s observed support. **d–i**, Paired RNA, regulon and pathway profiles for the six states in panel c, with rejection above tolerance. Heatmaps show all 15,668 significant RNA and 1,168 TF profiles, ordered by fitted peak time and shown as row z scores. TF curves and points show fitted activity and animal AUCell means on a common within-regulon z scale across paths. Numbers beside TF names are association q values; arrows give fitted end-minus-start directions. Upper-right counts list significant RNA/TF profiles in rejection and tolerance. Directional ORA uses the sign of fitted RNA–pseudotime correlation. The decreasing pDC rejection foreground has no significant ORA term. **j**, Five distinct immune-involving ligand–receptor pairs per selected cell state. Rows name sender, receiver, ligand and modelled receptor subunits. Strips show row-standardized fitted response components from early to late; coloured dots identify paths. All displayed profiles have association q<0.05. Molecular BH families contain 686,304 RNA, 4,844 TF and 3,613,040 ligand–receptor hypotheses. **k**, Direction-matched scDrugPrio candidates. Each direction shows its five leading whole-RNA entries plus a Tier 3-supported candidate when available, otherwise the next entry. Tied candidates share their average rank. Dots identify qualifying cell-state contexts; colour denotes network proximity −Z and area the number of rising matched targets, capped at eight for display. Bars show priority scores on separate within-path scales. Target names are one-to-one human orthologues of mouse-source genes. Complete curves, pathway terms, pharmacological evidence and candidates are provided in Extended Data Fig. 8a–e and Supplementary Data 2.

The rejection-associated path gathered day-7 BALB/c acute-rejection, presensitised and tacrolimus-treated grafts, day-7 DBA/2J grafts, day-7 and day-28 DBA/2N grafts, and day-28 presensitised and severe mixed-rejection grafts. The tolerance-associated path gathered day-28 and day-90 DBA/2J grafts, day-28 DBA/2N and mild mixed-rejection grafts, and day-90 BALB/c grafts (Fig. 6a). Along the rejection path, presensitised grafts dominated the middle of the path (about 78% of the smoothed composition). The end of the path was dominated by DBA/2N grafts (55.4%), followed by day-7 DBA/2J (19.6%), BALB/c acute-rejection (16.9%) and tacrolimus-treated grafts (7.6%). The end of the tolerance path was dominated by DBA/2J grafts (72.1%), followed by day-90 BALB/c grafts (18.1%). On both paths, T cells rose from about 5% to about 54% of cells and B cells from 1.0% to about 7%. The paths differed in myeloid and tubular cells. Monocytes/macrophages first rose to 42.5% along the rejection path and remained at 22.6% at its end, but fell to 6.4% along the tolerance path. Proximal tubule cells fell from 7.9% to 2.2% along the rejection path and rose to 12.1% along the tolerance path, with a mid-path peak of 23.7%. Neutrophils and endothelial cells fell along the rejection path from 24.8% and 6.9% to 1.3% and 1.9%, and plasma cells appeared only at the end of the tolerance path (0.9%; Fig. 6a). Day-7 DBA/2J grafts lay on the rejection path, consistent with reports that accepted grafts also show early immune infiltration^29,30^. A co-abundance network of 70 subtypes and 359 positive edges connected classical monocytes, plasmacytoid dendritic cells (pDCs), cDC1s, MGL2 macrophages and proximal-tubule states with immune and tissue partners in other lineages (Fig. 6b and Extended Data Fig. 5d). Along the rejection path, the fitted endpoint fractions of classical monocytes and failed-repair proximal tubules (PT FR) rose within their parent lineages, and differentiated PT S1/2 cells declined. Along the tolerance path, PT S1/2 cells increased (Fig. 6c). We tested these changes with parent-lineage and all-cell denominators. Classical monocytes rose and then fell along the tolerance path (all-cell denominator, q = 0.0067). Within its parent lineage, PT FR rose to about 60% in the middle of the rejection path and remained above its starting level at the end. As a fraction of all cells, this change was significant (q = 0.00086). Along the tolerance path, PT FR fell below 10% mid-path. PT S1/2 and thick ascending limb cells increased late on the tolerance path without reaching significance (q ≥ 0.18; Extended Data Fig. 5e).

We then tested transcriptional changes of cell states along each path with tradeSeq^49^. Within these states, RNA and regulon profiles changed along each path (Fig. 6d–i). Along the rejection path, classical monocytes increased *Stat1*, *Irf8* and *Nampt*, cDC1s increased *Irf1* and *Cxcl9*, and MGL2 macrophages increased *C1qa*, *C1qb* and *C1qc*. Interferon-response, antigen-presentation and immune-effector programmes rose in these myeloid states and in PT FR. SCENIC^50,51^ regulon activity rose along the rejection-associated trajectory for *Irf8* and *Stat1* in classical monocytes, *Relb* and *Rfx5* in cDC1s and *Stat1* and *Stat6* in MGL2 macrophages, and fell for *Spib* and *Runx2* in pDCs. Along the tolerance path, PT FR cells reduced *Havcr1*, *Vcam1* and *Lcn2* and lost *Sox9* and *E2f2* regulon activity. PT S1/2 cells increased *Slc5a2*, *Slc34a1*, *Hnf4a* and *Ppara*, with rising *Hnf4a* and *Ppara* regulon activity and fatty-acid metabolism, respiration and transport programmes (Fig. 6i). Complete tolerance-path results for pDCs, PT S1/2 and thick ascending limb cells are shown in Extended Data Fig. 5f–i. PT S1/2 cells also increased MHC class II antigen-presentation genes such as *Cd74* and *H2-Aa* (*Esrra* and *Stat1* regulons, q = 0.014 and 0.024). Thick ascending limb cells instead reduced ribosomal and translation-initiation programmes (Extended Data Fig. 5g,h). The tubular programme recovered along the tolerance path is the one lost in human atrophic tubules (Fig. 5d), and *Havcr1*, induced in human proximal tubules beside isolated arteritis (Fig. 4i), declined in PT FR as grafts were accepted.

Ligand–receptor profiles also changed along the trajectories. Pdcd1lg2–Pdcd1 scores between MGL2 macrophages and PT FR, H2-D1–Cd8b1 scores between PT FR and CD8 T cells and Col4a5–integrin scores between PT S1/2 and T cells changed along the paths (all q < 0.05; Fig. 6j). Along the tolerance path, 18 directed ligand–receptor relationships reached q < 0.05. Most linked PT S1/2 cells through H2-D1 and H2-K1 to Cd8a/Cd8b1 on CD8 T cells, and through Col4a5–ITGAV/ITGB8 and H2-DMb1–Cd4 to CD4 Tregs. Tnf–Tnfrsf1a signalling from CD8 T cells to PT S1/2 cells was also among them (Extended Data Fig. 5j,k). The corresponding changes in pDC, PT S1/2 and TAL fractions are shown in Extended Data Fig. 5l.

In human single-cell data, we built a sample tree rooted in Control samples from 275,474 profiles in 94 tissue collections. It resolved a thick-ascending-limb-enriched reference path (60 samples) and two rejection-associated paths enriched for proximal tubule (39) or T cells (43; Extended Data Fig. 6a). Along these paths, the within-parent fractions of natural killer cells, CD8 T cells, CCD-IC-A, CNT and aPT followed different courses, but none was associated with pseudotime at q < 0.05 (Extended Data Fig. 6b). Path composition by fine diagnosis is shown in Extended Data Fig. 6c–e. Along the proximal-tubule-enriched path, CCD-IC-A cells increased HNF4A and NR1H4 regulon activity (q = 8.3 × 10⁻⁶ and 9 × 10⁻⁷) with transport and xenobiotic-metabolism transcripts. RNA and transcription-factor associations were detected on all three paths (Extended Data Fig. 7a–e), and drug2cell target scores for each state are shown in Extended Data Fig. 7f.

To identify druggable targets that change along the trajectories, we first used NicheNet^52^ priors to build intercellular networks among mouse cell states on each path (Extended Data Fig. 8a). We then ranked candidates with scDrugPrio^53^, which combines trajectory-associated programmes with degree-matched protein-network proximity and cell-state centrality (Extended Data Fig. 8b). Direction-matched screening nominated 34 inhibitors of rising rejection-associated signals and 11 activators of rising tolerance-associated signals (Fig. 6k and Extended Data Fig. 8c). Febuxostat and allopurinol ranked jointly first among rejection candidates through XDH in classical monocytes, MGL2 macrophages and PT FR. Xanthine oxidase inhibition with febuxostat protects the kidney from ischaemia–reperfusion injury by reducing oxidative stress^54^. Methacholine chloride and carbachol ranked jointly first among tolerance candidates through CHRM3, and the PT S1/2 PPARA programme linked to gemfibrozil with RNA, regulon and ligand–receptor support. Mechanism maps connected rejection candidates to XDH, ESR1, CHRM3, AR, ADA and MET, and tolerance candidates to CHRM3, ADRB2, CNR1, PPARA and VDR (Extended Data Fig. 8d). For the top five candidates in each direction, intercellular centrality of qualifying cell contexts accounted for almost all of the score (Extended Data Fig. 8e). Among states that rose along the rejection path, classical-monocyte *Ada* corresponded to the adenosine deaminase inhibitor pentostatin (RNA q = 0.0391). PT FR *Ar* corresponded to six androgen receptor antagonists (q = 0.0478), whose target expression was concentrated in PT FR (Extended Data Fig. 8i,k,m). Along the tolerance path, the TNF–TNFRSF1A score of PT S1/2 rose (q = 0.0365) and corresponded to the TNFR1 agonist tasonermin, whose effect on kidney allograft tolerance has not been tested (Extended Data Fig. 8j). Strict tier 2 drug-ingredient lists contained 204 human and 426 mouse ingredients, 190 of which were shared (Extended Data Fig. 8g,h). Lesion-associated core genes, target expression across mouse and human states, and the separate TAL drug screen are shown in Extended Data Fig. 8f and Extended Data Fig. 8k–n.

To compare states between human and mouse rejection, and to assess how far mouse rejection models represent human rejection, we used CellSpectra^55^ to compare mouse trajectory states with human rejection references. Across endothelial, myeloid and tubular populations represented in all three clinical references, mouse expression fitted the human AMR reference more closely than the others. We further assessed pathway coordination in 13 mouse model–time conditions. They comprised day-7 syngeneic, BALB/c acute-rejection and tacrolimus-treated grafts, presensitised grafts on days 7 and 28, and BALB/c mixed rejection on day 28 (mild and severe) and day 90. DBA/2N grafts were sampled on days 7 and 28, and DBA/2J grafts on days 7, 28 and 90. Classical monocytes and PT FR, which rose along the rejection path, were paired with 15 and 5 human states of the same lineage, respectively. These 20 pairs gave 260 pair–condition combinations across the 13 conditions (Extended Data Fig. 7g). Day-28 DBA/2N PT FR and human PT FR formed the pair with the most reference-preferential pathways, 44 of 69 evaluable gene sets (six animals). In this comparison, *Ar* and *Cd44* had positive residuals in the early oestrogen-response programme, and Cd44 also had positive residuals in the epithelial–mesenchymal transition programme. The source states were PT FR for Ar and classical monocytes for Cd44 (Extended Data Fig. 7h,i). The mouse trajectories thus identify interferon-activated myeloid states and persistent tubular injury states, and point to the particular importance of tubular metabolic recovery in tolerance.

### Plasma-cell ecology dominates a multilevel model of functional non-response

We classified 20 recipients with rejection-spectrum diagnoses, all of whom received standard treatment after biopsy-proven rejection, as responders (11) or non-responders (nine). Response required an eGFR increase of at least 10% and a creatinine decrease of at least 10% between 30 and 180 days after biopsy (Fig. 7a). Biopsies from non-responders had higher interstitial fibrosis scores and a longer interval from transplantation to biopsy (both q = 0.0363 across 24 baseline comparisons; Fig. 7b).

**Fig. 7.**
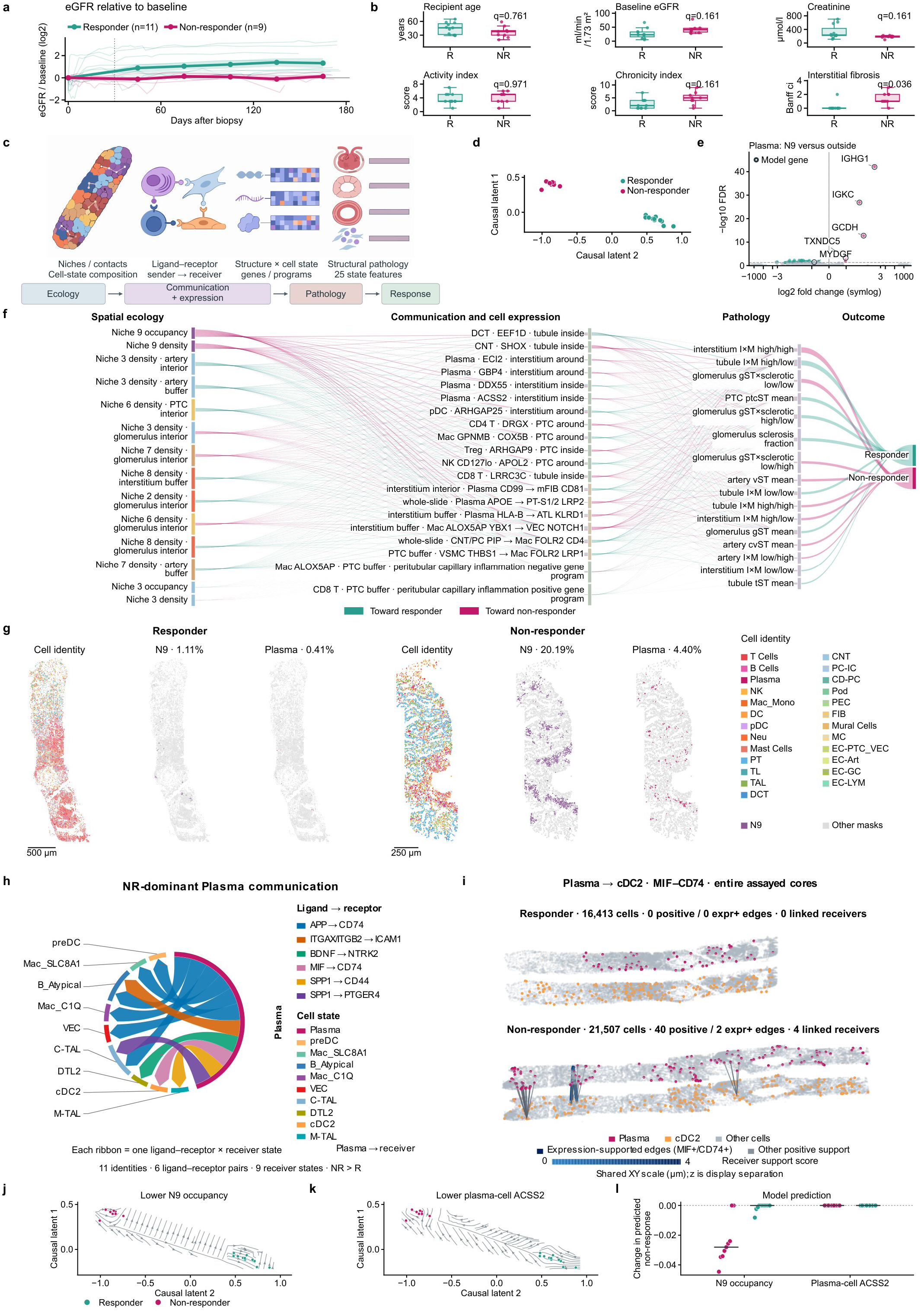
| Spatial ecology, molecular states and structural pathology associated with functional response. **a**, Observed log₂(follow-up eGFR/pre-biopsy baseline eGFR) in 11 responders and nine non-responders. Thin lines represent patients; thick lines connect patient-equal means in 30-day windows, and bands show s.e.m. Baseline is the median observed daily eGFR on days −30 to 0; day 0 is zero by definition. **b**, Age, baseline eGFR and creatinine, activity and chronicity indices, and interstitial fibrosis. Each point represents an observed patient. Boxes show medians and interquartile ranges; whiskers extend to observations within 1.5 interquartile ranges. Two-sided Mann–Whitney U tests used tie and continuity correction. q values use BH correction across all 24 estimable baseline comparisons. **c**, CauTrigger decoder cascade from spatial ecology through communication and expression to structural pathology and response. Ecology is encoded; measured molecular and pathology layers are reconstruction targets. **d**, Fitted two-dimensional causal latent coordinates of the 20 patients, coloured by observed response and displayed with equal x/y unit scaling. Patient-held-out performance is reported in the text and Source Data. **e**, Plasma-cell expression inside N9 versus pooled outside regions. All 1,258 finite patient-level meta-analysis results are shown; 83 have full-family BH q<0.05, including five higher inside N9. Rings identify IGHG1, IGKC, GCDH, TXNDC5 and MYDGF among the model’s plasma-cell gene features. Forty patients contribute at least one usable effect from 42 identity-linked cores representing 41 patients; gene-specific support is 9–40 patients. Repeated cores are combined within patient. The x axis uses a symmetric logarithmic scale, linear for |log2 fold change|≤3 (linear-region scale factor, 5); y is −log10[max(q,10⁻³⁰⁰)]. **f**, Local sensitivities of the non-response logit across ecology, the molecular layer, pathology and response. The network shows 14 ecological, 20 molecular, 16 pathology and two outcome nodes. Magenta and teal ribbons indicate positive and negative sensitivities, respectively; widths are normalized within each layer boundary. Niche colours follow Fig. 2. Labels specify cell identity, molecular feature, anatomy and spatial scope. The expanded network includes 86 molecular nodes and all 25 pathology features. **g**, Cell masks in complete TCMR cores from one responder and one non-responder. Each core is shown by level-1 lineage, N9 and plasma-cell identity. The cores contain 16,413 and 5,048 cells, including 182 and 1,019 N9 cells and 68 and 222 plasma cells, respectively. Background masks show all remaining cells. The three views within each core share a scale; cores are fitted to the same displayed height at different magnifications. Scale bars, 500 µm (R) and 250 µm (NR). **h**, Eleven plasma-cell communication identities with higher patient-equal NR values for both adjacency-attributed scores and positive neighbourhood support. Arrows run from plasma cells to nine receiver cell states. Ribbon colour identifies one of six ligand–receptor pairs; sector colour identifies the receiver cell state. Equal ribbon widths encode the presence of each ligand–receptor–receiver combination. Group means include nine NR and 11 R patients. **i**, Whole-core plasma-cell-to-cDC2 MIF–CD74 neighbourhoods in the responder core of g and a second non-responder core. The cores contain 16,413 and 21,507 cells. The NR core contains 40 positive support edges to four cDC2 receivers; two edges with detected sender MIF and receiver CD74 are highlighted. The R core has zero plasma-linked positive edges. Both views use shared physical coordinate limits, camera, score range and edge rules. Vertical spacing separates sender and receiver display planes. **j,k**, CauTrigger streamline plots in the causal latent space after whole-biopsy N9 occupancy (j) or the two interstitial views of plasma-cell ACSS2 (k) were capped at their training-set 10th percentile. All other inputs of the fitted model kept their observed values. Points are the 20 patients, coloured by observed response, with the coordinates and display limits of d. After N9 occupancy was lowered, six of nine non-responders flowed towards a responder-like state and 10 of 11 responders retained a responder-like state. The other two non-responders already had zero N9 occupancy, so their pathology state did not change. After plasma-cell ACSS2 was lowered, three of nine non-responders flowed towards a responder-like state, and responder flows pointed towards the non-responder side. Flow definitions are given in Methods. **l**, Change in each patient’s predicted probability of non-response relative to the fitted baseline; lines show group medians. Lowering N9 occupancy reduced the predicted probability in non-responders by 0.025 on average, whereas lowering ACSS2 changed it by less than 10⁻⁵ in either group. Direction-concordant plasma-cell communication features lay below the cap in every patient, so this layer was not perturbed. mFIB, myofibroblasts; R, responder; NR, non-responder. Complete estimates, flows and probability changes are provided in Source Data.

To test whether the transcriptional state of single structures could explain differences in response, we modelled expression-based trajectories separately in structural units, including glomeruli, tubules, arteries, peritubular capillaries, interstitium and lymphoid aggregates. This yielded 59 Slingshot^35^ lineages across 25 anatomical scopes, on which we overlaid lesion scores, diagnoses and response (Extended Data Fig. 9a–f). In glomeruli, for example, 461 units formed three lineages. Lineage 1 was dominated by AMR/MVI (37%) and mixed rejection (27%), and lineage 3 by AMR/MVI (29%) and TCMR (23%). Lineage 2 contained more sclerotic glomeruli (20%) and was enriched for TCMR (26%) and borderline changes (19%) (Extended Data Fig. 9a). The 13,093 interstitial units formed a single lineage. Among 2,974 response-labelled units from 19 biopsies, responder units lay slightly later in pseudotime (Extended Data Fig. 9d).

We then used the hierarchical causal-inference method CauTrigger^56^ to encode spatial ecology and to decode, in turn, cell communication, gene expression, structural pathology and response (Fig. 7c). This design resolves how spatial ecology influences transcriptional programmes and pathological burden. In the causal latent space, the fitted latent coordinates separated responders from non-responders well (Fig. 7d). In patient-held-out five-fold evaluation, the AUROC was 0.949 and the Brier score 0.095.

Among ecological features, N9 occupancy and density had the largest local sensitivities, and both raised the non-response logit (Fig. 7f). Two whole TCMR cores illustrate this difference. N9 cells and plasma cells made up 1.11% and 0.41% of cells in a responder core, and 20.19% and 4.40% in a non-responder core (Fig. 7g). In an alternative model that encoded communication as receiver-cell signal scores, N9 occupancy and density likewise had the largest local sensitivities (AUROC 0.939, Brier score 0.106; Extended Data Fig. 9g). Genes of high importance in this model were enriched for mitochondrial envelope and mitochondrion in regulatory T cells and for endosome in AVR endothelium (FDR = 0.0286, 0.0367 and 0.0366; Extended Data Fig. 9h). The highest-ranked downstream communication feature of the main model was plasma-cell APOE signalling to PT S1/2 LRP2 (rank 20 in the full model), whose sensitivity raised the non-response logit. Each of the 50 immune-cell ligand–receptor identities was positive in only 1–3 of the 20 patients (Extended Data Fig. 9i).

We next focused on the properties of plasma cells inside N9 and their possible biological effects. Plasma cells inside N9 differed in expression from plasma cells outside it (Fig. 7e). Using smiDE Optimal models to correct for transcript spillover from neighbouring cells, followed by patient-level meta-analysis, we detected 83 differentially expressed genes (q < 0.05). Five were higher inside N9: IGHG1 (log2FC = 2.65), IGKC (1.78), GCDH (2.02), TXNDC5 (0.97) and IGLL5 (0.96). The other 78 were lower, among them MYDGF (−0.86, q = 0.034), PAX5 (−2.54) and the proliferation-associated HJURP and NUF2. High immunoglobulin heavy-chain and light-chain expression with loss of the B-cell identity factor PAX5 and of proliferation genes is consistent with a terminally differentiated, non-dividing state inside N9. TXNDC5 is an endoplasmic-reticulum protein disulfide isomerase that, in kidney fibroblasts, promotes fibrosis by enforcing TGF-β signalling^57^.

Plasma cells in N9 lay next to many myeloid, B-cell, vascular and tubular cells. Eleven ligand–receptor–receiver combinations had higher values in non-responders for both plasma-adjacency-attributed scores and positive neighbourhood support (Fig. 7h). They included MIF–CD74^58,59^ and SPP1–CD44^60^, both previously linked to kidney injury, together with APP–CD74, ITGAX/ITGB2–ICAM1, BDNF–NTRK2 and SPP1–PTGER4. Receivers included SLC8A1-positive and C1Q-positive macrophages, cDC2s, atypical B cells, venous endothelium and thick ascending limb cells. In a MIF–CD74 example from plasma cells to cDC2s, one non-responder core contained 40 positive support edges to four cDC2 receivers. MIF and CD74 transcripts were detected at both ends of two of these edges, and one responder core contained no such edges (Fig. 7i).

Finally, we used CauTrigger to simulate attenuation of candidate drivers in the fitted model. Each target feature was capped at its training-set 10th percentile with all other inputs held fixed, and the resulting CauTrigger streamline plots were visualized in the causal latent space (Fig. 7j,k). After N9 occupancy was lowered, 67% of non-responders (6 of 9) showed local flow directions towards a responder-like state, whereas 91% of responders (10 of 11) retained a responder-like state (Fig. 7j). The predicted probability of non-response in non-responders fell by 0.025 on average (Fig. 7l). By contrast, lowering ACSS2, the top-ranked plasma-cell gene, directed only 33% of non-responders (3 of 9) towards a responder-like state, and responder flows pointed towards the non-responder side (Fig. 7k). Predicted probabilities changed by less than 10⁻⁵ in either group (Fig. 7l). Plasma-cell communication could not be perturbed, because no direction-concordant feature exceeded the attenuation threshold in any patient (Fig. 7l). This in silico analysis indicates that, within the learned model, a smaller N9 niche is associated with pathology states more similar to those of responders, thus offering a hypothesis for niche-directed modulation of treatment response.

### Supporting analyses of graft response

The complete supporting series contains the sensitivity networks, pathway and communication results of the main and alternative models, structure-unit trajectories across 25 anatomical and lesion scopes, lineage-associated genes and pathways, and clinical and cellular profiles. Each page and its Source Data state the cohort used: the treatment-response cohort, the creatinine-improvement cohort or an alternative PILOT-defined endpoint.

### Spatial niche features inferred from H&E sections predict kidney allograft outcomes

Niche extent and arrangement are properties of tissue architecture. Spatial transcriptomics, however, is not yet widely available and remains costly, which makes it unsuitable as a routine test in large clinical cohorts. We therefore asked whether spatial niches and their spatial relationships could be identified from routine H&E sections. We trained a multiscale deep-learning model with the nine niche labels and evaluated it by patient-grouped five-fold cross-validation on 10,139 H&E tiles from 49 CosMx-matched biopsies in 47 patients. The macro-averaged one-versus-rest AUROC was 0.914 (95% confidence interval 0.898–0.927), and the macro-F1 was 0.579 (0.537–0.606; Fig. 8a,b). Applied to an independent same-centre H&E validation cohort of 326 sections, the model registered H&E morphology and predicted niche identity at the same coordinates. Plasma-cell effector, remodelling and preserved parenchymal regions could then be followed across whole sections (Fig. 8c).

**Fig. 8.**
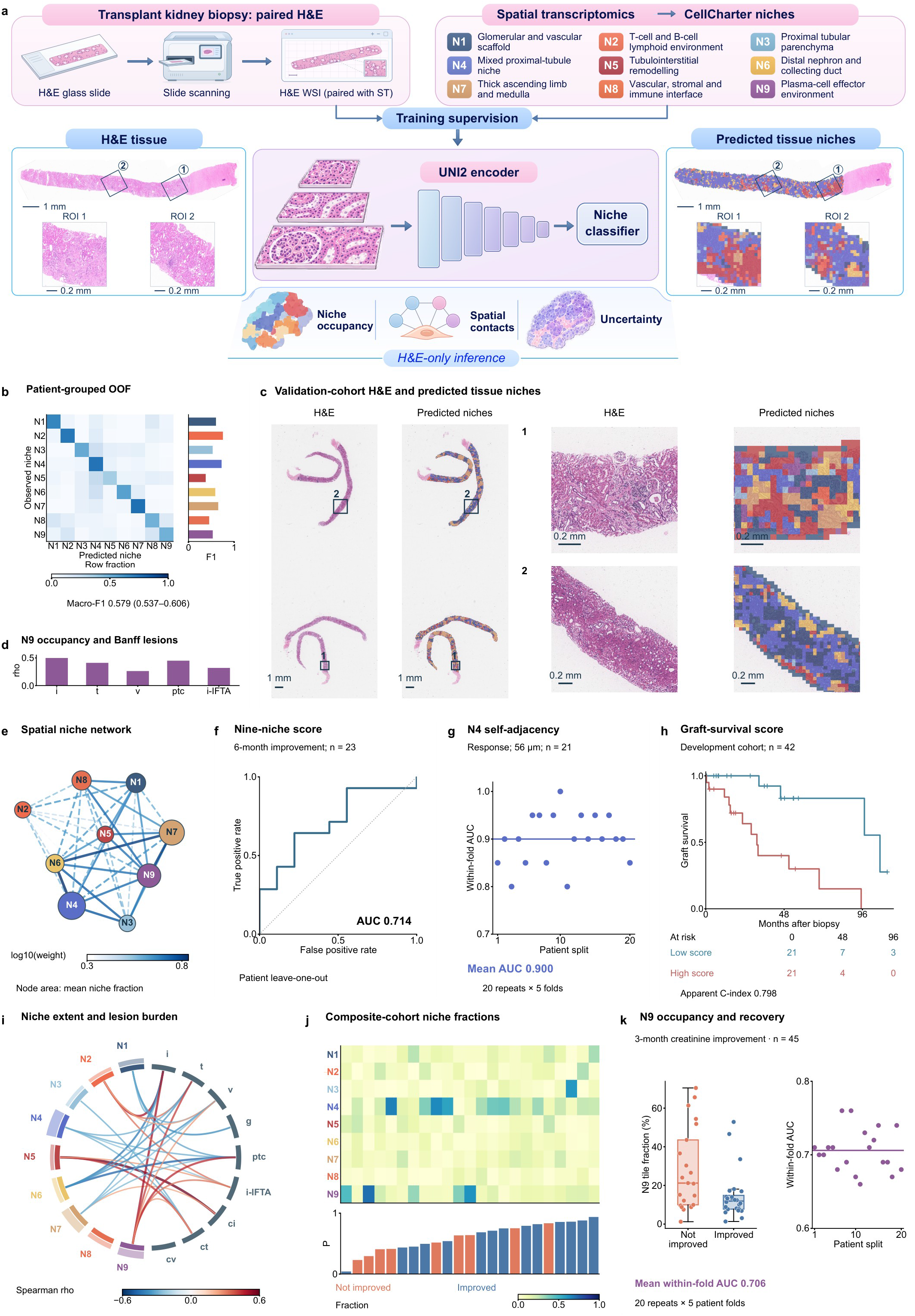
| Niche arrangement in routine H&E and post-biopsy functional outcomes. **a**, Spatial-niche supervision of multiscale H&E classification, with tissue and matched predictions from a representative biopsy. Scale bars, 1 mm in overviews and 0.2 mm in local views. **b**, Confusion matrix and per-niche F1 from five patient-grouped folds: 10,139 tiles from 49 biopsies in 47 patients. Patients receive equal weight and rows are normalized within observed classes. The macro-F1 interval is a patient-bootstrap 95% confidence interval. **c**, H&E, matched niche maps and local views from a second representative biopsy. Prediction fields have 56-μm sides; scale bars are indicated. **d**, Spearman rho for N9 occupancy versus Banff i, t, v, mononuclear ptc and i-IFTA; all five associations have joint FDR<0.05. **e**, Niche-distance network across the 326 sections of the validation cohort. Node colour identifies the niche and area represents mean occupancy. Edge weight is the inverse of the section-median bidirectional mean nearest distance for each niche pair (mm⁻¹). Colour indicates log10 weight and line width increases with weight; edges above the median weight are solid and the remainder are dashed. Node positions use a weighted spring layout. **f**, Patient leave-one-out ROC for a nine-niche composite predicting six-month creatinine improvement: 14 of 23 patients improved and nine did not. AUC, 0.714; fixed-prediction patient-bootstrap 95% interval, 0.471–0.921; Brier score, 0.226. **g**, Repeated five-fold performance of N4 self-adjacency z at 56 μm for the Fig. 7 response definition: 13 responders and eight non-responders. Each point is a repeat’s within-fold AUC weighted by positive–negative patient-pair counts; the line shows the mean of 20 repeats, 0.900. **h**, Kaplan–Meier curves in the 42-patient survival development cohort. The median training Cox score based on N5-to-N4 mean distance defines groups of 21; numbers at risk are given at 0, 48 and 96 months. There were 16 graft-loss events. C-index 0.798 is apparent development performance. **i**, Chord diagram linking nine niche fractions to nine Banff lesions. Outer bars show mean niche occupancy across 326 sections. Edges show associations with joint FDR<0.05; colour and width indicate the direction and magnitude of Spearman rho. The ptc field is the mononuclear score. Clinical correlations use one selected biopsy per patient. **j**, Niche fractions and held-out probabilities for the 23 patients in f, ordered by probability. Each column represents a patient. **k**, N9 tile fractions in patients with and without three-month creatinine improvement (left; 24 improved and 21 not improved). Each dot represents a patient. Boxes show medians and interquartile ranges; whiskers reach the most distant observations within 1.5 interquartile ranges. The right plot shows 20 repeats of five patient folds using N9 tile fraction as the single predictor. Each point is a repeat’s within-test-fold positive–negative pair-weighted AUC; the line marks the mean, 0.706. N9-to-N5 distance, N5–N9 adjacency and N9–eGFR associations are shown in Extended Data Fig. 10a–e. Composite-model preprocessing and tuning used training patients. The single feature in g was selected by repeated-split performance from 920 spatial features using outcomes in this cohort. The endpoint and cohort in f were chosen after comparing several endpoints. These internal estimates have distinct patients, endpoints and estimators, and probabilities are uncalibrated.

Niche extent in H&E-derived maps followed Banff lesions. With one biopsy per patient, N9 occupancy rose with interstitial inflammation (ρ = 0.498, q = 2.9 × 10⁻⁶), peritubular capillaritis (ρ = 0.450, q = 3.5 × 10⁻⁵) and tubulitis (ρ = 0.411, q = 7.3 × 10⁻⁵). Its associations with i-IFTA (ρ = 0.319, q = 0.0032) and intimal arteritis (ρ = 0.262, q = 0.021; Fig. 8d) were weaker. N9 occupancy also rose with tubular atrophy (ρ = 0.262, q = 0.022) and did not track interstitial fibrosis (ρ = 0.10). N5 occupancy instead rose with fibrosis (ρ = 0.588, q = 2.3 × 10⁻¹⁰), atrophy (ρ = 0.427) and i-IFTA (ρ = 0.255; Fig. 8i). N2 occupancy rose with interstitial inflammation (ρ = 0.491), tubulitis, intimal arteritis and i-IFTA, and N1, N4 and N6 declined with inflammatory or atrophic lesions. In the niche-distance network, N5 lay at the centre, close to most other niches, and N2 at the periphery (Fig. 8e). N9 therefore tracked inflammation in residual tissue, consistent with its extension into the residual interstitium of the spatial cohort (Fig. 2e,f), whereas N5 tracked scar extent.

We then asked whether H&E-derived maps carry information about subsequent function, defining improvement as a creatinine decrease of at least 10% from the pre-biopsy baseline. For six-month improvement, a composite of the nine niche fractions reached a patient leave-one-out AUC of 0.714 in 23 patients with complete follow-up, 14 of whom improved. Its 95% CI of 0.471–0.921 included 0.5 (Fig. 8f,j). N9 tile fraction alone predicted three-month improvement with a mean within-fold AUC of 0.706, and its regression coefficient was negative in every training fold (Fig. 8k). The spatial arrangement of niches also carried prognostic information. Responders had greater distances from N9 to N5 (mean within-fold AUC 0.833), and N5–N9 adjacency predicted three-month improvement (0.835; Extended Data Fig. 10a,b). In patient-level out-of-fold probabilities, patients who improved had a higher median nine-niche composite than those who did not (Extended Data Fig. 10c). The N9-to-N5 distance correlated with the twelve-month increase in eGFR (ρ = 0.383, joint FDR = 0.039, n = 101; Extended Data Fig. 10d,e). Homotypic adjacency of N9 and of N5 correlated with lower three-month eGFR (ρ = −0.361 and −0.363; FDR = 0.039 for both; n = 114; Extended Data Fig. 10d). These spatial relationships suggest a possible tissue mechanism. In grafts that did not recover, the plasma-cell effector and fibrotic remodelling regions each expanded into large contiguous areas that touched each other less than expected from their abundance (negative N5–N9 adjacency enrichment z-score). In grafts that recovered, N9 formed sparse foci lying farther from N5, with contacts close to random. Persistent sheets of plasma-cell effector tissue may continuously release immunoglobulin and inflammatory signals into adjacent residual tissue and thereby impede repair. This interpretation requires validation in independent longitudinal biopsy cohorts.

Arrangement of the mixed proximal-tubule niche related to treatment response and graft survival. Among 21 patients with response outcomes, N4 self-adjacency at 56 µm ranked first among 920 single spatial features, and responders had greater N4 self-adjacency (Fig. 8g and Extended Data Fig. 10c). In the 42-patient development cohort with 16 graft-loss events, a score based on the mean distance from N5 to N4 stratified graft survival, and the high-score group lost grafts earlier (apparent C-index 0.798; Fig. 8h). These results support the link between N9 and treatment response, and show that niche features defined by spatial omics can be read from routine H&E and related to outcome. The public atlas provides access to the underlying integrated single-cell resources (Extended Data Fig. 10f).

## Discussion

The SPECTRA atlas links the cellular content of kidney allograft biopsies to function after treatment. Of the nine niches defined in 83 biopsies, the plasma-cell effector niche was the only one whose extent tracked eGFR recovery. Its plasma cells carried a terminally differentiated secretory state, and the same niche dominated the ecological sensitivities of a multilevel model of non-response. Beyond this niche, TLS retained TCF7-high T cells beside CCL19-expressing fibroblasts, acute lesions engaged each compartment through its own endothelial or tubular programme, and chronic lesions combined plasma-cell secretory adaptation with loss of tubular fatty-acid oxidation. In mice, interferon-activated myeloid states accompanied persistent tubular injury along the rejection path, and tubular metabolic recovery accompanied tolerance.

Plasma-cell-rich rejection has long been associated with poor graft survival^17–19^. Our data place these cells in a niche of defined composition and position. N9 extended into unscarred and scarred residual interstitium, its fibroblasts expressed CXCL12 and C7, and its plasma cells expressed more IGHG1, IGKC and TXNDC5 and less PAX5 and proliferation-associated genes than plasma cells elsewhere. In scarred interstitium, plasma cells raised XBP1, HSPA5 and SEC61A1 as fibrosis advanced, the secretory programme that XBP1 drives during terminal differentiation^46^. CXCL12-expressing stromal cells retain long-lived plasma cells in bone marrow^32,33^, and in rejecting lung allografts IL-6 trans-signalling regulates stromal CXCL12 expression and the plasma-cell niche^61^. N9 may provide a comparable stromal survival environment within the kidney graft. Its plasma cells also expressed more TXNDC5, a disulfide isomerase that supports protein folding in the endoplasmic reticulum and promotes fibrosis in kidney fibroblasts^57^. In H&E-derived maps, N9 tracked interstitial inflammation, tubulitis, arteritis, capillaritis and i-IFTA, whereas N5 tracked scar extent. N9 therefore marks inflamed tissue that has not yet been replaced by scar. CD38-directed antibodies reduce microvascular inflammation and molecular rejection activity in AMR^20,21^ and have been used in chronic active disease^22^. This niche therefore provides a possible biological mechanism and an observable histological marker for evaluating plasma-cell-directed treatments.

TLS in the graft retained a TCF7-high stem-like T-cell state. Similar observations have been made in other diseases. TCF1-high CD4+ T cells in perivascular TLS sustain autoimmune vasculitis^62^, TCF1-high CD8+ T cells provide the proliferative burst after PD-1 blockade^63^, and stem-immunity hubs in lung cancer place TCF7-positive CD8+ T cells beside CCR7-positive dendritic cells^64^. In allografts, TCF7 was higher in T cells inside TLS, naive-like subtypes concentrated at the centre, and fibroblast CCL19 with T-cell CCR7 formed the strongest stromal homing signal. CCL19-expressing fibroblasts also organize TLS in aged kidneys^65^. Fibroblasts in kidney allografts may thus recruit stem-like T cells through chemokines and support TLS formation, and under systemic immunosuppression these structures may serve as a local immune reservoir for rejection. Stage II structures predominated in AMR/MVI, and Stage I fibroblasts raised CCL21 and C3. Advanced TLS in protocol biopsies have been associated with progressive graft dysfunction^66^; given the size of our cohort, we did not test the relationship between TLS stage and outcome. The TLS signature also declined with time in mouse grafts and showed no time association in human samples. Tolerant mouse grafts form regulatory lymphoid structures^28^, so the effect of TLS on graft outcome is likely to depend on the cell states they hold. Bulk and even single-cell RNA sequencing may fail to capture TLS-associated signals accurately, which underlines the value of spatial analysis.

Resolving lesions by compartment changes how several Banff findings are read. Peritubular endothelium raised interferon-γ-associated antigen presentation, including HLA-F, which marks poor prognosis in AMR^39^, whereas glomerular endothelium reduced MHC expression as glomerular inflammation rose. We propose two possible explanations for the latter. First, MHC downregulation may reflect an accommodation-like state. Accommodation describes a graft that keeps functioning despite humoral immunity directed against it^67^, and reduced antigen presentation could make glomerular endothelium a less visible immune target. Second, it may reflect endothelial dedifferentiation or endothelial-to-mesenchymal transition, whose markers are expressed by peritubular-capillary endothelium in AMR^68^. Distinguishing these possibilities will require endothelial identity markers and longitudinal biopsies. In microvascular lesions, endothelial fractions rose in the peritubular-capillary region and immune cells lay closer to peritubular endothelium. Banff 2022 recognizes double-negative MVI as a distinct category^1^ with outcomes similar to those of AMR^7^, and some cases have been linked to missing-self recognition by natural killer cells^69,70^. Here, double-negative MVI shared with active AMR the approach of immune cells to peritubular endothelium and endothelial interferon-γ and antigen-processing programmes, and its natural killer-cell fraction was also raised. This is consistent with missing-self-driven endothelial injury and suggests that similar effector mechanisms may act on the microvasculature even without detectable antibody. Isolated intimal arteritis, whose status as rejection has been debated^3,42,43^, coincided with a failed-repair-associated tubular programme even though immune cells did not approach tubules. A multicentre study found that the v-lesion itself did not affect graft outcomes and questioned the place of isolated v in rejection diagnosis^44^. Isolated v may also reflect non-immune injury such as ischaemia–reperfusion^71^, and a tubular injury programme that arises without immune contact is consistent with this view. In our data, immune cells in isolated-v cores avoided tubules more than in Control, yet proximal tubules expressed failed-repair genes such as HAVCR1 and DCDC2. Some isolated v may therefore represent a state dominated by parenchymal injury rather than typical cellular rejection, a possibility that requires validation in larger cohorts. In chronic lesions, atrophic tubules lost fatty-acid oxidation, a metabolic defect that promotes kidney fibrosis^47^ and that resembles the metabolic shutdown of structural cells in rejection^10^. This suggests that, beyond immunosuppression, metabolic protection of parenchymal cells may also be important. Plasma cells, T cells and macrophages in scarred interstitium retained secretory and interferon programmes, in line with the poor prognosis of inflammation in scarred areas^5,6^. This further supports attention to plasma-cell-directed treatment.

In mice, graft trajectories connected myeloid activation with tubular injury and tubular metabolism with acceptance. Along the rejection path, classical monocytes, cDC1s and MGL2 macrophages raised interferon and complement programmes, and failed-repair proximal tubules expanded, as described after acute kidney injury^24,25^ and in mouse TCMR^26^. Along the tolerance path, failed-repair tubules lost *Havcr1*, *Vcam1* and *Lcn2*, and differentiated proximal tubules regained *Hnf4a*– and *Ppara*-associated transport and fatty-acid metabolism. DBA/2J grafts are accepted despite intragraft T-cell infiltration^29,30^, and our data show that acceptance is accompanied by recovery of the tubular programme that atrophic human tubules lose. Drug-target ranking nominated xanthine oxidase inhibitors against the rejection path, consistent with their protection against oxidative ischaemia– reperfusion injury^54^, and PPARA agonism for the tolerance path. The effects of these targets remain to be tested in animal experiments.

Niche extent and arrangement are features of tissue architecture, which allowed a model trained on spatial labels to read them from routine H&E sections. This step extends spatial atlases^13,15,72^ to standard pathology material. In H&E-derived maps, N9 extent, its separation from N5 and the self-adjacency of N9 and N5 related to subsequent kidney function. The public single-cell part of SPECTRA can be queried through a web portal (Extended Data Fig. 10f). The H&E validation cohort, however, comes from a single-centre observational study, so independent multicentre cohorts are needed to validate the spatial niche features more broadly.

In summary, a plasma-cell effector niche in residual and scarred interstitium tracks non-recovery after kidney allograft rejection. It sits alongside compartment-specific endothelial and tubular responses to acute lesions and a tubular metabolic programme whose recovery accompanies tolerance in mice. Reading these niches from routine H&E offers a route to bring them into clinical biopsy assessment.

## Methods

### Human biopsy cohorts and clinical endpoints

The spatial cohort comprised 83 kidney-allograft biopsies represented by 84 tissue cores. The study was approved by the Ethics Committee of Zhongshan Hospital, Fudan University (B2026-459R), which waived informed consent. Diagnoses were harmonized into eight groups: reference (n = 7), borderline changes (9), T cell-mediated rejection (TCMR; 14) and antibody-mediated rejection or microvascular inflammation (AMR/MVI; 18). The remaining groups were mixed rejection (20), BK polyomavirus-associated disease (10), calcineurin-inhibitor toxicity (CNI; 4) and C4d deposition without rejection (1). The reference group contained six implantation biopsies and one normal post-transplant biopsy. Banff lesion scores were taken from the pathology reports^1^, and BK viraemia, polyomavirus nephropathy and coexisting rejection were recorded as separate fields. Fifteen biopsies had DSA-negative/C4d-negative MVI.

The human single-cell reference contained 352,911 cells or nuclei from 120 libraries, 117 sampling units and 12 public projects^8,9,26,73–81^. Whole-tissue analyses used 310,779 profiles from 113 libraries combined into 110 sampling units; seven immune-sorted units entered only the cell-state analyses. Diagnosis-based analyses used the 94 units (97 libraries; 275,474 profiles) with a verifiable diagnosis. In these analyses, Control comprised 13 normal or nonspecific post-transplant units and 22 native-kidney units.

Functional endpoints were defined relative to the clinical index biopsy and evaluated in the patients who had the corresponding measurements. For niche–function associations (Fig. 2g), implantation biopsies were excluded and the first eligible biopsy of each known patient was used, giving 48 index biopsies. Relative eGFR change was (median eGFR on days 60– 120 − baseline)/baseline, with the median on days −14 to +7 as baseline and the eGFR recorded with the biopsy as fallback; 35 patients had this endpoint. The functional-response cohort of Fig. 7 comprised 20 recipients, each with one index biopsy, classified as 11 responders and nine non-responders. Baseline was the median on days −14 to +7, and follow-up comprised measurements on days 30–180; an assay was eligible with at least two follow-up measurement days spanning at least 28 days. Response required an eGFR increase of at least 10% and a creatinine decrease of at least 10% when both assays were eligible, or the corresponding threshold for the single eligible assay. The primary diagnosis, taken from the first component of composite diagnoses, was active, chronic, chronic active or probable AMR; acute TCMR grade IB, IIA or IIB; DSA-negative/C4d-negative MVI; borderline changes; and C4d staining without rejection. Unclassified outcomes were treated as missing. Endpoints for the H&E cohort are described with the H&E analyses.

### Spatial molecular imaging, registration and segmentation

The human spatial cohort used archival formalin-fixed, paraffin-embedded (FFPE) allograft needle biopsies from the biobank of Zhongshan Hospital, Fudan University. Eighty-four tissue cores were arranged on two slides (38 and 46 cores) and sectioned at 4 µm. Each section was first assayed for 64 protein features (CosMx Human Immuno-Oncology Protein Panel; 62 targets plus mouse IgG1 and rabbit IgG isotype controls) and then for 18,934 RNA targets (CosMx Human WTX Panel)^16^. Morphology channels were PanCK, CD68, CD45, a membrane marker and DAPI. The CosMx SMI instrument (Control Center 2.0.0.24) acquired 4,256 × 4,256-pixel fields (120.28 nm per pixel; about 0.51 mm) as eight z-steps of 0.8 µm. The two slides yielded 800 fields (439 and 361), with 3–22 fields per biopsy (median 8). After the run, the same section was stained with H&E and scanned on a Vectra Polaris at 20× magnification.

RNA and protein cell segmentations were registered at cell level in 795 fields. H&E images were aligned to CosMx coordinates by tissue outline and then refined field by field with nuclear point sets. After refinement, median nearest-neighbour distances between CosMx and H&E nuclei were 0.948 and 1.055 µm on the two slides, and 756 fields reached cell-level registration. Anatomical compartments were inferred on the registered H&E by several deep-learning structure-segmentation models that we trained on H&E images annotated by pathologists at our centre. Pathologists reviewed and corrected each result in QuPath^38^, and the compartments were then mapped to each CosMx cell by coordinate. They comprised glomeruli, tubules, arteries, veins, peritubular capillaries (PTC), interstitium, interstitial fibrosis and tubular atrophy (IFTA) and tertiary lymphoid structures (TLS). PTC and glomerular capillary loops were segmented by the models described below, and TLS were outlined separately by a pathologist.

PTC and glomerular capillary loops were segmented in a shared frame of 0.5 µm per pixel from the CD34 protein channel and the registered H&E of the same section. Several pathologists drew training instances manually in QuPath^38^. The PTC training set contained 174 instances in ten fields from eight cases. A residual U-Net^82^ with group normalization^83^ and 16 base channels received CD34 and H&E RGB as four input channels and predicted foreground and contour maps. Training used 256 × 256-pixel crops, batches of four, random 90° rotations and BF16 mixed precision, with three-quarters of crops centred near a true boundary between adjacent capillaries. The foreground loss was 0.30 × binary cross-entropy plus 0.70 × Tversky loss^84^, with false-positive and false-negative weights of 0.30 and 0.70. The contour loss combined 0.20 × binary cross-entropy, 0.25 × soft Dice and 0.55 × Tversky loss (weights 0.20 and 0.80), and the total loss added 1.5 times the contour loss to the foreground loss. AdamW used a learning rate of 3 × 10⁻⁴ and weight decay of 10⁻⁴, and the learning rate was halved on validation plateaus (patience eight epochs; floor 10⁻⁶). Seeds from the foreground minus contours were expanded over the full foreground by watershed. Thresholds were 0.65 for foreground and 0.45 for contours, minimum seed and instance sizes were 16 and three pixels, and holes up to 256 pixels were filled. Post-processing parameters were selected on pooled out-of-fold predictions, and the final model was trained for 22 epochs. Random seeds were 20260822 plus a fixed offset for each fold and for the final model.

The glomerular-loop training set contained 1,112 instances in 13 multimodal-stained fields from 13 cases, comprising five 256 × 256-pixel and eight 512 × 512-pixel crops. Cellpose-SAM^85^ was fine-tuned for 30 epochs on CD34 and the haematoxylin and eosin components of H&E, with a learning rate of 10⁻⁵, weight decay of 0.1 and a scale-augmentation range of 0.5. Random seeds were 20260824 plus a fixed offset for each fold and for the final model. Inference used no diameter rescaling, a tile overlap of 0.25, a cell-probability threshold of −0.5, a flow threshold of 0.6, a minimum instance of five pixels, a maximum area fraction of 0.05 and 150 dynamics iterations. When Cellpose-SAM returned fewer than 40% of the instances found by a distance-map U-Net, the U-Net output was used.

Both models were evaluated by five-fold cross-validation grouped by case, with all fields of a case in one fold (Supplementary Figs. 7 and 12). Out-of-fold semantic Dice, instance F1 at IoU 0.5, panoptic quality (PQ) and absolute relative count error were 0.875, 0.753, 0.581 and 0.150 for PTC. For glomerular loops they were 0.759, 0.453, 0.303 and 0.176. Pathologists reviewed all deployed masks in QuPath 0.7.0, accepting, reshaping or deleting model structures and adding missed ones. Models were trained on one RTX 5090 GPU. PTC masks include the wall and enclosed lumen. Whole glomeruli and their capillary loops are separate objects. Overlapping PTC masks were resolved within slide and core by centroid assignment to the smallest mask, with ties broken by identifier.

Native CosMx segmentation yielded 1,256,052 cells, and cell quality control was performed within each slide. The lower RNA-count limit was set per core. For shallowly sequenced cores (median count below 250), it was the smaller of 200 and the 10th percentile of the core’s counts; for the other cores, it was the slide mean of log10 counts minus 2.5 standard deviations. All cells also required at least 20 counts and at least 10 detected genes, and neither counts nor detected genes could exceed the slide log10 mean plus 2.5 standard deviations. Cell fragments split by field boundaries in the vendor segmentation (split fraction below 0.5) were then removed, as were cells with a local signal-to-background ratio below 1. This ratio divided counts smoothed with a Gaussian kernel of 10-µm bandwidth (truncated at 80 µm) by the mean negative-probe count. Field-level quality control flagged fields without removing cells. Pixel size was corrected to the measured 0.12143 µm, multiplying lengths and areas by 1.0096 and 1.0192. Within overlaps between adjacent fields, cell pairs with centroids less than 3 µm apart or a polygon intersection over union of at least 0.15 (search radius 12 µm) were treated as duplicates. The cell with more counts was kept. After removal of tissue folds annotated manually on H&E, 1,133,890 cells (90.27%) remained, 653,588 of 716,353 and 480,302 of 539,699 on the two slides. Transcripts spilling over from adjacent cells were removed with cellAdmix^86^. This method fits ten non-negative matrix factorization factors to the 2,000 genes with the highest counts and identifies contaminating transcripts within each core with a conditional random field. Transcripts that FastReseg^87^ (version 1.1.2; likelihood-ratio threshold 5) flagged for trimming were then subtracted gene by gene from the decontaminated counts, and cells flagged as merged or anomalous were removed, leaving 1,133,651 cells. Cells with an ovrlpy^88^ (version 1.2.0) vertical signal integrity below 0.7 were removed, and cells without an estimate were kept. The annotated object contained 1,011,984 cells, 597,383 and 414,601 on the two slides.

### Cell annotation and cross-dataset integration

The human single-cell reference combined manual and KPMP labels^23^ with six confidence-weighted voting sources, giving four compartments, 32 lineages and 148 states. Ambiguous clusters were resolved by cluster-level majority vote modelled on CellVote in OmicVerse^89^, and a popV^90^ majority vote over six project predictions served as a comparison (Supplementary Fig. 10). High-confidence composition analyses applied additional identity-quality filters; fractions within a parent lineage used all cells of that parent.

CosMx cells were annotated in a nested hierarchy (Extended Data Fig. 1a and Supplementary Fig. 1 and Supplementary Source Data S78–S93) of four compartments, 26 level-1 lineages, 65 level-2 cell states and 114 level-3 identities. Lineage identity was assigned by a weighted logarithmic opinion pool of eight expert models: RCTD^91^, TACCO^92^ and its depth-corrected centred-log-ratio variant, cell2location^93^, hierarchical InSituType^94^ with biopsy reference profiles from the human atlas, Spotiphy^95^, TACIT^96^ and scANVI^97^. Experts were admitted on the effective number of classes in their calls; HierAnnot, the ninth candidate, was excluded because one class dominated its calls. Expert posteriors were calibrated with depth-dependent confusion estimated by paired downsampling of the same cells, and pool weights reflected spatial compliance and the retention of each label across sequencing depths. Protein-assisted HieraType^98^ calls and selected protein channels defined 217,865 anchor cells (21.7%) that constrained compartment membership, and anatomical rules for glomeruli, cortex and large vessels were applied after pooling. For each cell, candidate labels were added in decreasing posterior order until their probabilities summed to 0.90, forming a prediction set. Cells with up to six candidates kept their lineage identity, and the remainder received a compartment identity only.

Cell states were assigned within each parent lineage. TACCO, depth-corrected TACCO, cell2location and RCTD used a human single-cell reference of 82,620 cells in 148 classes. HierAnnot ran on a kidney-transplant hierarchy, TACIT used marker signatures, and scANVI and a classifier on a ResolVI latent representation (random seed 0) were trained with labels from 479,169 high-confidence cells. Arms whose posteriors had a cosine similarity above 0.85 were treated as redundant. Raw TACCO (0.918 with depth-corrected TACCO) was therefore removed, and all pairwise cosines among the remaining seven arms were below 0.50. Arm weights were estimated within each parent lineage, each cell combined only its informative arms, and conformal prediction (α = 0.10) gave candidate sets within the parent.

Fig. 1d shows the first marker gene of each lineage, scaled 0–1 per gene. Spatial protein intensities were transformed as arcsinh(intensity/35), with raw values of 5 or less set to zero (Supplementary Source Data S253). The two cores of one biopsy were pooled by cell count, cell-type means were averaged with equal biopsy weight, and each protein was scaled 0–1 for display.

Harmony^99^ integrated each data system with slide and biopsy as covariates for CosMx, sample and project identifiers for human single-cell data, and sample identifier for mouse data; mouse clustering used Leiden^100^ on the integrated graph. ResolVI^101^ was trained semi-supervised. Coarse scANVI calls in nine classes labelled 954,408 cells with confidence of at least 0.7, and the remaining 57,576 cells were unlabelled. Slide was the batch key, spatial neighbourhoods comprised the ten nearest cells, and ribosomal, haemoglobin and control probes were removed. Training used batches of 512 for 50 epochs and produced a ten-dimensional latent representation. The two-dimensional CosMx views in Fig. 1b and Extended Data Fig. 1b use the UMAP of this representation, with 15 neighbours, cosine distance, minimum distance 0.02, spread = 2 and random seed 0. All 1,011,984 cells are matched by cell ID to the final cell identities and slide labels. Extended Data Fig. 1b shows the same cells before and after integration with equal axis scaling. It contains 1,011,984 CosMx cells coloured by slide, 275,474 human profiles coloured by the 97 library identifiers of the 94 diagnosis-eligible sampling units, and mouse cells coloured by animal. Extended Data Fig. 1c shows principal-component coordinates of sample-level expression for 83 CosMx biopsies, 97 human libraries and the mouse samples.

### Mouse kidney transplantation and single-cell RNA sequencing

The main mouse cohort comprised 34 C57BL/6 recipients of syngeneic (C57BL/6), BALB/c, DBA/2J or DBA/2N kidney grafts, with tacrolimus treatment and donor presensitization as additional arms. Both DBA substrains carry a C5-deficient background. DBA/2J grafts were classified as tolerant. In BALB/c grafts, postoperative day (POD) 7 represents acute rejection, and POD28 and POD90 represent mixed rejection; the four POD28 grafts comprised two with severe and two with mild rejection. At POD7, groups contained two syngeneic, two BALB/c, three tacrolimus-treated BALB/c and two presensitized BALB/c grafts. Later groups contained four BALB/c grafts at POD28 and two at POD90, and two presensitized BALB/c grafts at POD28. DBA/2J groups contained three, four and two grafts at POD7, POD28 and POD90, and DBA/2N groups two and six grafts at POD7 and POD28. Each timepoint used different animals. Nephrectomy history and donor–recipient assignments were recorded per animal. The mild-versus-POD7 and severe-versus-POD7 contrasts share the two POD7 animals and do not test a severity-by-time interaction.

Mouse kidney transplantation followed a published microsurgical technique^102^. Donors and recipients were male mice aged 8–12 weeks from commercial suppliers, and surgery was performed under isoflurane inhalation anaesthesia. The donor kidney was procured en bloc with an aortic cuff and the renal vein, which were anastomosed end to side to the recipient infrarenal aorta and inferior vena cava. The ureter was anastomosed to the bladder. One native kidney was removed at transplantation, and analgesia was given after surgery. Recipients observed beyond POD7 underwent contralateral native nephrectomy on POD7, leaving the graft as the only life-supporting kidney. Tacrolimus-treated recipients received 1 mg kg⁻¹ tacrolimus subcutaneously each day from transplantation until harvest. Presensitized recipients received a BALB/c donor skin graft 7 days before kidney transplantation. Animal experiments were approved by the institutional animal ethics committee.

Grafts were perfused before harvest. Tissue for single-cell sequencing was placed in preservation solution. Tissue for histology was fixed in paraformaldehyde, embedded in paraffin and stained with H&E. Sections were scanned on a 3DHISTECH Pannoramic scanner at 20× magnification (0.2738 µm per pixel), and several pathologists blinded to group graded lesions by Banff criteria. Tissue was digested with collagenase I/II (1 mg ml⁻¹) and neutral protease (0.5 mg ml⁻¹) at 37 °C for 30 min and passed through 70-µm and 30-µm strainers. Red cells (Miltenyi 130-094-183) and dead cells (Miltenyi 130-090-101) were removed, and cells were counted with AO/PI and adjusted to 700–1,200 cells µl⁻¹. Libraries were built with 10x Genomics GEM-X Single Cell 3′ v4 reagents (kit PN-1000691, chip PN-1000690, user guide CG000732). They were sequenced on a NovaSeq 6000 to a target of at least 20,000 reads per cell, and alignment used 28 bp of read 1 and 91 bp of read 2. Reads were aligned to GRCm39 with Cell Ranger 9.0.1 or 10.0.0 including intronic reads, and CellBender^103^ (150 epochs; false-positive rate 0.01) removed ambient RNA. Cells were retained with at least 200 detected genes and at most 10% mitochondrial counts. Total counts and detected genes on a log scale, and the fraction of counts in the 20 most abundant genes, had to lie within five median absolute deviations of the sample median.

Scrublet^104^ (expected doublet rate 0.03) and scDblFinder^105^ (fixed doublet rate 0.004 with the automatic threshold) removed doublets. Souporcell^106^ genotype clustering assigned donor or recipient origin and identified inter-genotype doublets where strains differed, and scalpi^107^ recovered neutrophils. After Harmony integration^99^ and Leiden clustering^100^, cells were annotated hierarchically into 79 level-2 cell states. The atlas contains 442,320 cells.

### Cell composition

Lineage fractions used all captured cells of a sampling unit or animal, and state fractions the complete parent lineage. States absent from an observed parent were counted as zero, and fractions with an absent parent were undefined. Regional fractions used all cells in the biopsy compartment, and niche occupancy all cells of the biopsy.

Fig. 1e shows T-cell, B-cell, monocyte/macrophage, plasma-cell, fibroblast and endothelial fractions in mice, 94 human sampling units and 83 CosMx biopsies. The myeloid-to-lymphoid ratio divided the summed fractions of monocytes/macrophages, dendritic cells, plasmacytoid dendritic cells (pDC), neutrophils and mast cells by those of T, natural killer (NK), B and plasma cells, without a pseudocount. Mouse numerators used monocytes/macrophages, dendritic cells, neutrophils and basophils. Units with a zero denominator were undefined. Group inference used CosMx biopsies. Lineage fractions were truncated to [10⁻⁴, 1 − 10⁻⁴], logit-transformed and analysed with limma^108^ using robust empirical-Bayes moderation, with diagnosis, standardized sampling year and standardized RNA depth as covariates and BH correction over 154 diagnosis coefficients. The log2 myeloid-to-lymphoid ratio was analysed by ordinary least squares with the same covariates and BH correction over seven coefficients. Human single-cell data are shown as sampling-unit distributions.

Fig. 1f shows biopsy-mean composition within glomeruli, arteries, PTC, tubules, interstitium and scarred interstitium. Glomeruli, arteries and tubules used histological cell membership, PTC used the 113,564-cell PTC assignment, and unscarred and scarred residual interstitium used their mapped geometries; biopsies lacking a region contributed no value. The microvascular three-group comparison includes seven controls, nine biopsies with MVI negative for both donor-specific antibodies and C4d without concurrent TCMR or borderline changes, and four with active AMR. MVI with TCMR or borderline changes, BK polyomavirus nephropathy and probable AMR are excluded. Each biopsy-compartment cell fraction is one observation, with all cells of the corresponding biopsy compartment as the denominator. Fractions are clipped at ε = 10⁻⁴ and transformed by the natural-log logit; robust empirical-Bayes limma models include diagnostic group, standardised sampling year and standardised capture depth. Three pairwise contrasts cover 21 glomerular and 23 peritubular-capillary cell fractions, the latter including aggregate endothelium, with BH correction over all 132 tests. Both compartments contain all 20 biopsies, and the design has rank five and 15 residual degrees of freedom. Tubular T-cell fractions are shown by Banff t grade, and scarred-interstitial plasma-cell and fibroblast fractions by i-IFTA grade. Supplementary Source Data S248–S249 show immune composition in biopsy strata of local load. Strata were quartiles of interstitial, arterial, glomerular and PTC load, inflammation tiers for tubules, within-core quartiles for fibrosis and intimal-area cut points of 65.6057%, 83.6932% and 91.6444%. Biopsies were weighted equally and could enter several strata.

Mouse group comparisons used exact two-sided Mann–Whitney tests that enumerated every group-size-preserving assignment of animals, with average ties; the smallest attainable P with two animals per arm was 1/3. Separate BH families contained 416 lineage tests, 1,264 planned state tests (1,226 estimable) and 1,200 programme tests. Milo^109^ tested animal-level neighbourhood abundance with robust edgeR quasi-likelihood^110^ and graph-weighted spatial FDR.

### Spatial niches

Niches were identified with CellCharter^31^ (0.3.7) on scVI^111^ embeddings. scVI was trained in scvi-tools^112^ 1.5.0.post1 on 4,000 variable genes selected per slide using the Seurat v3 method, with a zero-inflated negative binomial likelihood and ten latent dimensions. Slide was the batch key and biopsy a categorical covariate, and training used batches of 256 with the default epoch number, early stopping and seed 42. Within-core Delaunay graphs were pruned of edges longer than the 99th length percentile or 100 µm, and each cell’s features were concatenated with one-to three-hop neighbourhood aggregates (40 dimensions).

Gaussian mixtures with K = 3–15 were fitted ten times each, and stability and pathology-associated structure supported K = 9 (Fig. 2a). The two cores of one biopsy were combined for biopsy summaries. Niche representations from Novae^113^, Nicheformer^114^ and COVET/ENVI^115^ were compared with the CellCharter solution (Supplementary Fig. 15 and Supplementary Source Data S125–S126). Fig. 2b shows the log2 fold enrichment of each lineage within each niche.

Niche fractions (Fig. 2d) were analysed with logit-limma (ε = 10⁻⁴; design including diagnosis, standardized sampling year and standardized RNA depth) and with sccomp^116^, which reports posterior FDR. Logit-limma results were BH-corrected across the nine niches for each diagnosis coefficient. Fig. 2d shows the coefficients on a colour scale saturating at ±2 with unadjusted model P values; q values are in Source Data. Pairwise diagnosis comparisons were corrected across 15 diagnosis pairs × nine niches. Fig. 2f shows niche fractions in 83 biopsies and in unscarred and scarred residual interstitium from 82 and 81 biopsies, giving 249 biopsy–niche and 489 region–niche observations; cores of one biopsy were pooled by cell count. Extended Data Fig. 2a shows all 84 cores, with registered segmentation polygons coloured by niche at 8 µm per mosaic pixel; 1,011,981 of 1,011,984 cells had a drawable polygon. Extended Data Fig. 2b,c shows niche fractions across the eight diagnosis groups and the 15 recorded rejection subtypes.

Niche neighbourhood enrichment (Fig. 2c) was the observed-minus-expected adjacency between niches. Diagnosis labels were permuted 100 times between biopsies, P values used an add-one correction, and BH correction covered 72 directed non-self pairs across three contrasts. The rejection-minus-Control matrix is the difference between the two group matrices. A recovery contrast used 12 and nine cores with 10,000 permutations.

Anatomical niche composition (Fig. 2e) averaged within-biopsy niche fractions in each compartment, each with its own cell denominator; the i-IFTA bar uses scarred residual interstitium from 54 biopsies with Banff i-IFTA > 0. Anatomical enrichment compared a niche’s compartment fractions with whole-core compartment fractions in cores with at least 20 niche cells, using Wilcoxon tests with BH correction over 72 niche–compartment comparisons; niche–Banff Spearman correlations formed a family of 90 tests. Annular composition used rings of 0–20, 20–40, 40–60 and 60–80 µm, excluding centres with an empty annulus and the 5% of centres with the lowest local density. BallTree distances averaged up to ten nearest hub cells per partner before biopsy medians were taken, with BH families defined by contrast, annotation level and hub (Supplementary Source Data S127 and S282–S283).

### Niche occupancy and kidney function

In the 48 index biopsies, the occupancy of each of the nine niches was tested against relative eGFR change in the 35 patients with this endpoint (Fig. 2g). Each variable was rank-transformed and residualized on standardized year and depth by least squares, and Spearman correlation was computed on complete residual pairs. BH correction covered the nine niches within each metric, endpoint, adjustment and indexing rule. Fig. 2g reports the adjusted estimates; for N9, the unadjusted Spearman ρ was −0.586 (P = 0.000219; q = 0.00197). The ≥1-year absolute eGFR endpoint also included 35 patients, and density endpoints included 20 (Source Data).

### Within-lineage spatial differential expression

smiDE 0.0.2.5^117^ compared a fixed lineage between niches, anatomical states or diagnoses. Each level required at least three cells per core, and nine negative-probe features were excluded. Standard negative-binomial models adjusted for rank-normalized expression of the same gene in neighbouring cells of other types within 50 µm, with the logarithm of the cell’s RNA count as offset. Genes with Standard q < 0.05 and |log2FC| ≥ log2(1.5) were re-estimated by Optimal models that added a biopsy-nested Matérn Gaussian-process spatial effect. Testing families comprised the genes estimated for each label and contrast. Per-core effects were combined within each biopsy and then across biopsies by restricted maximum likelihood (REML) random-effects meta-analysis in metafor^118^, or by fixed effects when two units contributed. Standard estimates replaced spatial fits that failed. Proximal tubule (PT) and thick ascending limb (TAL) analyses used pooled Optimal coefficients. Rejection comparisons within a lineage and niche required at least three cells per unit and three units per group, with BH correction within niche across prespecified candidates. Fibroblast regional programme models additionally adjusted for subtype composition and used whole-recipient bootstrap intervals.

Fig. 2h shows 60 niche-associated genes in ten cell classes. Thirty-two genes, eight each from T cells, B cells, monocytes/macrophages and fibroblasts, represent receptor signalling, antigen presentation, myeloid response and stromal chemokine and matrix states. The other 28 come from NK cells, dendritic cells, pDC and peritubular/venous, arterial and glomerular endothelium. Each displayed gene has at least one positive niche-versus-rest effect with q < 0.05 within its class, excluding estimates with |log2FC| ≥ 20. Colour shows the smiDE/metafor log2FC, saturating at ±2.

For niche over-representation analysis (Supplementary Source Data S280–S281), genes with positive log2FC and FDR < 0.05 in each class–niche model were tested against all genes with finite estimates in the same model by one-sided hypergeometric tests. Libraries were GO biological process, cellular component and molecular function 2026, Hallmark 2026.1, KEGG 2026 and Reactome 2022. All gene sets intersecting the background were included, with zero-overlap sets at P = 1, and BH correction was joint across the six libraries within each class–niche family. Of 580,377 tests in 213 families, 399 had q < 0.05.

### Niche ligand–receptor expression and receiver scores

Fig. 2i compares ligand–receptor expression products between a niche and the rest of the same core, using LIANA^119^ consensus pairs. Expression was normalized to ln(1 + CP10k) per cell and averaged within core, cell class and region, and multisubunit ligands or receptors used the minimum subunit mean. Each ordered pair included at least one immune class. Each class required at least three cells per region, and each molecular component a mean of at least 0.02 in both regions. The core statistic was log2[(L × R + 10⁻⁶) in the niche/(L × R + 10⁻⁶) in the rest], clipped to ±10, where L and R are the ligand and receptor means of the sending and receiving classes. Pairs with at least six paired cores were tested with two-sided one-sample t-tests, and BH correction covered testable pairs within each niche. Displayed associations required q < 0.05, a mean log2 ratio of at least 0.5 and at least 12 paired cores, with orientation matching CellChatDB^34^.

Receiver scores from CytoSignal^120^ integrated ligand fields within 200 µm for diffusion signals or over Delaunay neighbours for contact signals. For Supplementary Source Data S322, scores were compared between each niche and the rest of the core within each receiving cell lineage using NEBULA^37^ (runNEBULA; core random effect, total-count offset, cpc_thresh = 0). Eligibility required at least five receiver cells per arm and core, five paired cores and 20 cells overall, and models were capped at 20,000 cells by sampling within core and arm (seed 20261025). BH correction covered interactions within each receiving cell lineage, niche coefficient and signal type. Coloured cells require positive logFC, q < 0.05 and a mean LRscore of at least 0.0062 in the higher arm. The panel shows 21 interaction–receiver combinations, and all 189 niche coefficients are in Source Data. Receiver scores do not identify the ligand-producing cell. Receiver-weighted neighbourhood composition multiplied each receiver’s LRscore by the normalized incoming diffusion weights of its neighbours, excluding the receiver itself (200-µm radius; Gaussian threshold 0.001). Contributions were summed by the lineage of the neighbouring cells, averaged within core and then across eligible cores, which required five receiver cells in the niche and in the reference region.

### Metaprogrammes and niche correspondence

Metaprogrammes (MPs) were derived separately from human and mouse single-cell data by non-negative matrix factorization (NMF) of each sample within each cell scope, either all cells or one lineage. Expression was transformed as log2(1 + CPM/10), centred per gene and set to zero where negative, and factorized with NNDSVD initialization over a range of ranks. Robust factors were represented by their 50 top-weighted genes and clustered into MPs by gene overlap following Gavish et al.^121^. A scope required at least 20 cells, and each MP required support from at least two decompositions and two transplant or diagnosis groups. The human catalogue contains 175 MPs, including 16 all-cell MPs, and the mouse catalogue 124 MPs, including four all-cell MPs. MP names summarize member genes and enrichment annotations. Each MP was linked to the cell state with the highest mean programme score. Scores were mean expression of measured MP genes after normalization to 10,000 counts per cell and log1p transformation, averaged within library and cell state (at least ten cells) and then with equal library weight.

Extended Data Fig. 2d shows the Jaccard index between the top-50 gene sets of sample-derived factors, arranged by MP: 366 human factors from 118 libraries and 120 mouse factors. Whole-sample similarity was the Spearman correlation between complete MP-score vectors, where a cell’s score was its mean ln(1 + CP10k) across measured MP genes. Human vectors were library means (120 libraries), and mouse vectors were cell-count-weighted means of cell-state scores within each animal. Supplementary Figs. 37–40 apply the same calculation within 25 human and 12 mouse cell scopes, and scopes with a single MP have undefined correlations. These similarities are descriptive.

Programme–niche correspondence used rank AUC, the Mann–Whitney U statistic divided by the product of member and background gene counts, with ties counted as one half. For all-cell MPs (Fig. 2j and Extended Data Fig. 2e), each of the nine niche-versus-rest rankings contained 18,918 genes from depth-adjusted CosMx niche pseudobulk edgeR specificity models (no intercept; design including niche, diagnosis and standardized RNA depth). Mouse MPs were mapped through one-to-one human orthologues. Two-sided asymptotic Mann–Whitney tests were BH-adjusted separately across 144 human and 36 mouse comparisons. Within-class correspondence (Fig. 2k and Extended Data Fig. 2f,g) used smiDE/metafor gene effects within each lineage. Eligible genes had finite effects, positive standard errors and |log2FC| < 20, with at least 40 eligible genes and five MP members per test. Genes were stratified into standard-error quintiles for van Elteren stratified AUC, and BH correction covered all evaluable MP–niche pairs within each lineage. For each class, Fig. 2k follows the MP with the largest AUC in any niche across all nine niches. Extended Data Fig. 2g links each within-lineage MP to its highest-AUC niche, splitting tied maxima equally, for 79 human and 64 mouse lineage–MP identities.

Supporting analyses in the Extended Data Fig. 2d–g Source Data used three further readouts. GSEA used the same edgeR ranks with 1,000 permutations (seed 42), gene sets of 15–500 genes and BH correction across eight human or four mouse MPs per niche. pyUCell^122^ mean differences were computed in 686 niche-by-biopsy pseudobulks from 83 biopsies, using greater-tail Mann–Whitney tests with BH correction across 72 human and 36 mouse programme–niche comparisons. Competitive enrichment compared mean member log2FC, truncated to [−2, 2], with 10,000 member-sized draws from nonmembers (seed 42). N3 proximal-tubule comparisons tested MIOX, GPX3, ALDOB, FBP1, TINAG, SLC4A4, FGGY and SLC13A1 in 81 sampling units within a 655-gene family.

### TLS detection and staging

TLS were detected by two-step DBSCAN spatial clustering. First, DBSCAN was run on the coordinates of B, T, NK and plasma cells (eps = 15 µm; min_samples = 12). Aggregates in which lymphocytes made up at least 50% of cells and numbered at least 60 were retained. Second, follicles were identified within aggregates by DBSCAN on B cells (eps = 20 µm; min_samples = 12; at least 15 B cells), and follicle-containing aggregates surrounded by at least 20 T cells were counted as TLS. Clustering parameters were calibrated against a null model that permuted cell labels within cores (50 permutations; target FDR 0.05). Two collections were used: 49 TLS in 27 cores (30 Stage I and 19 Stage II) and 167 lymphoid aggregates in 40 cores (102 Stage I and 65 Stage II). Stage was assigned from depth-corrected CR2 and FCER2 signals, with BCL6 and MKI67 used to assess maturation (Fig. 3a). RNA-defined stages describe cross-sectional states; CD21/CD23 morphology is annotated separately. Same-core comparison structures and 27 burden-matched pairs in 19 cores defined the respective comparisons. The protected exterior was same-core tissue within 100 µm of a TLS boundary that was not assigned to another structure.

Programme scores were sparse rank-based scores, a top-rank U statistic with Rmax = 500 in which absent or zero genes received rank 501. The Homing programme comprised CCR7, CXCR5, CXCL12, CXCL13, CCL19, CCL21, SELL, SELP, SELE, ACKR1 and CHST4. Aggregate means were averaged with equal aggregate weight within core and stage. For Fig. 3a, FDC, germinal-centre and high-endothelial-venule scores were compared between stages in the 17 cores containing both, using two-sided paired Wilcoxon tests with BH correction over the three programmes. Interior and protected-exterior core means of 13 programmes were compared by paired Wilcoxon tests separately by stage, with BH correction over 26 tests. Extended Data Fig. 3a shows ten stage-associated metrics in the same 17 cores, with BH correction over ten. Extended Data Fig. 3i shows stage fractions by diagnosis in the 40 TLS-positive cores and by location for the 167 structures. Location was the compartment with the greatest member overlap of at least 10%, or otherwise the nearest compartment within 50 µm by the within-core 10th-percentile member distance. Manual artery, arteriole and vein annotations subdivided vessel-adjacent structures.

### TLS spatial organization

For Fig. 3c, each partner cell’s distance to a hub class was its mean distance to the ten nearest hub cells in the same field and compartment. When fewer than ten were present, all hub cells were used, with at least five required. This distance was divided by λ, the compartment median distance from a cell to its tenth nearest neighbour of any type. The plotted effect is the median paired log2 ratio of inside to outside normalized distance. Paired Wilcoxon tests were BH-corrected within stage, cell level and hub across partners, and only edges with q < 0.05 are drawn.

Radial profiles binned signed distance to the TLS boundary at 10 µm, using mutually exclusive structure masks; negative distances lie inside. Counts summed within structure and bin were normalized by the complete T-cell or whole-cell RNA library of the bin and log1p-transformed. Transcript and pathway gradients (Fig. 3d,e and Extended Data Fig. 3b– d,j) were fitted on aggregate-by-bin observations by weighted least squares with cell-count weights and RNA depth and detection covariates. Standard errors were clustered by aggregate, and BH correction was applied within scope and feature family. Curves were predicted at each stage’s median quality covariates. Pathway scores used the panel genes measurable for each set. The upper block of Fig. 3e contains 12 pathways significant in both stages. The lower block adds pathways significant in at least one stage in ascending order of the Pearson correlation between their stage curves, each with a panel-gene Jaccard similarity of 0.70 or less to terms already selected. Extended Data Fig. 3b,c shows the union of the 20 transcripts and 30 pathways per scope with the largest absolute partial Spearman correlations among those with linear q < 0.05.

T-cell TCF7 (Fig. 3f) and IL7R, CCR7, SELL and MAL profiles (Extended Data Fig. 3e,f) averaged bins with equal TLS weight and were smoothed by LOWESS with span 0.4 and no robustifying iterations. The pooled profile used 49 TLS, 27 cores and 11,998 T cells. Its 16 estimates between −55 and 95 µm were joined by shape-preserving cubic (PCHIP) interpolation at 401 positions. Stage subsets contained 30 Stage I TLS in 18 cores and 19 Stage II TLS in 13 cores. Extended Data Fig. 3g shows subtype fractions (38 subtypes) per aggregate–distance bin, averaged with equal aggregate weight and smoothed with a Gaussian kernel (σ = 1 bin). The transcript-density maps in Fig. 3g were computed on a 5-µm grid with a Gaussian kernel (σ = 35 µm; truncated at 3σ) and are expressed per 10,000 µm². Representative cores were drawn from the 27 cores whose whole-core density maximum for the gene lay within an annotated TLS.

Slingshot^35^ ordering of B and plasma cells and Palantir^36^ ordering of T cells were mapped to 104,856 cells in protected contexts. Radial means used a separate interior category plotted at −5 µm and 10-µm exterior bins to 100 µm. Cells were averaged equally across TLS within core and then across cores within stage and bin, with at least three cores per bin, and cubic smoothing splines were fitted with smoothness selected by generalized cross-validation. The local maps in Fig. 3f each show one Stage I and one Stage II structure with a 100-µm margin. The Slingshot colour limits of 0.366–0.696 are the pooled 1st and 99th percentiles of 1,729 cells in four candidate windows. Extended Data Fig. 3k orders the 167 structures by cell-class composition and shows transcription-factor correlations with this order.

### TLS communication and cellular programmes

Receiver scores were modelled with CytoSignal 0.5.1 runNEBULA within each receiving cell lineage, with a core random effect, total-count offset and cpc_thresh = 0. Diffusion fields extended 200 µm, and contact scores used contact neighbours. Inside-versus-outside models required at least five receiver cells at each level per core. Effects are on the runNEBULA natural-log scale (β), with 95% intervals of β ± 1.96 SE and BH correction within each receiving cell lineage, covariate and signal type. Fig. 3i shows the seven positive pooled effects with q < 0.05, from models containing 24–40 cores. Stage models used Stage I as reference. Diagnosis models used the largest supported diagnosis as reference and at least three cores per level, and compared rejection (TCMR, AMR/MVI and mixed rejection) with BK/CNI, excluding borderline changes.

Extended Data Fig. 3h shows the 13 non-epithelial coefficients with q < 0.05 for pooled or Stage I TLS together with core-paired expression support. Expression was averaged as ln(1 + CP10k) per source and receiver class, using the minimum subunit mean for complexes and requiring three cells per region and six paired cores per route. Core-level log2 ratios of ligand(source) × receptor(receiver) products (ε = 10⁻⁶; clipped at ±10) were tested by two-sided paired t-tests with BH correction across routes within each region. For Extended Data Fig. 3l, the seven pooled effects were paired with eleven non-epithelial source classes. Of the resulting 77 candidate routes, 45 had concordant spatial expression support at both ends. Scores were computed in human single-cell data per sample and in mouse data per animal. A score multiplied the source-cell-weighted mean ligand ln(1 + CP10k) by the receiver receptor mean, taking the minimum across required subunits and requiring ten cells per class. CCL19–CCR7 and CCL19–CXCR3 in T cells, CCL19–ACKR4 and CCL19– CXCR3 in monocytes/macrophages and APP–CD74 in pDC used 11, 10, 10, 11 and 3 spatially supported source classes; IL18 and ENTPD1 used all eleven candidate classes. The IL18 score used IL18, IL18R1 and IL18RAP, and a sensitivity score used IL18R1 and IL18RAP. Group means were scaled to the largest group mean within effect and species and were compared descriptively. Human peritubular and venous endothelium were combined, as were arterial endothelial classes. Supporting CellChat 2.2 summaries in the TLS source atlas pooled human data into eight diagnosis groups and mouse data into model–time groups.

Fig. 3j shows Optimal smiDE/metafor estimates for 69 class–gene pairs in ten classes, comparing TLS interiors with nonmember cells of eligible cores. Each core required at least three cells per class and condition. Candidates met Standard q < 0.05 and |log2FC| ≥ log2(1.5), Optimal models used a Matérn term with knots at 5% of cells (at most 20), and REML synthesis required three sampling units. GO biological-process over-representation used the positive gene lists. For Fig. 3k, MPs were compared with the lineage-resolved TLS effect vectors for pooled TLS, Stage I and Stage II versus exterior. Effects were winsorized to [−2, 2], and competitive AUC ranked MP members against all other tested genes, requiring 40 tested genes and five members. BH correction covered all MP-by-class tests within species and contrast, giving 831 estimable tests across 76 human and 61 mouse MPs. Within each class and catalogue scope, the MP with the largest pooled AUC is shown across contrasts. The leading cell state of all-cell MP16 contained 33 cells from two libraries.

### TLS population dynamics

Pairwise population dynamics (Fig. 3h and Extended Data Fig. 3n) were inferred with OSDR^123^ (tdm v5, commit 7d049700ece90a29bd08c0a354b1d3d599d2784d). Models used 80-µm neighbourhoods with boundary extrapolation, degree-two features (six columns including the intercept), mean death estimation and maximal-density enforcement. Division labels came from protein Ki-67 (noise floor 0.94; threshold 0.5 s.d.) or RNA MKI67 (noise floor 0; same threshold), computed within cell class and compartment. Each class required at least 250 cells in three cores. Each model required at least 1,000 cells, two marker values above the noise floor, two dividing cells, variation in division labels, a full-rank design and finite probabilities and phase fields. Regularization followed α = 0, 0.0001, 0.001 and 0.01, and the first fit that converged for both classes was used. Pooled interior and exterior models used 40 cores. Stage-specific models used TLS interiors and protected exteriors in 33 Stage I and 24 Stage II cores, giving 16 T-cell-containing and six B–plasma-cell combinations. B– plasma models were also fitted in the DBSCAN lymphoid-aggregate compartments and in the union of protected Stage I/II contexts. All 16 T-cell models converged at α = 0, and eight protein and five RNA B–plasma models met the criteria. Three RNA strata could not be estimated: Stage II interior plasma cells had no MKI67-positive cells among 328, and the pooled and Stage I interiors had two among 1,033 and 705. Fields and fixed points were evaluated within the 99.5th-percentile density range. Growth rates are relative model quantities.

### TLS signatures over transplant time

Composite TLS signatures (Fig. 3b) were the unions across lineages of genes with positive Optimal/metafor log2FC and FDR < 0.05 for the pooled, Stage I and Stage II contrasts. They contained 125, 114 and 175 genes, with 119, 110 and 167 mouse one-to-one orthologues. pyUCell 0.5.0 scored integer counts over the full measured gene universe (maximum rank of 1,500; average ties; missing genes skipped; at least five measured members). Scores were averaged per sample or animal across all cells and within eight lineage groups with at least ten cells, using all 352,911 human cells. Time correlations used 59 whole-tissue post-transplant human samples with recorded intervals and mice ordered by POD. The 59 samples came from 26 distinct qualified patients, and the sample was the human unit. Two-sided Spearman tests were corrected across all cells and eight lineages for the three signatures within each species (27 tests). Mouse pDC (1,357 cells) were separated from conventional dendritic cells (11,120 cells). Diagnosis summaries used 106 human samples after excluding uncertain rejection. Control comprised 22 native-kidney and 28 post-transplant reference samples, and the BK group combined resolving BK, viraemia and nephropathy. Extended Data Fig. 3m shows activated B, CD4+ TH17DA, CD8+ TCTLDXt and CD8+ TSMp cells as fractions of their parent lineage in BALB/c grafts at POD7, mild POD28 and POD90 (two animals each), with animals weighted equally.

### Anatomical lesion measurements

Nine immune classes were analysed: T, B, plasma, NK, monocyte/macrophage, dendritic, pDC, neutrophil and mast cells. Mononuclear counts used the seven classes other than neutrophils and mast cells.

Local inflammatory burden (Fig. 4a) was defined by five measures. Interstitial and tubular burden were iST = Nmono/Aresidual (mononuclear cells per mm² of residual interstitium) and tST = 10 × Nmono,wall/Nepi,wall. Vascular burden was vST = 10³ × (Nmono,intima + 0.5)/Aintima (cells per 1,000 µm² of intima). Glomerular burden was gST = Nimmune,glom (immune-cell count), and capillary burden was ptcST = 10³ × Nimmune,ptc/Aptc (immune cells per 1,000 µm² of PTC wall and lumen). N is a cell count and A the corresponding anatomical area.

Glomerular RNA analyses used composite outlines including sclerosis and at least two target cells per structure. Raw immune counts, immune proportions and area density were separate exposures. PTC masks required an area above 100 µm² and anatomical exclusions. TLS-adjacent PTC were included in the primary analysis and removed in a sensitivity analysis. Tubular walls excluded inferred lumen cells. Eligible cortical, non-IFTA, nonatrophic tubules required an inferred lumen and at least five epithelial cells, with ten in a sensitivity analysis. tST tiers 0, (0, 4], (4, 10] and >10 mapped to grades 0–3, and the core grade was the highest grade present in at least two positive tubules. The acute RNA epithelial branch counted nine-class immune cells per ten wall epithelial cells, and joint RNA and receiver analyses counted immune cells per ten non-immune wall cells.

Chronic burden (Fig. 5a) comprised three measures. ciST = 100 × RF/Rall is the scarred fraction of residual interstitial area, and ctST = max(0, 1 − d/dref) is the loss of minimum Feret tubular diameter relative to a reference. cvST = 100 × Ai/(Ai + Al) is the intimal fraction of intimal plus luminal area, with folds removed. Residual interstitium excluded glomeruli, tubules, vessels, PTC and TLS before intersection with scar. The reference diameter was the same-core non-IFTA cortical median when at least ten tubules were available, then the same-slide non-IFTA median and finally the median of all eligible cortical tubules on the slide. Intimal inflammation excluded chronic cv thickening, and its core exposure was the maximum band density. Chronic and active arterial domains were analysed separately. Local exposures used residual tissue within 100-µm tiles. Inflammatory-area measurements used 25-µm pixels with 50-µm neighbourhoods, requiring a mononuclear fraction of at least 0.35 in unscarred cortex, at least 20 mononuclear neighbours in IFTA and at least 80 eligible pixels.

### Lesion-stratified cell composition

Fig. 4b contains 43 composition curves: nine interstitial, ten tubular, eight arterial, eight glomerular and eight PTC series, from 82, 83, 25, 80 and 82 biopsies. Interstitial curves used all-cell fractions within each core and burden quartile. Tubular curves used an epithelial denominator, glomerular curves a non-immune denominator, PTC curves an immune denominator and vascular curves all wall cells. Immune-cell fractions in t used a within-immune denominator, and the tubular immune curve spans tiers 1–3. Ordinal strata were fitted by weighted least squares with cell-count weights, choosing polynomial degree one to three by lack-of-fit and AICc, and continuous vascular burden was smoothed by LOWESS. Fits were unconstrained, and Source Data include fitted values outside the physical fraction range.

Fig. 5c uses all cells of each structural compartment as the denominator, including units with zero counts of a class. The chronic axes were ciST quartiles, ctST tiers and continuous cvST. Each axis shows T, B, NK, dendritic, plasma and monocyte/macrophage cells together with the resident classes of the compartment. Logistic polynomials of degree one to three were selected by leave-one-unit-out fractional-binomial loss, with cell-count weights normalized to one per unit. Pointwise 95% intervals came from 200 bootstrap resamples of units (seed 20261105). The ci and ct branches contained 82 units each, with the two cores of one biopsy as separate units, and the cv branch 53 units with those cores merged. The i-IFTA plasma-cell series used clinical Banff i-IFTA grades 0–3 and all cells of the i-IFTA compartment. Its 56 biopsies each had at least 50 compartment cells, with 11, 24, 16 and 5 biopsies at grades 0–3. The panel shows the first-degree weighted-least-squares fit at 80 points with its 95% confidence interval, and the Spearman correlation in Results is unadjusted.

### Gene–lesion associations and pathway enrichment

Lesion-associated RNA aggregated each target class within a structure, tile or core and normalized it by the complete RNA library. Interstitial associations were within-core tile correlations combined by core-level meta-analysis. Tubular, arterial and glomerular associations used structure-level correlations, with glomerular models requiring at least ten structures and three distinct exposure values. Scar-associated core RNA required at least two immune or three non-immune cells, six informative cores and three distinct exposure values. Spearman, Pearson, quadratic and spline estimates formed separate testing families. Pathway enrichment of the gene rankings used GSEApy^124^ prerank and clusterProfiler^125^, each with its own gene-set libraries and q-value family. Finite-permutation q values of zero are drawn at the display cap.

Fig. 4c shows 50 gene–burden associations and 14 pathway enrichments in 11 lesion and cell-class contexts. Interstitial fibroblast, monocyte/macrophage and T-cell estimates come from core meta-analyses of 46, 65 and 71 cores. Tubular PT, TAL and T-cell series contain 16,480, 5,091 and 1,233 tubules, arterial endothelial and mural series 64 and 104 arteries, and glomerular endothelial and monocyte/macrophage series 332 and 240 glomeruli. The PTC endothelial series uses 82 cores. Gene colours show the coefficient from −0.8 to 0.8, stars mark gene-family q < 0.05, and bubble area shows −log10(q) capped at eight.

Fig. 5d pairs 150 gene tiles with 27 pathway readouts under 15 pathway names. A gene– pathway match required a significant gene, enrichment q < 0.05, leading-edge membership and concordant signs of correlation and NES. Seven readouts, marked by daggers, used whole-core RNA from the joint arterial analysis. They related fibroblasts to the eccentricity-corrected geometry axis and mural or vascular smooth-muscle cells to the inflammation-density axis. Plasma-cell HSPA5, SEC61A1, PDIA4 and XBP1 associations with ciST were within-core tile correlations combined across 32 cores. A supporting dual-axis display selected up to three genes per context by the larger absolute correlation with inflammation or morphology among genes with q < 0.05 on either axis.

### Compartment observed-to-expected ratios

Observed-to-expected ratios (Ro/e)^40^ were computed for all 1,011,984 CosMx cells assigned by centroid to glomerulus, artery, PTC, tubule or interstitium. Within each compartment and parent lineage, expected counts in the cell-state × diagnosis table were row total × column total/grand total. Fig. 4d shows 12 immune cell states across the five compartments on a colour scale from 0.25 to 4. Observed zeros with valid expectations are shown, and absent parents are missing. Ro/e pools counts over biopsies and is descriptive.

### Microvascular and arterial lesion comparisons

Microvascular comparisons (Fig. 4e,g,h) used seven Control, ten DSA-negative/C4d-negative MVI and four active AMR cores, excluding the single probable-AMR core. Arterial comparisons (Fig. 4f,i) used seven Control, three isolated intimal arteritis (isolated v) and eight arteritis-with-tubulointerstitial-inflammation cores.

For pseudobulk expression, raw counts were summed by core and lineage. Each core required at least 20 cells, each arm three qualified cores and each gene a summed count of at least ten, with probe-blacklist genes removed. Each diagnosis was compared with the pooled other two arms in PyDESeq2^126^ with a design containing diagnosis only or additionally standardized sampling year and standardized RNA depth. Expression maps show row z scores of log2(size-factor-normalized counts + 1), capped at ±2, for genes with independent-filtered padj < 0.05 in the adjusted model. The readouts used 7/10/4 cores for PTC endothelium, monocytes/macrophages and NK cells and 5/8/4 for glomerular endothelium (26 entries), and 7/3/8 cores for PT, TAL, monocytes/macrophages and fibroblasts (19 entries). Fig. 4h shows over-representation among genes higher in pooled MVI and active AMR than in Control. One-sided hypergeometric tests used Hallmark, Reactome, KEGG and GO biological-process sets with 10–500 background genes, with BH correction within cell type, contrast and direction.

For spatial organization, Squidpy^127^ 1.7.0 computed within-core neighbourhood enrichment of lineage labels on Delaunay graphs pruned at 30 µm (60 pixels at 0.5 µm per pixel). Immune– endothelial and immune–tubular analyses used 200 and 1,000 label permutations (seed 42). Heatmaps show unweighted means of finite core z scores per diagnosis for PTC/venous and glomerular endothelium and for PT and TAL, with colour limits of ±5 and ±29. The cores contained 42,679 Control, 148,319 MVI, 53,284 active-AMR, 20,100 isolated-v and 118,780 arteritis-with-inflammation cells. Fig. 4e shows the median nearest-neighbour distance from immune cells to PTC endothelium and the directional relative likelihood of immune– glomerular-endothelial contact (7/10/4 and 7/9/4 cores). Fig. 4f shows the immune–tubular enrichment z score and the immune-to-PT relative likelihood. Relative likelihood is normalized for abundance and shown on a log2 scale. Displayed two-sided Mann–Whitney P values have within-metric BH q ≤ 0.05.

Receiver scores for diagnosis comparisons were computed with CytoSignal 0.5.1 from imputed diffusion or contact ligand fields and raw receptor expression (recep.imp = Raw; norm.method = none) and modelled with NEBULA 1.5.6^37^. Models contained analysis-arm fixed effects, a core random effect, the raw total-count offset and cpc_thresh = 0. Receiver strata required at least five cells per core, 20 overall and three cores per diagnosis. PTC and glomerular masks selected receivers, whereas ligand fields used the whole core. BH correction covered all ligand–receptor results within receiving cell type, scope, signal type and diagnosis coefficient. Fig. 4g compares arterial endothelial APP–CD74 receiver scores between active AMR and MVI (2,061 cells; ten MVI and four AMR cores; 2,895 tests in 44 families), with unadjusted 95% Wald intervals. Displayed ligand sources were detected in at least three core aggregates with at least 20 source cells. Source Data contain all 17,628 one-versus-rest and 2,895 pairwise results. An i-IFTA scope (107,914 cells in 81 cores) used borderline changes as the baseline because reference biopsies contain no i-IFTA cells, and its T-cell model contained 54 cores.

Proximal-tubule expression in isolated v (Fig. 4i) was modelled with smiDE Optimal NB2 mixed models. Fixed effects were rank-normalized expression of neighbouring cells of other types and diagnosis, the random effect was a biopsy intercept, and the logarithm of the cell’s RNA count was the offset. Models used 20 Matérn knots per core and 200 in total (seed 20261018; pilot seed 20261104). The model compared three isolated-v and seven Control cores (3,702 and 13,978 cells) across 17,361 eligible genes, with BH correction over the complete family. Positive log2FC indicates higher expression in isolated v. GSEA ranked genes by sign(log2FC) × Φ⁻¹(1 − P/2), with P bounded below at 10⁻³⁰⁰. It used 1,000 permutations (seed 20261018) and 10–500-gene sets from Hallmark 2026.1, Reactome 2022, KEGG 2026 and GO biological process 2026, and Fig. 4i shows positive-NES terms with FDR < 0.05. Transcriptome-wide pseudobulk statistics for HAVCR1 and DCDC2 came from DESeq2^128^ comparisons of three isolated-v cores with seven Control cores and with the 15 other cores (18,923 genes). Analyses used R 4.3.3, smiDE 0.0.2.5 and spaMM 4.4.0.

### Lesion–receiver-score associations

Fig. 4j shows 18 positive Spearman associations between local burden and receiver scores with interaction-family q < 0.05. Fourteen are tubular routes across 10,854 tubules, with immune cells per ten non-immune wall cells as exposure. Three are glomerular routes with raw immune count as exposure, in 313 structures for RSPO1–LGR5 and 380 for EFNA5– EPHA5 and EFNA2–EPHA5. The last is FAM3C–HLA-C across 2,990 PTC with the area-based score. Contact and diffusion scores formed separate families, with BH correction within domain, hierarchy, receiver population, signal type and depth-adjustment branch. In the complete source families, HLA-F–KIR3DL2 used 21,466 tubules and MIF–CD74 240 glomeruli. Point size shows −log10(q), capped at 40. Candidate ligand sources were annotated from full-panel CP10k expression averaged with equal core weight, requiring at least ten cells and three cores.

Anatomical receiver models compared tubular, vascular and glomerular environments with interstitium in NEBULA, adjusting for clinical group, year and depth with a sample random intercept and total-count offset; provisional PTC coefficients were excluded. These models include the 23 SPP1–CD44 records across 82 samples summarized in Supplementary Data 2, seven of them with q < 0.05. Interstitial tile associations used within-biopsy Spearman correlations synthesized by Fisher-z random-effects meta-analysis (DerSimonian–Laird). Each biopsy required at least eight supported tiles, and each test six informative biopsies and five score-positive tiles.

### Diagnosis-associated RNA and local neighbourhoods in chronic lesions

Diagnosis-associated RNA (Fig. 5e) was modelled with smiDE Optimal NB2 mixed models. Each model included neighbouring other-cell expression, a core random intercept, the cell RNA-library offset and a Matérn spatial effect. The estimated marginal mean of each diagnosis was contrasted with the average of the other eligible diagnoses. BH correction covered the candidate genes within compartment, lineage and diagnosis. Scopes were whole-core immune cells, scarred-interstitial fibroblasts, tubular epithelium, PTC endothelium and glomeruli. The 77 smiDE entries are the union of two selections, with 51 shared. One took up to two positive FDR < 0.05 genes per cell type and diagnosis, ordered by FDR, P and gene name (65 entries). The other covered all 48 positive diagnosis–lineage combinations (63 entries).

Local edgeR models used two masks. The i-IFTA leaf mask (102,250 cells) intersected with residual interstitium gave 63,229 cells in 54 cores and biopsies, and local expression and communication used its 39,429 immune cells and fibroblasts. The atrophic-wall mask (moderate or severe diameter loss, lumina excluded) contained 11,971 cells in 79 cores from 78 biopsies, including 7,345 PT cells. Pooled tubular epithelium comprised 10,530 PT, TAL, thin-limb, DCT, CNT, principal and intercalated cells. Raw counts were summed by biopsy and target population, merging cores of one biopsy, with at least 20 target cells per biopsy and three biopsies per arm. PT and pooled-epithelium models included 64 and 67 biopsies.

Each diagnosis was compared with the pooled remainder in edgeR^110^ 4.0.16. The design contained the binary diagnosis, biopsy RNA depth, complexity, negative-probe background, slide and nine depth-driven degradation surrogate variables (dSVs). The dSVs came from composition-standardized profiles of 83 biopsies. The 1,000 genes most associated with depth underwent scaled principal component analysis. Clinical group and Banff t, g, ptc, i, ci, ct, v and cv were protected, and missing scores were median-imputed with indicators. sva::num.sv (BE method, 20 permutations, seed 17) selected nine components. Redundant nuisance columns were removed by pivoted QR decomposition, and designs required full rank and three residual degrees of freedom. Nineteen probe-family genes were excluded. filterByExpr used the full design, and TMM-normalized filtered library sizes formed the offset. Models used estimateDisp(robust = TRUE), glmQLFit(robust = TRUE) and glmQLFTest. BH correction covered tested genes within population, diagnosis, adjustment, count variant and year stratum. Fig. 5e shows six i-IFTA entries, IGHM in five cell classes and T-cell JCHAIN, all from BK contrasts, together with 14 atrophic-PT entries. No PT or pooled-epithelium contrast had a positive FDR < 0.05 result. Unscarred Control interstitium was not part of the i-IFTA comparisons.

Sensitivity analyses repeated the models without adjustment, with quality covariates only and within sampling year. A further analysis thinned each cell with at least 200 transcripts to 200 by multivariate hypergeometric sampling (seeds 0, 1 and 2). In pure interstitium, 26 of 80 primary comparisons were estimable. Five of the six positive entries kept their direction and at least half their magnitude under all three thinning seeds, and three kept FDR < 0.05 under all three. T-cell JCHAIN did not pass filtering with seed 0. All 584 single-year comparisons were non-estimable. DEqual diagnostics correlated sign(log2FC) × √F with degradation-driver t statistics; 20 of 26 contrasts had |r| < 0.1.

Neighbourhood enrichment (Fig. 5f) used Squidpy 1.8.1 on full-core Delaunay graphs clipped at 30 µm and restricted to each mask, with 1,000 within-core label permutations (seed 42) and no edges between cores. Epithelial labels were pooled before permutation for the pooled-epithelium view. The Control fibroblast reference used unscarred residual interstitium from seven reference biopsies (9,888 cells, including 923 fibroblasts). Z scores describe organization within each mask, and diagnoses were not compared statistically.

Communication in chronic lesions was modelled with CytoSignal 0.5.1 and NEBULA 1.5.6 as diagnosis-versus-rest contrasts for 99 i-IFTA and 207 atrophic-wall sender–receiver lineage routes, each with an immune endpoint. Raw ligand counts of the local sender class were combined with raw receptor components of local receivers on full-core diffusion graphs (findNNGauEB; self.weight = 1) or Delaunay contact graphs without renormalization, and scores below one were removed. Models included the binary diagnosis, standardized year and depth, the raw total-count offset and a biopsy random intercept. Receivers required five cells per biopsy, 20 overall and three biopsies per arm (mincp = 5; cpc = 0), with all 735 diffusion and 108 contact interactions as inputs. Primary BH correction covered eligible hypotheses within region, signal type and diagnosis, and a full-input correction covered every route × input. The i-IFTA and atrophic-wall analyses completed 110 and seven models. Two contact routes reached primary significance in families of 161 and one hypotheses, with full-input q = 1 in families of 10,692 and 22,356. No fibroblast–immune route had a positive effect significant under both corrections, and no PT–immune interaction met eligibility. A pooled-epithelium sender analysis of 18 routes × two signal types × eight diagnoses (288 states) had no eligible interaction in 166 states and insufficient replication in 122.

### Programme–lesion correspondence and lesion geometry

Programme–lesion correspondence used rank AUC within each lesion and cell-class gene ranking. The AUC was the fraction of programme member–non-member pairs in which the member ranked higher. Ties counted as one half, and the background included all non-members with finite ranks. Each test required at least five mapped members and 20 non-members. Two-sided asymptotic Mann–Whitney tests were BH-corrected within species, across 142 human and 122 mouse tests. Mouse MPs were mapped through one-to-one Ensembl orthologues. Fig. 4k shows the 53 positive matches with q < 0.05 in 44 contexts, 22 per species. Fig. 5g shows 45 chronic-lesion estimates. They comprise the highest ci AUC within each measured lineage and species (14), the 15 positive ct results with q < 0.05, all 14 estimable cv results and the highest i-IFTA AUC in each species. Clinical i-IFTA rankings were Spearman correlations of year– and depth-residualized expression with grade in scarred-interstitial fibroblasts, monocytes/macrophages and T cells. They used 2,000 high-variance genes per lineage, with separate BH families of 21 human and 17 mouse tests. Programme names and leading cell states come from the metaprogramme catalogue.

Fig. 5h places matched structures in the inflammation–morphology plane: 99 arteries, 11,120 tubules and 10,000 interstitial observations. Diagnosis contours are descriptive 80% bivariate-normal covariance regions with radius √(−2 ln 0.2).

### Lesion signatures in single-cell and external tissue RNA-seq

Spatial lesion gene sets kept the direction and gene-level q values of their lesion models. Lineage sets required q < 0.05 and at least eight source units per gene and took the 250 top-ranked genes. After removal of ribosomal and negative-probe features, each set needed ten source genes and five measured genes, giving 46 direction sets. All-cell sets were the complete up– and down-regulated unions across lineages without the 250-gene limit; genes with opposite directions in different classes entered both. Sources were core or tile pseudobulk for i and ci, joint tubular t and t-IFTA branches for t and ct, raw-count glomerular sources for g, joint arterial v and cv branches and pooled-core ptc sources. The v source contained only IGFBP7, which was scored as a separate single-gene readout. No cv set or independent i-IFTA source qualified, and the t-IFTA PT, DCT and TAL sets contained two, two and three genes, below the size threshold.

Single-cell scoring used all 310,779 human cells from the 110 whole-tissue units and the mouse cells, ranking all 81,388 human or 57,132 mouse genes. pyUCell 0.5.0 used average ties, skipped missing genes and used a maximum rank of 1,500 for lineage sets and 20,000 for all-cell unions. Human genes were mapped to mouse through one-to-one NCBI orthologues. Lineage readouts were cell-score medians within the target class and all-cell readouts means across all cells, each requiring at least ten cells. Human primary tests used one eligible sample per qualified patient, chosen by unit identifier before grades or scores were inspected; mouse units were animals. Two-sided Spearman tests required eight units and variation in grade and score. All six grade pairs were compared by two-sided Mann– Whitney tests with at least two units per grade. BH families were defined by species, endpoint and unit tier, with trends and pairwise tests corrected separately. Of the planned tests, 160 of 166 trends and 590 of 996 grade pairs were estimable. Mouse lesion-grade panels (Supplementary Source Data S284–S285) show human-derived signatures whose animal-level correlation with mouse Banff grade had q < 0.05; syngeneic grafts have no grade.

External validation used 28 formalin-fixed, paraffin-embedded kidney-allograft biopsies from GSE214703: 13 mixed rejection, nine acute TCMR and six stable references^129^. Counts generated in the source study with HISAT2 2.1.0 and HTSeq 0.9.0 against GRCh38 were matched to gene symbols (60,653 features). Each biopsy was scored with pyUCell 0.5.0 over all features (maximum rank of 1,500 for lineage sets and 20,000 for all-cell sets). Banff grades 0–3 were analysed as integers. Two-sided Spearman tests used 19,999 permutations (seed 20261105 plus the endpoint index), with BH correction within the nine all-cell and 46 lineage signatures. The 201 grade-pair Mann–Whitney comparisons were BH-adjusted within the same two families.

### Mouse cell-state co-occurrence

The co-occurrence network (Extended Data Fig. 5a–c) used the cell-state fractions of each animal, with all 79 states in the denominator. The 69 states with more than 500 cells formed nodes. Two-sided Pearson tests covered all 2,346 unordered pairs, and edges required r > 0.3 and unadjusted P < 0.05; no multiplicity-adjusted significance is assigned to edges. Of 448 qualifying edges, 446 are drawn. Highlighted pairs and three-state combinations required distinct parent lineages, at least one immune state and qualifying edges between all members, with all T-cell states assigned to one parent. Triangles were ranked by their minimum and then mean edge correlation. The eligible ranking contained 143 pairs and 216 triangles with 648 constituent edges. Of the triangles, 159 had one immune and two tissue states, 44 had two immune states and 13 had three. The top 20 pairs and triangles are shown, and triangle strength summarizes pairwise correlations without a separate test. Day-28 abundance overlays used Ro/e across mild and severe BALB/c mixed rejection and DBA/2J tolerance (2, 2 and 4 animals), with expected counts from the cell-state × group table. These groups were too small for outcome-specific correlation networks.

### Graft-level trajectories

PILOT 2.0.15^48^ ordered grafts by their cellular composition and state similarity across 34 animals (442,320 cells). The input was a 46-component PCA representation of log1p(CP10k) expression and highly variable genes. PILOT computed unregularized optimal-transport distances from cosine distances between component-wise cell-state medians, with costs scaled by their maximum and composition prior c = 0.1. Diffusion used two eigenvectors, epsilon = 1, alpha = 0.5 and 33 neighbours. ElPiGraph^130^ 0.3.4 fitted a 20-node elastic principal tree (Lambda = 0.01, Mu = 0.1, alpha = beta = 0; seed 20261107). The root was the node closest to the mean diffusion coordinates of the two POD7 syngeneic grafts, and pseudotime followed root-to-terminal paths. Path names describe the experimental and pathological groups distributed along each path.

Diagnostic and lineage composition were smoothed along each path with equally weighted sample-level Gaussian local means. All 79 cell states were assessed as fractions of all cells and of their parent lineage. A state required at least 20 cells in at least 12 animals overall and in at least eight animals on each path. Focal states were chosen from fitted fractions and differences over the interval supported by both paths, together with marker annotations and functional evidence: classical monocytes, pDC, cDC1, MGL2 macrophages, PT FR and PT S1/2. Whole-sample and within-parent fractions were fitted separately with continuity-corrected logits, three-degree-of-freedom natural splines and equal animal weights, and plots show pointwise 95% intervals over the support of each path. The co-abundance network (Fig. 6b) used animal-level fractions of 70 eligible cell states. Pearson correlations across all 2,415 pairs were BH-corrected, and positive edges required r > 0.3 and q < 0.05. For each focal state, the highlighted partner was its most positively correlated state from another parent lineage, with at least one immune state in the pair.

### Molecular associations along graft trajectories

Molecular changes in the six focal cell states were modelled along the two PILOT paths, with animals as response units. Each state required at least 20 cells per animal. RNA responses were sums of integer counts within animal and state. Genes had to be measured in all 34 animals, reach CPM ≥ 1 in at least max(4, ⌈0.2n⌉) supported animals and have at least 20 total counts in the context. Negative-binomial GAMs in tradeSeq 1.24.0^49^ used TMM effective-library log offsets without further depth covariates. Candidate knot numbers ranged from three to min(5, ⌊n/3⌋). evaluateK used up to 60 eligible genes at each of three seeds (20261110, 20261111 and 20261112) and chose the smallest knot number within two mean AIC units of the minimum.

TF responses were animal-by-state means of AUCell regulon scores, requiring at least 20 scored cells, and were fitted with Gaussian identity-link GAMs with zero offsets and k = max(3, min(4, ⌊n/3⌋)). Ligand–receptor responses were square-root expression compatibilities with multimeric subunits. Sender and receiver each required at least 20 cells per animal, and each route and path eight animals. Interactions had to be measured in all supported animals, positive in at least four and nonconstant, and complex positivity required every subunit mean to exceed 0.01. Interactions active in at least half the supported animals were fitted by Gaussian GAMs of asinh intensity scaled by the median positive response. Sparser interactions were fitted by binomial presence models and, with at least eight positive animals and three distinct times, by a conditional Gaussian model of positive intensity. When both components were testable, we doubled the smaller of their two P values and capped the result at 1. The ligand–receptor universe comprised 760 directed routes from focal to partner states on both paths, with 2,377 molecular interactions and at least one immune endpoint among the 79 partner states.

Association and start-versus-end tests were BH-corrected in separate complete families, and Wald P values used the upper tail of the chi-square distribution. Unsupported, ineligible and nonconverged hypotheses entered as P = 1. The association families contained 686,304 RNA, 4,844 TF and 3,613,040 ligand–receptor hypotheses, of which 15,668, 1,168 and 16,571 profiles had q < 0.05. Of 229,825 eligible ligand–receptor hypotheses, 535 had unsupported or nonconverged fits. Each significant profile was evaluated at 400 positions within its observed support. Association q values describe variation along a path, and end-minus-start differences carry their own adjusted tests. Direction-specific over-representation analysis split significant RNA profiles by the sign of the Pearson correlation between each fitted curve and its time grid. The background was all 57,132 gene IDs measured in every animal, tested against MSigDB 2026.1 mouse Hallmark, Reactome, KEGG and GO sets with 15–500 measured members. BH correction was joint across six databases and both directions within each state and path (167,448 tests; 35,961 significant records). ClusterGVis 0.99.12^131^ displayed every significant RNA and TF profile as row z scores ordered by fitted peak.

Mouse regulons were inferred with the SCENIC framework^50,51^ implemented in OmicVerse^89^, separately for each lineage with at least 50 cells. Inputs were the 3,000 highly variable genes and all mm10 motif-annotated TFs, keeping genes detected in at least ten cells. RegDiffusion 0.2.3^132^ inferred TF–target networks from raw counts (1,000 training steps; batch size 128; GPU). cisTarget pruning used the mm10 motif table motifs-v10nr_clust-nr.mgi-m0.001-o0.0.tbl with dropout masking for correlations, thresholds of 0.75 and 0.9, the top 50 targets and the top 5, 10 or 50 regulators. Regulon activity was scored by AUCell on library-normalized (10,000 counts) log1p expression, and regulon specificity scores were computed by cell state.

Tolerance-path screens in Supplementary Data 2 used the same PILOT analysis. Five cell states rose on the tolerance path and fell on the rejection path within their parent lineage. The three with the largest tolerance-path increases, TAL, PT-S3 and PT-S1/2, were selected before molecular or drug results were examined. Molecular responses were fitted with the coefficient-gated Huber procedure described for human trajectories below. PT-S3 had separate families of 114,384 RNA, 924 TF and 380,320 ligand–receptor hypotheses over both paths. An immune–tubular panel of pDC, PT S1/2 and TAL had families of 571,920, 4,230 and 2,567,160 hypotheses. After removal of identities that were also significant on the rejection path, 136 RNA, 11 TF and 18 ligand–receptor profiles remained. RNA and TF profiles numbered 3 and 0 in pDC, 60 and 9 in PT S1/2 and 73 and 2 in TAL. Over-representation analysis of this panel used 6,977 terms per query with 15–500 measured genes, giving 13,954 tests with 246 significant terms for the PT S1/2 positive query and 24 for the TAL negative query.

### Human sample trajectories

Human sample trajectories used 275,474 profiles from 94 tissue collections, with formal molecular fits in 87 representative collections, one per source-project title block. The Control-rooted tree had three paths in PCA coordinates corrected by Harmony for project and technology and by within-state residualization of RNA depth. Path I was TAL– and Control-enriched, Path II PT-enriched and Path III T-cell-enriched; Paths II and III were rejection-enriched. The paths contained 60, 39 and 43 collections and 54, 39 and 42 formal representatives, with shared-trunk collections included in each applicable path. The five focal states were CD16dim_CD56bright NK cells, CD8+ TCTLDM, CCD-IC-A, CNT and aPT. Composition models used continuity-corrected-logit cubic splines with equal weight per representative unit, adjusting for project and median collection log depth. Within-parent and whole-collection fractions were modelled separately, with pointwise 95% intervals.

RNA, TF and ligand–receptor responses were collection-by-state means of log1p(CP10k) expression, AUCell activity and the square root of directed ligand-complex–receptor-complex products. Each state required at least 20 cells or nuclei per collection, and each model eight representative collections and three distinct pseudotimes. RNA features required a zero fraction of at most 95%, and complex support required at least four collections with all subunit means at or above 0.01. PILOT fitted linear, linear–quadratic and quadratic Huber models (epsilon = 1.35). All coefficients, including the intercept, had to reach P ≤ 0.05, and the model with the greatest adjusted R² was selected. Association P values were Pearson correlation P values after model selection. BH correction covered complete families, with unsupported hypotheses and those failing the coefficient criterion set to P = 1. The families contained 1,220,820 RNA, 5,037 TF and 6,515,490 ligand–receptor hypotheses; 52 RNA and 760 TF profiles reached q < 0.05, and no ligand–receptor profile did. Directional over-representation analysis used all 46,709 measured feature IDs as background and 7,161 terms per query from Hallmark, Reactome, KEGG and GO, for 214,830 tests with BH correction across the six libraries within each query.

### Drug prioritization

Trajectory-guided drug prioritization (Fig. 6k and Extended Data Fig. 8a–e) used scDrugPrio^53^ with the six-state mouse RNA, TF and ligand–receptor results. Twelve signatures of all significant RNA profiles and 36 path × cell-state × tier signatures gave 48 queries, transferred to human genes through unique one-to-one orthologues. Tier 1 combined rising RNA, TF genes with rising activity and selected endpoints of rising ligand– receptor profiles. Tier 2 required same-context regulon membership for RNA or ligand– receptor endpoint seeds, and also included significant TFs with significantly rising RNA targets. Tier 3 required rising RNA and rising activity of the same TF together with a regulon-to-ligand/receptor or signed receptor-to-TF link.

Drug targets came from an open ChEMBL adaptation with 366 approved-drug entries and 169 distinct signed target profiles. The human protein-interaction network contained 17,651 nodes and 346,324 edges. Each query used 1,000 degree-matched reference draws (minimum bin size 100; master seed 35; helper seeds 1–1,000). Selection required a closest distance dc < 1 and a one-sided normal-null P = Φ(Z) < 0.05, and within-query BH values across the 366 entries are reported separately. Signals rising on rejection paths were matched to inhibitory or antagonistic target actions, and those rising on the tolerance path to activating actions. Siponimod was excluded from tolerance-activator nominations because sustained S1PR1 modulation causes functional antagonism.

Intercellular weights used the NicheNet prior^52^ with background expression from animal × cell-state means, requiring at least 20 cells per animal, unique orthology and positive mean expression. Foregrounds were the significant RNA profiles in either direction, and 72 ligand-activity evaluations built six-state path-specific graphs from positive Pearson activities. Intercellular eigenvector centrality used the complete path graph. Intracellular centrality used CINNA^133^ eigenvector centrality in the largest connected protein-network component induced by signature and core genes, requiring at least three nodes; 42 components were evaluated. Drug-target centralities were summarized by geometric means. The priority score was the sum of intercellular centralities over qualifying cell contexts plus 0.1 times the mean intracellular drug-target centrality, with average ranks for ties. After review of product identity and functional direction, nominations from all significant RNA profiles comprised 34 rejection inhibitors and 11 tolerance activators, with no ingredient in both lists. Fig. 6k shows the five leading entries plus a Tier 3-supported candidate where available, otherwise the next entry. Guideline roles came from KDIGO transplant-recipient recommendations and the EAU renal-transplantation recommendations^134,135^; other approved ingredients were labelled repurposing candidates.

Species-matched network support used the PathoPlex tiered strategy^72^ on mouse PILOT features, with the same tier definitions and ordered fallback to the first tier yielding approved ingredients. Seeds were expanded through STRING^136^ 12.0 one-hop edges with combined score > 750. Chemical–protein actions came from CTD^137^ records dated 29 October 2026, restricted to 15 protein-process classes with increase or decrease directions in human, mouse or rat and excluding cotreatment and bracketed reactions. CTD reference genes were joined through unique Ensembl 110 orthologues. Ingredients required an FDA approval or EMA human authorization record for the same active ingredient in the snapshots of 6 and 7 November 2026. Activity nominations from the tolerance-path screen required an explicit activity-increase record for the named chemical and protein. Of nine automatically nominated ingredients, four were selected after review of the primary literature of chemical and protein identity, functional response and treatment dependence, and their mechanisms were annotated from those sources^138–141^. A TNFR1 extension followed the PT S1/2 TNF– TNFRSF1A compatibility profile (q = 0.0365192). Tasonermin is the EMA-authorized ingredient of Beromun (recombinant human TNF alfa-1a), and binding and functional experiments show that human TNF signals in mouse cells through TNFR1 and not TNFR2^142,143^. drug2cell^144^ target-expression scores are provided as a separate supporting analysis.

### Cross-species pathway correspondence

Pathway-level correspondence between mouse states and human rejection references was measured with CellSpectra^55^ 0.1.1 using 73 Hallmark or selected GO sets. Human references were built separately for each mouse query, with three pseudoreplicates per reference collection (seed 123; at least ten cells). Comparisons used a shared gene axis with an expression threshold of −1, a gene-count threshold of 10 and a maximum chi-squared value of 4. Query R² measured correspondence, and reference leave-one-out R² summarized reference fit. References represented human AMR, TCMR and mixed rejection. Lineage markers came from a matched-gene leave-one-out analysis: of 379 significant positive markers, 347 were kept by descending mean leave-one-out R², with at most 50 per lineage. Markers were averaged within animal and lineage, then with equal weight across supported lineages and animals. All 34 animals supported endothelial, monocyte/macrophage, PT and TAL comparisons, with 25, 23, eight and six common markers.

Reference-preferential cell-state classification compared target-reference R² values with pooled values from other human cell-state references in the same parent class by one-sided rank-sum tests (scipy.stats.ranksums, alternative = “greater”). BH correction covered finite pathways within each reference × condition × mouse cell state × human cell state family, and classification required q < 0.05 and a positive mean R² difference. The comparison of classical monocytes and PT FR, the two rejection-rising states, covered 15 myeloid and five PT human counterparts across 13 mouse conditions: 260 count-table entries, 43 positive and 217 zero. The leading pair was POD28 DBA/2N PT FR versus human PT FR, with 44 of 69 evaluable preferential pathways and mean R² = 0.4681461114. Gene reports used an expression threshold of zero and kept genes with positive mean expression in both species, giving 438 query–pathway models. The 60 reports shown cover six POD28 DBA/2N animals and 4,039 gene positions. A gene residual was observed mouse expression minus its fitted value from least-squares regression on the first right singular vector of the human reference, divided by the s.d. of reference leave-one-out residuals. Gene inclusion effects were ΔR² = R²(all genes) − R²(without the gene), with the reference axis held fixed.

### Cell-level lineage trajectories and communication in mouse grafts

Cell-level trajectories in Supplementary Data 2 used 13 animals: BALB/c POD7 and mild and severe POD28 grafts and DBA/2J POD7 and POD28 grafts. Up to 3,000 cells per animal were sampled (seed 0). A 15-neighbour graph on 30 Harmony components was clustered by Leiden (resolution 0.6), and Slingshot^35^ 2.18.0 was run on the first 20 components with the root at the cluster with the highest POD7 fraction. Inputs contained 14,332 PT cells, 1,412 fibroblasts, 3,645 neutrophils, 39,000 T cells and 28,944 monocytes/macrophages. Separate CD4 (9,623 cells, including regulatory T cells) and CD8 (27,519 cells) graphs each gave four paths, and the PT, fibroblast, neutrophil and monocyte/macrophage models gave six, three, three and eight. Terminal cells were path members (curve weight > 0.5) with pseudotime at or above the 80th percentile of the path. Each animal’s terminal fraction used its sampled cells of the parent lineage as the denominator, and group means weighted animals equally. A separate outcome-composition display used 17 animals from BALB/c POD7, mild and severe POD28 and POD90 and DBA/2J POD7, POD28 and POD90 (2, 2, 2, 2, 3, 4 and 2 animals).

Raw counts were fitted along pseudotime with tradeSeq 1.24.0 negative-binomial GAMs (seed 61106), with cells as observations and animal as a fixed effect. TMM effective library sizes formed the offset, and genes were tested when detected in at least 5% of path members. Knots were chosen by AIC over 100 genes per model (seed 61107; four to eight knots): seven for PT, six for fibroblasts, neutrophils and CD4 T cells, and eight for CD8 T cells and monocytes/macrophages. Association, pattern, start-versus-end and differential-end tests were BH-corrected separately within each model and test type. Terminal-up genes had within-path association and start-versus-end q < 0.05 with a positive end-minus-start effect. Gene-module heatmaps grouped 69,138 significant gene–path curves into four KMeans modules per path (scikit-learn 1.8.0; random seed 176201). Terminal-up genes were tested for over-representation against the genes tested in each path, using Hallmark 2020, MSigDB Mouse 2026.1 Reactome and GO biological process and mouse KEGG 2026 with 10–500 background genes. Regulon AUCell scores were fitted with Gaussian tradeSeq GAMs using the same animal design and knots, across 462, 426, 241, 411, 411 and 383 positive regulons in the six models. The complete families comprised 2,334 global, 11,125 lineage and 24,587 lineage-pair hypotheses.

Day-28 communication compared cell-state-resolved CellChat^34^ communication probabilities between severe and mild mixed rejection. Routes required estimates in both groups, an immune endpoint and within-condition P < 0.05 in at least one group, giving 16,567 routes, of which 150 records around failed-repair and mature tubules, fibroblasts and neutrophils are displayed. Between-group probability differences are descriptive. Ligand–receptor compatibility along trajectories multiplied each cell’s ligand or receptor complex expression by the mean complementary expression of the partner cell state in the same animal. The resulting 252 features were fitted with Gaussian tradeSeq GAMs. The lineage-specific association family comprised 1,075 hypotheses, of which 212 could not be tested, and 441 significant feature–path profiles from 231 features are displayed. Human SPP1 expression in adaptive and failed-repair tubules was summarized across 87 observations from seven diagnoses.

### Functional-response cohort displays

For the eGFR trajectories in Fig. 7a, days were counted from the clinical report date. Values before graft-terminal events were expressed relative to the median on days −30 to 0, or days −90 to −31 when the primary window lacked measurements, so that no post-biopsy value entered the display baseline. Patient means in 30-day windows were averaged with equal patient weight, and bands show the standard error; day 0 has a log2 ratio of zero. Response labels follow the classification windows defined for the cohort. Baseline characteristics (Fig. 7b) formed 26 comparisons, of which 24 were estimable. Continuous and ordinal variables were compared by two-sided Mann–Whitney tests and 2 × 2 tables by two-sided Fisher exact tests, with BH correction over the 24 estimable comparisons.

### Structural trajectories and the multilevel response model

Structural states comprised 25 anatomical scopes and 59 Slingshot lineages^35^ covering glomeruli, arteries, tubules and their lineage subtypes, interstitial scopes and lymphoid aggregates (Extended Data Fig. 9a–f). Trajectories used unit-pseudobulk RNA with a full-library Pearson representation and PCA correction for depth, complexity, negative-probe background, slide and depth-associated degradation variables. Each scope used an elbow-selected number of components and Slingshot 2.18.0. Roots were Control or low-pathology states, and lymphoid aggregates were rooted at the molecular-stage and FDC reference. Response labels were not used for embedding or root selection.

The multilevel model (Fig. 7c,d,f) used 1,292 ecological features, 93,698 LIANA^119^ ligand– receptor communication features, 270,371 cell-state-resolved gene features, 203 expression-programme features and 25 pathology features. An alternative model replaced the communication layer with 17,236 CytoSignal communication features and was otherwise identical (Extended Data Fig. 9g,h). Gene features represented structural interiors or the 50-µm exterior within each cell state, adjusted for technical depth. CauTrigger3L^56^ encoded ecology into ten latent coordinates, two causal and eight spurious. Its decoder cascade reconstructed communication and expression, then pathology and response. The hidden width was 128. Training used the default four-stage CauTrigger objective for 300 epochs, with weight changes at epochs 30, 120 and 210. Reconstruction weights were 2, 1, 0.5 and 0.2 across the stages, causal and fidelity-KL weights rose from zero to 0.01, 0.1 and 2, and the state-prediction (DPD) weight was 2 in the last two stages. Batches contained 16 biopsies, the seed was 42, and Adam used a learning rate of 5 × 10⁻⁴ and weight decay of 10⁻⁶. Monte Carlo sample numbers were Nα = 25, Nβ = 100 and nx = 5. The learning rate of the first molecular-to-pathology layer was scaled by min[1, √(128/p)], where p is the molecular-layer width. Predictions used the causal posterior mean and the training-set mean of the spurious coordinates. Five-fold cross-validation grouped records by patient, with one index biopsy for each of the 20 patients. Preprocessing and feature eligibility were estimated in the 36-biopsy reference feature set. AUROC and Brier scores were computed from one held-out prediction per patient and describe internal performance after exploratory model selection.

Local sensitivity of a feature was its mean absolute derivative of the non-response logit, computed by automatic differentiation in PyTorch, multiplied by the s.d. of its observed encoded values. Gene sensitivities were averaged over interior and exterior regions and then over structures to rank genes within each cell state. Gene rankings of the alternative model (Extended Data Fig. 9h) were analysed with GSEApy 1.1.12 prerank^124^ using weighted scoring, 1,000 permutations (seed 17) and 15–500-gene sets. MSigDB 2026.2.Hs Hallmark, KEGG MEDICUS, KEGG LEGACY and GO collections were analysed together within each cell state. Of 49 encoded cell states, 29 had estimable tests (50,385 tests), and five terms reached FDR < 0.05. Three were at the high-sensitivity end in regulatory T cells and EC-AVR, and two at the low end in intercalated B cells and DCT cells. Network ribbons combined the adjacent-layer Jacobian, the downstream non-response derivative and the s.d. of the source state, averaging positive and negative patient contributions separately. The networks of the main model (Fig. 7f) and the alternative model (Extended Data Fig. 9g) each show 14 ecology, 20 molecular, 16 pathology and two outcome nodes. The corresponding expanded networks contained 86 and 85 molecular nodes with all 25 pathology features. Extended Data Fig. 9i shows 50 immune-involving cell-state ligand–receptor identities of the main model, each positive in 1–3 of the 20 patients; LIANA scores were one minus the normalized magnitude rank.

Structural expression dynamics used scope-specific tradeSeq models with technical and degradation adjustment and biopsy-clustered Wald F tests. Significant gene–lineage curves (78,864 pairs) were grouped by Ward clustering into 217 modules, with 38 numerically constant curves excluded. Module over-representation used hypergeometric tests against the genes tested in the same lineage. Gene sets were MSigDB 2026.2.Hs Hallmark, KEGG MEDICUS, KEGG LEGACY and GO sets with 10–500 background genes, with BH correction across modules and libraries within each lineage (1,462,939 tests; 13,613 significant). Structure-cluster comparisons used biopsy-paired, unit-weighted pseudobulk edgeR 4.8.2 models (design including biopsy and group) with TMM normalization and robust quasi-likelihood. They comprised 41 contrasts and 374,581 gene tests, with 29,917 significant scope–gene pairs. Structural composition along pseudotime was smoothed by unweighted LOWESS (frac = 0.35; it = 3), requiring eight observations and two distinct times; 5,067 of 5,980 requested profiles were estimable.

### Plasma-cell expression and communication in N9

The plasma-cell contrast (Fig. 7e) compared N9 with all other niches pooled in the 63 cores that had at least three plasma cells in each region. Clinical identifiers linked 42 of these cores to 41 patients. Repeated cores from one patient were combined by inverse-variance fixed-effect weighting before meta-analysis. Forty patients contributed at least one usable effect, with 9–40 patients per gene. smiDE Optimal negative-binomial models included a Matérn spatial effect, other-cell expression and a log full-library offset over all 18,934 measured genes. Assay, overlap, blacklist and detection rules defined 1,258 eligible genes. Log-link coefficients and standard errors were divided by ln 2. Patient effects were combined with metafor^118^ by REML when at least three patients contributed and by fixed effects when two did. For each gene, the REML objective was maximized over log-variance grids of 1,025 and 2,051 points, with refinement of local maxima, the zero-variance boundary and a tail-bound criterion. A 4,097-point scan checked numerical resolution. All 1,258 meta-tests were finite, and 83 genes had full-family q < 0.05. Source Data include sampling-variance diagnostics for 34 genes and all 565 plasma-feature contexts.

Cell maps (Fig. 7g) show complete cores with segmentation polygons and lineage annotations. The non-responder core had the highest N9 and plasma-cell fractions among eligible response-cohort cores, and the responder core was the designated reference. For communication (Fig. 7h), each receiver’s CytoSignal score was multiplied by the fraction of its weighted neighbours annotated as plasma cells. This plasma-adjacency score was averaged within patient and then with equal patient weight. Eleven of 25 ligand–receptor– receiver identities had higher non-responder values for both this score and positive-neighbourhood support. The chord diagram used CytoSignal 0.5.1 countEdges support and circlize 0.4.18, with one equal-width ribbon per identity. For MIF–CD74 to cDC2, patient-equal non-responder and responder means were 0.00084033 and 0.00004043 for plasma-adjacency scores and 0.016128 and 0.050006 for unallocated cDC2 scores. Neighbourhood views (Fig. 7i) used CytoSignal plotEdge on all 37,920 cells of the two cores, with a common cap of 1,719 candidate edges. Positive support required a positive graph weight and receiver score, and highlighted edges also required detected sender MIF and receiver CD74. Both views share coordinate limits of 0–4,715.9609 and 0–1,409.5651 µm, azimuth 0°, elevation 25° and score limits of 0–4.

### In silico perturbation

In silico perturbations (Fig. 7j–l) were performed in the fitted model without retraining. Spurious coordinates were set to their training-set mean, and all other features kept their observed values. A perturbed feature was set to the smaller of its current value and its training-set 10th percentile. Targets were whole-biopsy N9 occupancy and one plasma-cell gene or ligand–receptor feature. This feature had the highest local sensitivity among features with a positive derivative of the non-response logit and a higher observed mean in non-responders than in responders. All encoded views of one gene or ligand–receptor identity were capped together. N9 was capped in the ecological input, with a cap of zero. Plasma-cell ACSS2, selected by these criteria, was capped in the reconstructed molecular layer before pathology decoding, with caps of 0.0511 and 0.0724 for its two interstitial views. Pathology features were not perturbation targets. Of 1,426 candidate features in the plasma-cell communication layer, 14 were direction-concordant, but their reconstructed values lay below the training-set 10th percentile in every patient. Capping would not change any input, so this layer was not perturbed.

The 25 pathology features before and after perturbation were used to generate CauTrigger streamlines. It computed transition probabilities and each patient’s embedding displacement in the two-dimensional causal latent space of Fig. 7d (seed 17; sampled fraction 1; four neighbours; 40 × 40 grid; smoothing 0.8; minimum mass 0.008; scale 0.1). The algorithm replaces undefined correlations with one before exp(correlation/0.005) neighbour normalization, so patients whose pathology state did not change have zero displacement. A non-responder was counted as flowing towards a responder-like state when the cosine similarity between its displacement and the vector towards the responder centroid was positive. A responder was counted as retaining a responder-like state when it had no displacement or when the cosine similarity between its displacement and the vector towards the non-responder centroid was not positive. Changes in predicted non-response probability were computed relative to each patient’s fitted baseline (Fig. 7l). Analyses used CauTrigger 0.0.1, PyTorch 2.11.0 and velocyto 0.17.17.

### H&E niche mapping and spatial features

The H&E classifier (Fig. 8a–c) used physical context fields of 56, 112 and 224 µm to predict the nine spatial niches. The development cohort comprised 10,139 tiles from 49 biopsies in 47 patients, evaluated in five patient-grouped folds, with each tile weighted by the inverse of its patient’s tile count. Per-class F1 and macro-F1 came from the weighted confusion matrix, with patient-bootstrap intervals for macro-F1. Model selection used only this cohort. The same-centre H&E validation cohort comprised 326 sections. Of these, 319 were scored by the five-fold ensemble, and the other seven, from development-cohort patients, by the fold model that excluded that patient. Sections were the units for composition and spatial summaries.

Each tile was labelled with the dominant CellCharter niche in its 56-µm grid square, requiring more than 15 annotated cells and a dominant-class purity of at least 0.60. The central 56-µm field was read at 0.5 µm per pixel and bilinearly upsampled to 224 pixels, the 112-µm field was read at 224 pixels, and the 224-µm field was read at 448 pixels and resized. All fields were normalized with ImageNet means and standard deviations. A shared UNI2-h^145^ encoder (Vision Transformer; patch size 14; 24 blocks; embedding dimension 1,536) embedded each field. Field features were scaled with standardizers fitted on training tiles only and concatenated into 4,608 dimensions. A head with hidden layers of 256 and 128 units (ReLU) and dropout of 0.5 predicted the nine niches. Only the head was trained for the first ten epochs, after which the last two transformer blocks and the final normalization of the encoder were unfrozen. Training used Adam with a head learning rate of 10⁻⁴, exponential decay (γ = 0.95), focal loss^146^ (γ = 2) and sample weights balancing patients and classes. The effective batch of 64 was accumulated from microbatches of eight in BF16 mixed precision. The three fields shared one D4 flip and 90° rotation pose and one colour jitter (brightness, contrast and saturation 0.9–1.1; hue ±0.01), without cropping or zero filling. Inner splits trained for up to 60 epochs with early-stopping patience of ten. The encoder tail learning rate (10⁻⁵ or 10⁻⁴) and class-prior adjustment exponent (0, 0.5 or 1) were selected by nested splits of the training patients in each outer fold. Outer models were refitted for the median inner best epoch number, with seed 20261025. Whole-slide inference averaged class scores of the five fold models and took the highest, without stain normalization. Software was PyTorch 2.13.0 with CUDA 13.0 and timm 1.0.28, and training used one RTX 5090 GPU.

Prediction fields had 56-µm sides, and niche occupancy was the fraction of valid fields assigned to each niche. The adjacency graph connected valid fields sharing an edge. Within each section, 1,000 label permutations preserved class counts within each connected tissue component. Adjacency enrichment z was the observed count minus the permutation mean, divided by the permutation s.d. Nearest-niche distances were directed Euclidean distances between field centres within connected components. Conditional transition entropy H(neighbour label | central label) used up to six neighbours reachable in at most two steps. At a 224-µm radius, relative K was the ratio of observed to expected same-label pairs, G the fraction of same-label anchors with a same-label neighbour and F the fraction of eligible sites with the target class. Anchors had to lie at least the radius from the boundary of the valid grid. Kernel density used Gaussian weights (σ = 112 µm; truncated at 448 µm) normalized by local support, and occupancy Shannon entropy was −Σp ln p. Source Data contain all 741 section-level spatial features. The niche-distance network (Fig. 8e) averaged the two directional mean nearest distances within each section and took the median across estimable sections. Edge weight was 1,000 divided by this distance in µm, and node positions used a NetworkX weighted spring layout (seed 20261107; 100 iterations).

### H&E clinical associations and outcome models

Clinical associations (Fig. 8d,i) used up to 131 patients corresponding to the 326 sections of the validation cohort, with one biopsy per patient selected without reference to outcome. Spearman correlations used complete feature–clinical pairs with at least ten observations and no covariate adjustment. BH correction covered all estimable tests among 920 feature vectors and either 18 Banff fields or 23 eGFR endpoint definitions, and displayed subsets carry the q values of these complete families.

Endpoints were timed from the biopsy request date, with baseline creatinine as the median on days −30 to 0, or days −90 to −31 when needed, excluding post-biopsy values. Known dated graft loss or death truncated follow-up. Three-month improvement was a creatinine decrease of at least 10%, using the value nearest day 90 within days 60–134; observed follow-up fell on days 70–118. In the 45-patient cohort, 24 patients improved and 21 did not.

A spatial composite of 13 inputs comprised nine niche fractions, occupancy entropy, N9–N9 and N5–N8 adjacency z scores and N4–N4 mean distance. It was compared across nine binary and 52 continuous endpoints, with endpoints and features informed by earlier exploratory results. Binary outcomes used L2-penalized logistic regression (C = 0.1; liblinear) and continuous outcomes ridge regression (alpha = 100). Five patient-level folds were shuffled with seed 20261107 and stratified by outcome for binary tasks, with median imputation and standardization fitted within training folds. Clinical comparators used baseline creatinine or eGFR plus Banff i, t, g, ci, ct and cv, and combined models added the 13 spatial features. Performance used held-out predictions: pooled AUC, within-fold AUC weighted by positive–negative pairs, Brier score and log loss for binary outcomes, and MAE, RMSE and R² for continuous outcomes. Probabilities were not calibrated, and no clinical threshold was chosen.

The six-month composite of Fig. 8f,j used the nine niche fractions in 23 patients with a six-month outcome, 14 of whom improved. Outer leave-one-patient-out evaluation used training-only median imputation, zero-variance filtering and standardization. Inner stratified cross-validation selected the logistic C from 0.01, 0.1, 1 and 10 by log loss; C = 1 was selected in 21 outer folds and C = 0.1 in two. The patient-bootstrap interval (1,000 replicates) held predictions fixed and does not include training or selection uncertainty.

Single-feature analyses (Fig. 8g,k and Extended Data Fig. 10a–e) used univariable L2-logistic models with training-fold imputation and standardization (C = 1 for Fig. 8k and Extended Data Fig. 10a–e). They were evaluated over 20 repeats of five stratified patient folds (seed 20261107), averaging within-fold AUC weighted by positive–negative pairs. Features required at least 80% observed values and nonzero variation. In Fig. 8g, each of 920 features was ranked against the response label in 21 patients with a response outcome (13 responders, eight non-responders). N4–N4 adjacency z at 56 µm ranked highest, with mean Brier score 0.151 and repeat AUC range 0.800–0.976 (2.5th–97.5th percentiles), which describes split variation. Fig. 8k used N9 tile fraction for three-month improvement in the 45 patients (Brier score 0.224780), and its coefficient was negative in all training folds. Extended Data Fig. 10a–e shows median nearest N9-to-N5 distance against the response label in 38 patients (22 responders, 16 non-responders). It also shows N5–N9 adjacency z at 56 µm against three-month improvement in 25 evaluable patients (15 improved, ten not). Because features, endpoints and cohorts were chosen with these outcomes, the estimates are internal.

Graft survival (Fig. 8h) was modelled in a 42-patient survival development cohort. Training-only correlation reduction and selection combined 100 evaluable LASSO-Cox resamples with 1,000 random-survival-forest bootstraps and selected one feature, the mean N5-to-N4 distance. Its Cox coefficient was 1.249310 per training s.d. (training mean 233.979341 µm; s.d. 158.187319 µm). The training median defined high– and low-score groups of 21 patients, and the C-index is apparent performance in the development cohort.

## Data availability

In-house data generated in this study are available from the corresponding author by email upon reasonable request. The public-data component of SPECTRA can be previewed through the web portal at https://www.transplantation.com.cn/atlas/. Published datasets were obtained from GEO and ArrayExpress. The 12 projects of the human single-cell reference are GSE109564^73^, GSE145927^8^, GSE151671^75^, GSE169285^74^, GSE195718^76^, GSE195719^77^, GSE228300^78^, GSE230606^80^, GSE242909^79^, GSE284742^26^, GSE317012^81^ and E-MTAB-12051^9^. Longitudinal biopsy data were obtained from GSE284742^26^ and GSE275824^21^, and allograft-biopsy bulk RNA-seq from GSE214703^129^.

## Code availability

Analysis code is available from the corresponding author by email upon reasonable request. Software versions and parameters are given in the Methods.

---

**Extended Data Fig. 1.**
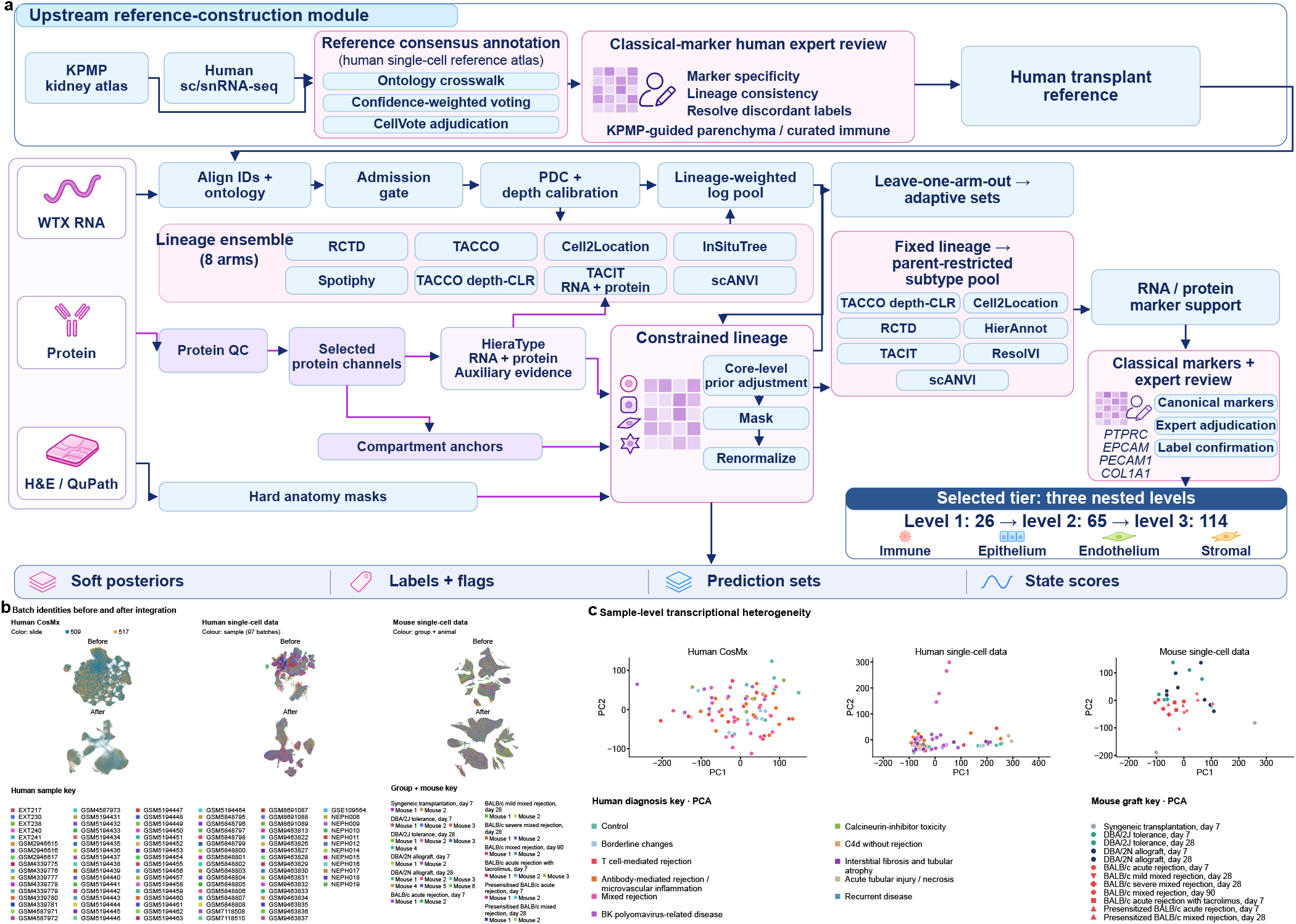
| Spatial annotation and integration of kidney allograft cells. **a**, Unified annotation workflow. Kidney single-cell references are harmonized through ontology mapping, confidence-weighted voting, CellVote and marker review. Eight RNA-based level-1 annotation arms are combined after cell-ID matching and paired-depth calibration. Protein-assisted HieraType, selected protein channels and anatomical masks supply complementary identity and compartment evidence. Core-level priors and compartment constraints guide level-1 calls, followed by parent-restricted subtype assignment and marker review. The selected nested tier contains 26 level-1, 65 level-2 and 114 level-3 labels. b, The same 1,011,984 CosMx cells, 275,474 human single-cell or single-nucleus profiles, and all mouse cells are shown before and after integration; the post-model CosMx view uses the ResolVI latent UMAP. Colours identify the CosMx slide (two slides), the human library (97 libraries) and the mouse animal. The 97 human libraries represent 94 diagnosis-eligible whole-tissue sampling units. The complete keys apply to both integration stages; mice are labelled by experimental group and within-group animal number, with one colour per animal. c, Sample-level RNA-expression PCA, with 83 CosMx biopsies, 97 human libraries and each mouse animal. Diagnostic colours and mouse transplant-group symbols apply to c. Human references combine native-kidney and post-transplant samples; mouse labels distinguish mild and severe mixed rejection. Complete plotting data are provided in Source Data.

**Extended Data Fig. 2.**
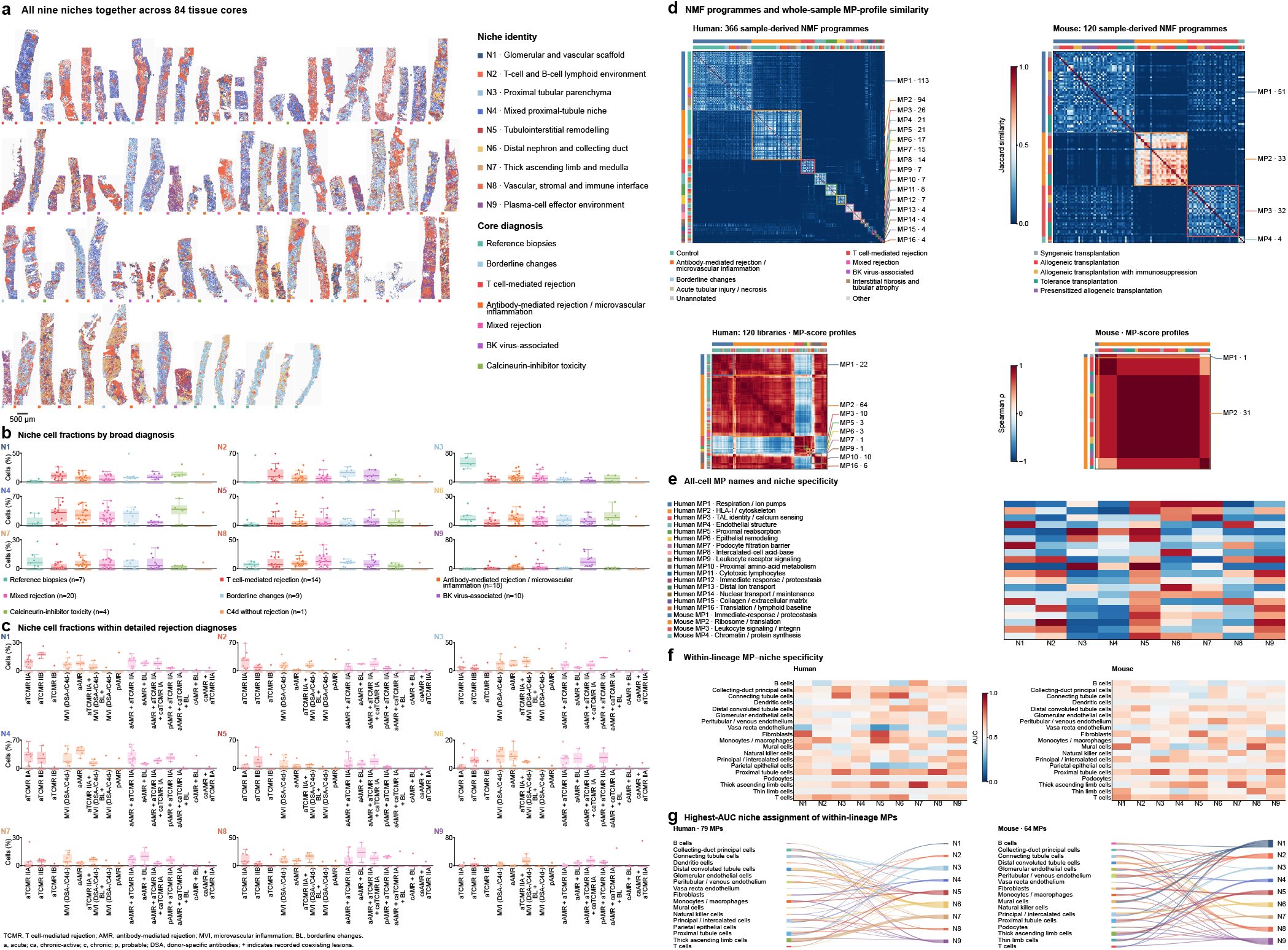
| Tissue niches and their transcriptional programmes. **a**, All nine CellCharter niches shown together in one mosaic of 84 tissue cores from 83 biopsies. Registered cell-segmentation polygons are coloured by their niche identity; small markers denote the seven recorded core diagnostic groups. N4 is the mixed proximal-tubule niche. All cores share the source physical scale; scale bar, 500 µm. **b**, Biopsy-level niche cell fractions across harmonised diagnoses: reference biopsies, n=7; T cell-mediated rejection, n=14; antibody-mediated rejection or microvascular inflammation, n=18; mixed rejection, n=20; borderline changes, n=9; BK virus-associated, n=10; calcineurin-inhibitor toxicity, n=4; and C4d without rejection, n=1. **c**, The nine niche fractions in the 15 recorded diagnostic subtypes within T cell-mediated, antibody-mediated and mixed rejection. Detailed diagnoses are labelled directly below the corresponding boxes, including Banff grades and coexisting lesions; abbreviations are expanded below the panel. Points are biopsies. Boxes show the median and interquartile range; whiskers extend to the last value within 1.5 interquartile ranges. Single-biopsy subtypes are points. Full subtype names, sample counts and all eight broad-group subtype displays are provided in Source Data. **d**, Jaccard similarity of top-50 gene sets among 366 human local NMF programmes from 118 source libraries and 120 mouse programmes from the mouse cohort. Programmes are arranged by MP assignment, giving 16 human and four mouse blocks. Coloured borders and outer strips identify MPs; inner strips identify the source diagnosis or transplant group. Leader labels give the MP number and programme count. Control combines native-kidney and reference libraries. The colour-matched full MP names are listed in b. The lower part of a shows whole-sample Spearman correlations across the complete 16– or four-MP score vectors, comprising 120 human libraries and all mice. Samples are ordered by their highest-scoring MP; rectangles show the nine human and two mouse MPs represented as sample maxima. Both similarities use cold-to-warm colours, with separate 0–1 Jaccard and −1 to 1 Spearman scales. MP and group strips share the upper-layer key. **e**, Niche-specific rank AUC for all 16 human and four mouse all-cell MPs. **f**, For each lineage and niche, the highest stratified AUC among its within-lineage MPs. Panels b and c share the 0–1 AUC colour scale; grey marks unestimated combinations. **g**, Sankey diagrams connect each lineage to the niche with the highest AUC for each within-lineage MP; link width is the MP count, with tied maxima sharing one MP’s total weight. Human and mouse panels contain 79 and 64 lineage–MP identities. Full-size Spearman views and all score components are provided in Source Data. Complete factor-Jaccard views for all available lineage scopes, 48 defined sample-profile scopes, all 299 MP names and leading cell states, and full MP-level niche matrices are provided in Source Data.

**Extended Data Fig. 3.**
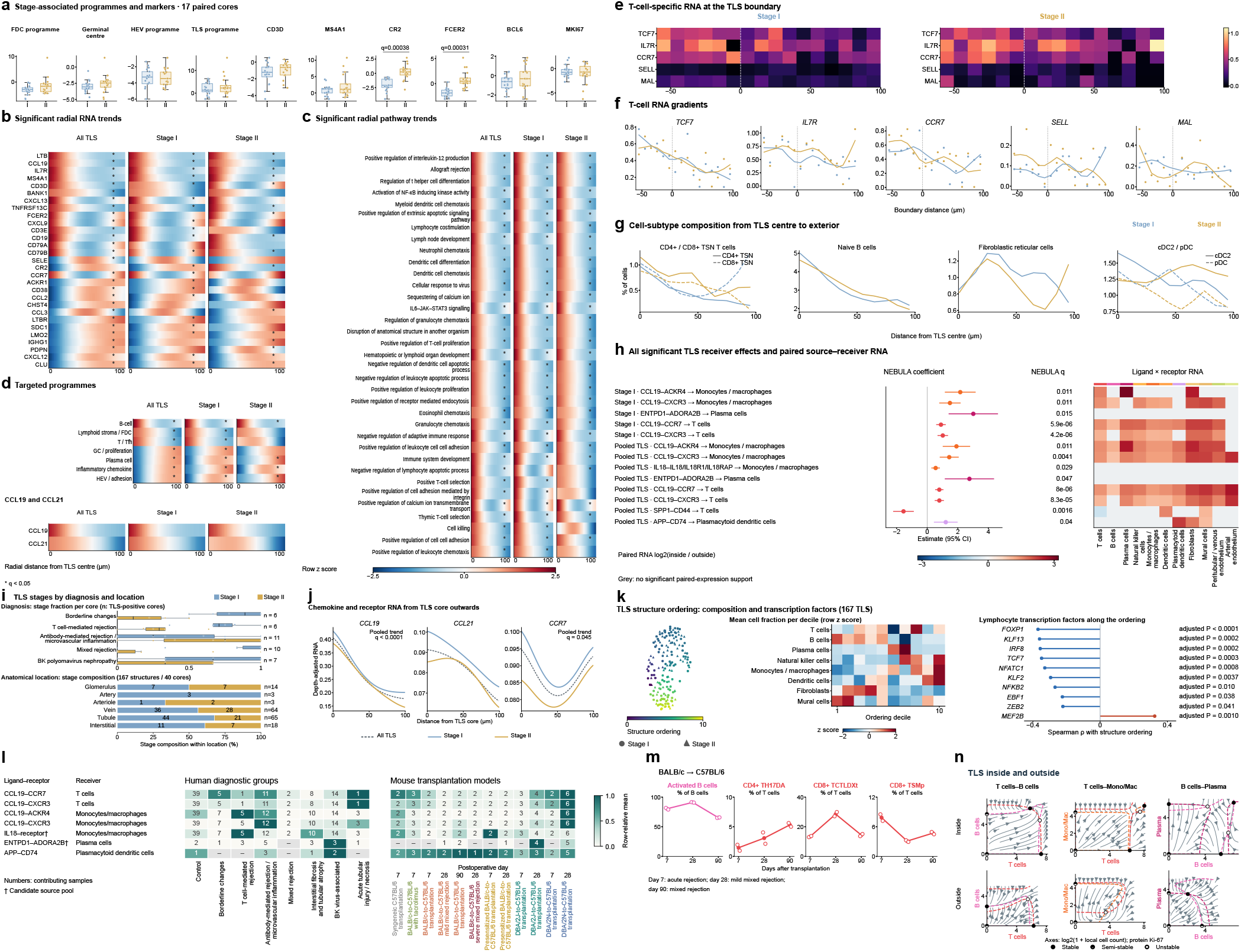
| Cellular organization, signalling and dynamics of tertiary lymphoid structures. **a**, Ten stage-associated measurements in 17 cores containing both stages. Each point is a stage-specific core mean. Boxes show the median and interquartile range; whiskers extend to 1.5 interquartile ranges. Two-sided paired Wilcoxon tests were corrected across ten measurements, with q < 0.05 values labelled. **b,c**, The 20 strongest significant transcript gradients and 30 strongest significant pathway gradients per scope were ranked by absolute partial Spearman correlation. Their unions contain 29 transcripts and 35 pathways. Columns show pooled TLS, Stage I and Stage II. **d**, Seven targeted programmes and CCL19/CCL21 smooth heatmaps. Rows are selected using linear q < 0.05 in at least one scope; each selected row shows every estimated scope fit. Asterisks mark q < 0.05. Both chemokines are shown across all scopes. Colour denotes within-feature z scores of depth– and detection-adjusted fitted curves. Grey denotes an unestimated fit. **e,f**, T-cell-specific TCF7, IL7R, CCR7, SELL and MAL RNA bin means and LOWESS curves. These panels include 30 Stage I TLS from 18 cores and 19 Stage II TLS from 13 cores. Negative boundary distances lie inside TLS. Per-bin contributors, fitted values and feature statistics are provided in Source Data. **g**, Stage-specific fractions of CD4+/CD8+ TSN T cells, naive B cells, fibroblastic reticular cells and cDC2/pDC from the TLS centre to exterior. Fractions use all cells in each aggregate–distance bin and equal aggregate weights, with Gaussian smoothing (σ = 1 bin). Solid and dashed curves distinguish CD4+ and CD8+ TSN cells, and cDC2 and pDC, respectively. The full 38-subtype distribution is provided in Source Data. **h**, Thirteen significant non-epithelial CytoSignal NEBULA receiver coefficients for pooled or Stage I TLS versus exterior, with 95% confidence intervals and adjusted P values. Twelve coefficients are positive and the SPP1–CD44 T-cell coefficient is negative. Stage II has no coefficient at q < 0.05. The matrix shows 77 significant core-paired ligand(source) × receptor(receiver) RNA-product routes covering ten receiver effects. Colour is the mean paired log2 ratio; grey denotes no significant matched expression route. Seventy-five routes agree in direction with the receiver effect. **i**, Stage fractions by diagnosis in 40 TLS-positive cores, preserving each core as a point, and stage composition within six anatomical locations across 167 structures from the same 40 cores. Diagnosis n counts cores; location n counts structures. Vessel-adjacent structures comprise 3 artery-, 3 arteriole– and 64 vein-adjacent structures. **j**, Depth-adjusted CCL19, CCL21 and CCR7 curves from the TLS centre to its exterior, with significant pooled q values. **k**, UMAP and ordering of 167 TLS structures, cell fractions by ordering decile and transcription-factor correlations. Shape denotes stage. **l**, Seven positive pooled spatial receiver effects shown as sample-level ligand–receptor expression products across human diagnoses and mouse model/postoperative-day groups. Scores multiply source-cell-number-weighted mean ligand ln(1 + CP10k) by the minimum receiver-subunit mean, requiring at least ten cells in each population. Five effects use spatially supported source rosters; daggers mark IL18 and ENTPD1 candidate source pools. Colour is the mean eligible sample score divided by the maximum group mean for that effect within species. Numbers count contributing materials or animals; grey denotes unavailable scores. **m**, Activated B, CD4+ TH17DA, CD8+ TCTLDXt and CD8+ TSMp proportions in BALB/c-to-C57BL/6 grafts at days 7, 28 and 90; day 28 uses mild mixed-rejection samples. **n**, Protein-Ki-67 pairwise OSDR phase portraits inside and outside TLS from 40 cores. T–B models contain 38,489 and 14,831 cells, T–monocyte/macrophage models 31,348 and 14,702 cells, and B–plasma models 12,282 and 4,727 cells inside and outside, respectively. Arrows show fitted dynamics and dashed curves show nullclines. Filled, half-filled and open points denote stable, semi-stable and unstable fixed points. Axes show log2(1 + local cell count), with shared limits across compartments within each pair.

**Extended Data Fig. 4.**
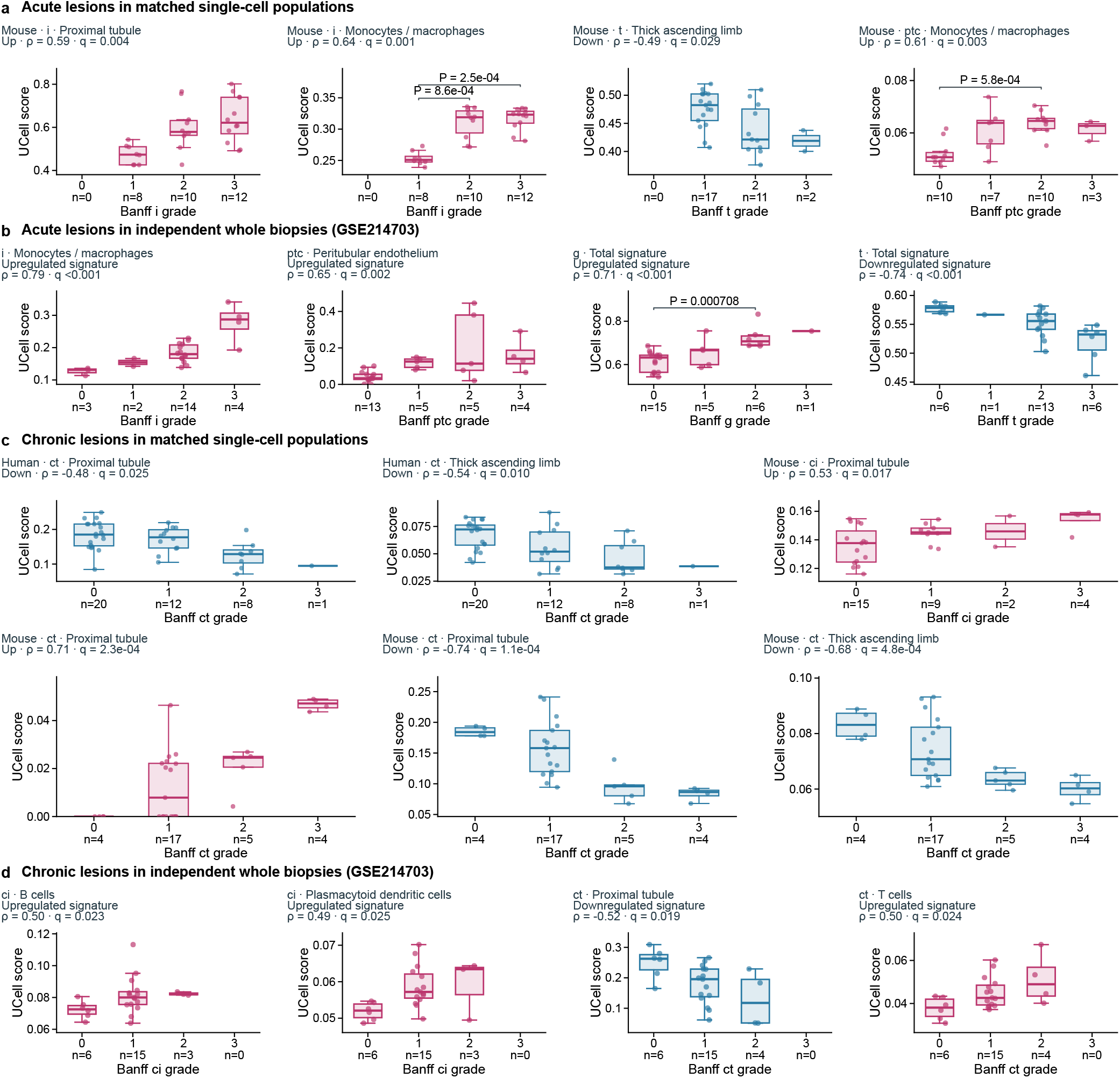
| Acute and chronic lesion programmes in single cells and whole biopsies. **a**, Acute lesion programmes including spatially derived interstitial-inflammation programmes in proximal tubules and monocytes/macrophages, the downregulated tubulitis programme in thick ascending limb cells, and the monocyte/macrophage peritubular-capillaritis programme scored within their corresponding mouse cell classes. Each dot represents an independent animal; values are the median pyUCell score within the indicated cell class. **b**, Lesion programmes in the independent GSE214703 cohort of 28 FFPE kidney-allograft biopsies. The first two gene sets originated from monocytes/macrophages and peritubular endothelium; the glomerulitis-up and tubulitis-down sets are the full direction-specific unions across source cell classes. Bulk scores describe whole biopsies. Up and down denote the direction of the spatial association and use magenta and blue, respectively. Boxes show medians and interquartile ranges; whiskers extend to the most distant observation within 1.5 interquartile ranges. Grade-specific counts appear below all four grade positions. Titles give two-sided Spearman ρ and BH q; single-cell and external bulk tests use separate families. Bars show two-sided Mann–Whitney P values for comparisons meeting complete-family q<0.05. Lineage-derived sets use a maximum UCell rank of 1,500 in both single-cell and bulk data, and full all-cell unions use 20,000. All planned human and mouse lineage and whole-sample all-cell boxplots, gene memberships, eligibility and tests are in Source Data (pp. 2– 24). The acute gene-correlation and pathway page is on Source Data p. 25. **c**, Chronic lesion programmes including the spatially downregulated tubular-atrophy programmes scored within proximal tubule and thick ascending limb cells from human and mouse single-cell data, together with the mouse proximal-tubule fibrosis-up and atrophy-up programmes. Human dots are qualified recorded patients represented by one eligible sampling material selected before inspecting grades or scores (41 patients per displayed readout); mouse dots are independent animals. Values are matched-cell median pyUCell scores. **d**, Spatially derived fibrosis programmes from B cells and plasmacytoid dendritic cells, and atrophy programmes from proximal tubules and T cells, scored across whole biopsies in GSE214703. Lineage labels identify the origin of the gene sets, and all displayed lineage-derived bulk sets use a maximum UCell rank of 1,500. Colours, box and whisker definitions, four grade positions, sample counts and statistical labels follow Extended Data Fig. 4a–b. Up and down sets keep the direction of the spatial association. Single-cell and bulk trend and pairwise tests use their respective complete BH families. All 166 planned lineage and all-cell readouts, including the separate human sampling-unit sensitivity analysis, are shown in Source Data pp. 5–18. Source Data pp. 2–4 describe lesion coverage and all trends, p. 19 shows the separate IGFBP7 gene-specific readout, and pp. 20–24 contain all estimable bulk boxplots. The chronic gene-correlation and pathway page is on Source Data p. 26.

**Extended Data Fig. 5.**
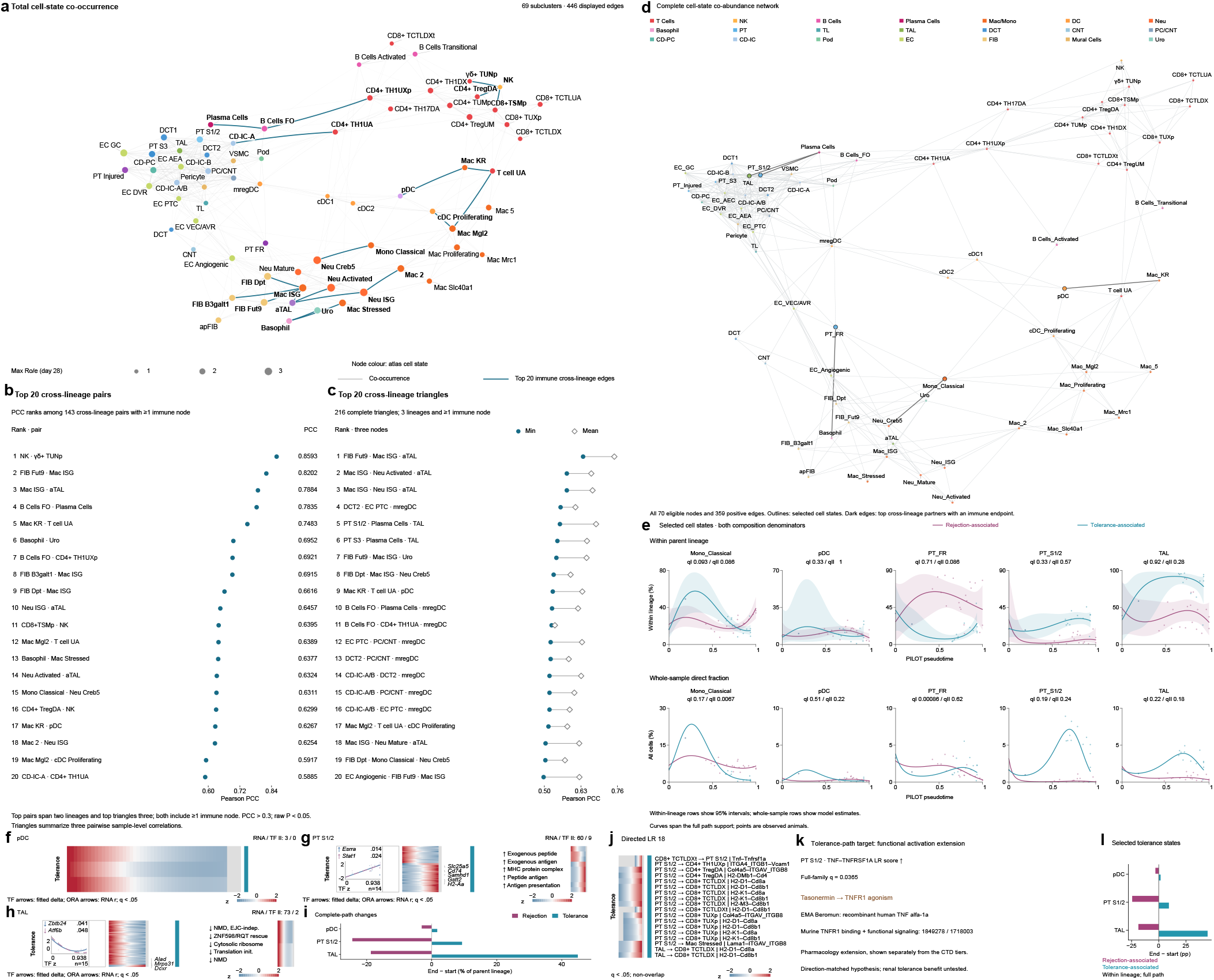
| Mouse cell-state co-occurrence and tolerance-associated trajectories. **a**, The full 69-node co-occurrence network with the 20 highest-PCC qualified cross-lineage pairs with an immune endpoint highlighted. **b**, The top 20 of 143 qualified cross-lineage pairs with at least one immune endpoint, ordered by descending PCC, ascending unadjusted P and state names. **c**, The top 20 of 216 complete three-state combinations involving three different parent lineages and at least one immune lineage. Every triangle contains three edges meeting PCC>0.3 and two-sided unadjusted P<0.05 across all animals. Triangles are ordered by descending minimum PCC, descending mean PCC and state names. Node colours follow the atlas cell-state colours, and node area represents the maximum day-28 Ro/e across the three outcomes. The network shows 446 qualified edges; all 448 qualified statistics are in Source Data. Pairwise fractions use all 79 cell states per animal, and eligible nodes have more than 500 cells in the complete cohort. Triangle strength is summarized by the minimum and mean of its three pairwise correlations. All T-cell states, including CD4 and CD8, share one T-cell parent lineage for these rankings. Complete eligible pair, triangle and 648 constituent-edge records are available in Source Data. **d**, Positive co-abundance network of 70 mouse cell states, with 359 edges at Pearson r > 0.3 and BH q < 0.05 among 2,415 pairs. Outlines mark the five focal states, and highlighted edges their strongest cross-lineage partners involving an immune cell. **e**, Composition curves for classical monocytes, pDC, PT FR, PT S1/2 and TAL with the parent lineage and all cells as denominators. Points are independent animals, lines are spline means and ribbons for within-lineage fractions are pointwise 95% intervals; association q values are given for paths I and II. The cohort comprises 34 animals; the two paths contain 21 and 15 animals, two of which lie on the shared trunk. **f–h**, Tolerance-path PILOT profiles in pDC, PT S1/2 and TAL that do not overlap with rejection-path molecular identities, totalling 136 RNA and 11 TF profiles (RNA/TF, 3/0, 60/9 and 73/2). Rows follow the fitted peak and colour is the row z score; significant regulons are shown as Huber-polynomial curves with animal AUCell means, association q values and the direction between fitted endpoints. Directional ORA used 57,132 measured mouse genes as background, with BH correction across 6,977 predefined terms within each cell-state, path and correlation-sign query. After excluding rejection identities, the PT S1/2 positive query (38 genes) and the TAL negative query (70 genes) yielded 246 and 24 significant terms. **i**, Change in the within-lineage fraction of the three states between the end and start of the path. **j**, All 18 nonoverlapping PILOT ligand–receptor profiles involving an immune cell in pDC, PT S1/2 and TAL; rows give the sender, receiver and molecular subunits, and colour is the row-standardized compatibility within the fitted range. **k**, The rising PT S1/2 TNF–TNFRSF1A compatibility profile (complete-family q = 0.0365) matched to the ingredient tasonermin, with receptor-binding and functional-signalling evidence for human TNF acting through murine TNFR1. **l**, Change in the within-lineage fraction of the three states across the full path, in percentage points.

**Extended Data Fig. 6.**
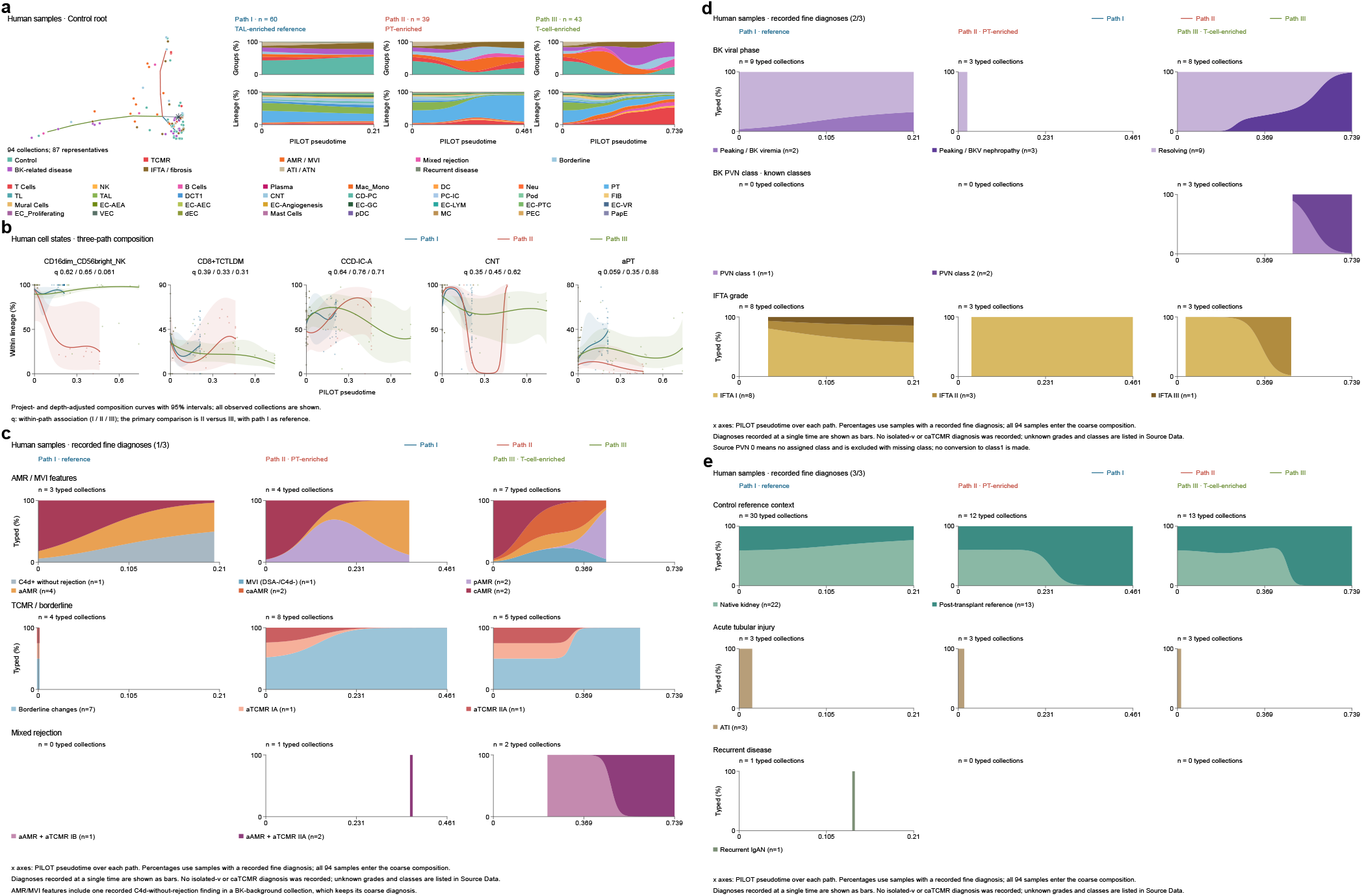
| Human graft trajectories and diagnostic composition. **a**, Control-rooted PILOT tree of 94 tissue samples with three paths enriched in thick ascending limb, proximal tubule and T cells. Including the shared trunk, the paths contain 60, 39 and 43 samples. Coarse diagnosis and lineage composition along each path are shown on the right. **b**, Within-lineage fractions of CD16dim CD56bright NK, CD8+ TCTLDM, CCD-IC-A, CNT and aPT cells. Points are samples; fitted means and pointwise 95% intervals are adjusted for project and sequencing depth, and composition models used 87 representative samples. **c**, Recorded AMR/MVI features, TCMR/borderline change and mixed rejection along each path, with samples of known category as the denominator and actual counts shown for each category. **d–e**, Profiles show BK viral phase, BK polyomavirus nephropathy class, IFTA grade, control reference context, acute tubular injury and recurrent disease. Each panel gives the counts of samples with a known category and the pseudotime range of each of the three paths. Percentages use known categories within each clinical family as the denominator; shared-trunk samples contribute to every path they belong to, and a single known time point is shown as a fraction bar. Unassigned BK classes and missing grades are excluded from the denominator, native-kidney and post-transplant references keep their recorded labels, and the coarse diagnostic profiles include all 94 samples.

**Extended Data Fig. 7.**
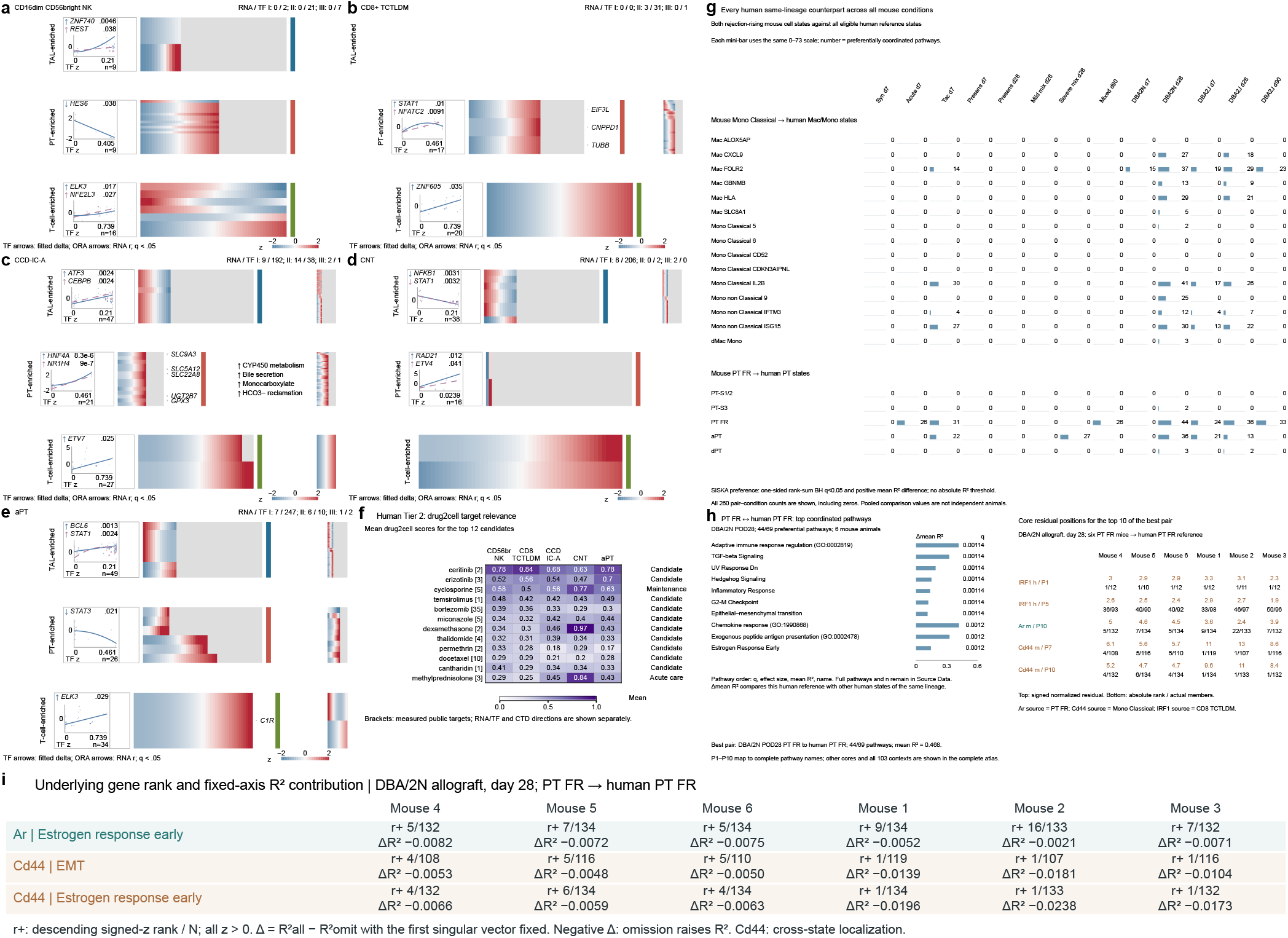
| Human cell-state programmes and cross-species pathway correspondence. **a–e**, Three-path RNA and regulon matrices for NK, CD8+ TCTLDM, CCD-IC-A, CNT and aPT cells, showing all 52 RNA and 760 TF profiles that met the coefficient criteria and complete-family q < 0.05. The RNA, TF and ligand–receptor families comprised 1,220,820, 5,037 and 6,515,490 hypotheses, and no ligand–receptor profile reached this threshold. Columns follow pseudotime, rows follow the fitted peak, and colour is the row z score. TF curves are Huber polynomials evaluated at 400 positions within the fitted range, and points are AUCell means of representative samples; numbers are association q values and arrows show the direction between fitted endpoints. Directional pathways used ORA against a background of measured features. **f**, drug2cell target-expression scores of 12 candidate ingredients across the five states, with the number of measured targets in brackets and clinical-use classes indicated. The replicate unit is the 87 representative samples among 94. **g**, Pathway counts for all 260 pair–condition combinations formed by pairing mouse classical monocytes and PT FR with 15 human monocyte/macrophage and five human proximal-tubule states across 13 mouse conditions. A pathway was counted when the one-sided rank-sum BH q was below 0.05 and the mean R² for the target reference exceeded that of the other references; 43 combinations had positive counts and 217 had none. **h**, The pair with most correspondences, day-28 DBA/2N PT FR and human PT FR, preferentially matched this reference for 44 of 69 evaluable pathways (mean R² 0.468). Normalized residuals of ten selected pathways and their core genes are shown for all six mice. **i**, Descending ranks of signed residuals and omission effects on a fixed reference axis for *Ar* and *Cd44*. ΔR² is the all-gene R² minus the R² without that gene, with coordinates on the first reference singular vector held fixed. *Ar* comes from mouse PT FR, *Cd44* from mouse classical monocytes and human IRF1 from CD8+ TCTLDM cells. Each mouse contributes one value to the target reference, whereas pooled comparator references can include the same mouse more than once.

**Extended Data Fig. 8.**
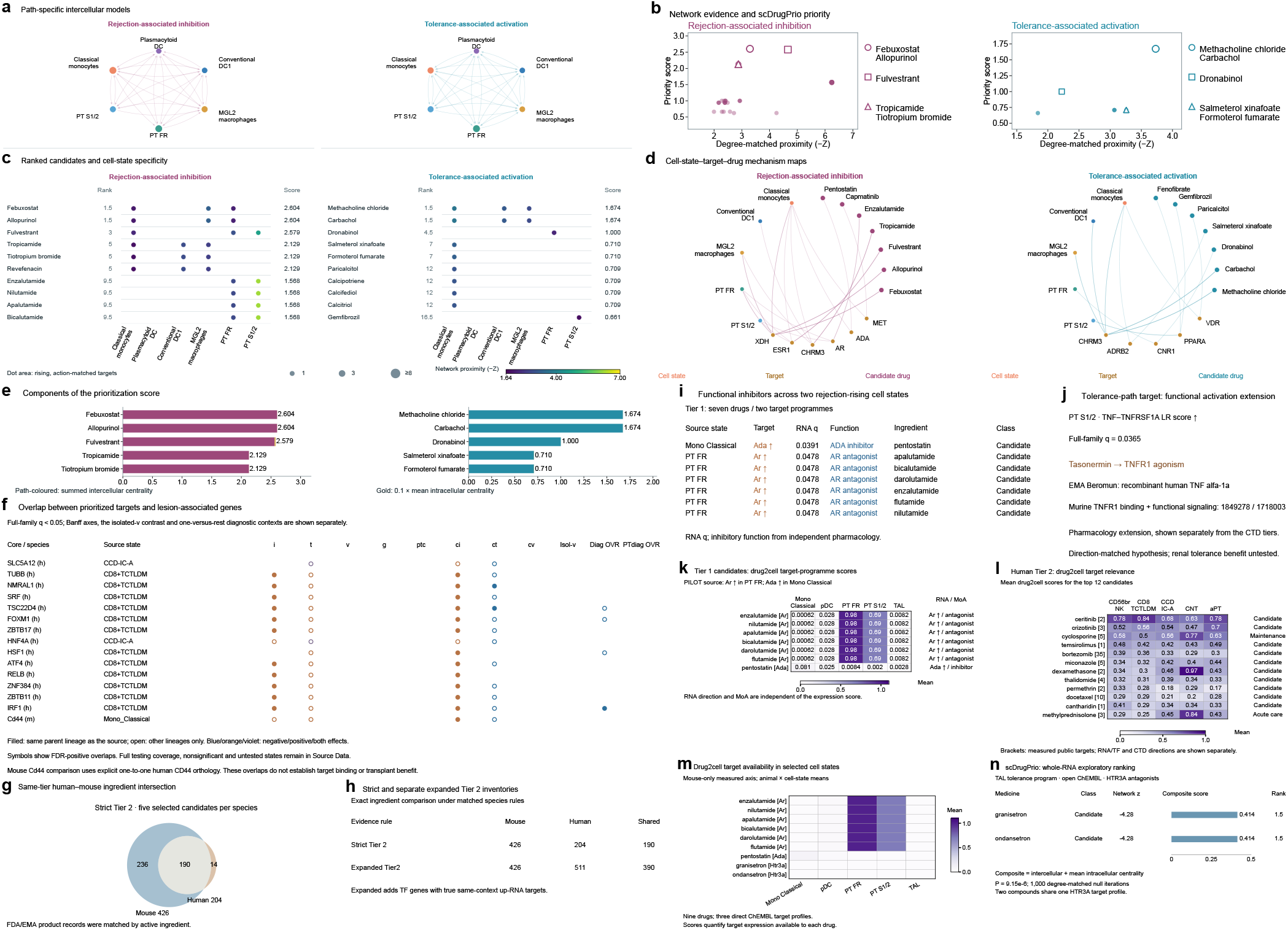
| Trajectory-associated targets and pharmacological candidates. **a**, Path-specific intercellular networks inferred from the RNA programmes of six mouse cell states using the NicheNet prior. Arrows denote positive ligand-activity associations, width denotes mean Pearson activity across supporting ligands, and node area denotes intercellular eigenvector centrality. Illustrated links include an immune endpoint. **b**, Whole-RNA scDrugPrio priority versus degree-matched network proximity, separately by path; dot area denotes the number of qualifying cell states. **c**, The first ten candidate entries per direction in score order. Dots identify qualifying cell contexts; colour is −Z and area the number of matched rising targets, capped at eight. Tied candidates share their average rank. **d**, Representative cell-state–target–drug networks spanning leading candidates, target families and Tier 3 support. Target symbols denote human orthologues of mouse-source genes. **e**, Score components for the first five entries: summed intercellular centrality (path colour) plus 0.1 times mean intracellular target centrality (gold). The whole-RNA nominations comprise 34 rejection-associated inhibitors and 11 tolerance-associated activators. Molecular inputs require complete-family FDR<0.05. Network selection uses dc<1 and one-sided normal-null P<0.05 after 1,000 degree-matched reference iterations; network P values are separate from molecular FDR. The ChEMBL adaptation contains 366 approved-drug entries and 169 distinct signed target profiles. Daggers mark ingredients that match guideline therapies. Priority scores rank computational candidates within each path and scope. Complete tier-specific results, pharmacological directions, source-cell identities and all candidates are documented in the accompanying study records (Supplementary Data 2 and Source Data). **f**, Overlap of PILOT core genes with FDR-positive lesion genes from Figs. 4 and 5, labelled by source cell state and species. Filled symbols denote the same lineage and open symbols a different lineage; colours show the direction of effect. **g,h**, Ingredient-level overlap between human and mouse under the same species-matched PathoPlex rules: strict Tier 2 contained 426 mouse, 204 human and 190 shared ingredients, and TF-expanded Tier 2 contained 426, 511 and 390. **i**, Seven direct-function candidates for two RNA programmes: the ADA inhibitor pentostatin in classical monocytes and six AR antagonists in PT FR. **j**, A separate tolerance-path activation extension links the rising PT S1/2 TNF–TNFRSF1A compatibility profile (q = 0.0365) with tasonermin and published evidence of murine TNFR1 activity. RNA direction, target pharmacology and clinical use are defined separately. **k,l**, drug2cell target-programme scores of mouse Tier 1 and human Tier 2 candidates in their respective five states, with measured targets in brackets; each row is the mean of one ingredient in its source cells. **m**, Target-expression availability in mouse for nine ingredients representing three direct ChEMBL target profiles. **n**, In a separate whole-RNA scDrugPrio query of the TAL tolerance programme, granisetron and ondansetron share one HTR3A antagonist target profile and rank jointly at 1.5. Their network Z is −4.28 (P = 9.15 × 10⁻⁶ from 1,000 degree-matched reference draws) and their score is 0.414. The direction-matched screen of the six main cell states is shown in Fig. 6k and Extended Data Fig. 8a–e.

**Extended Data Fig. 9.**
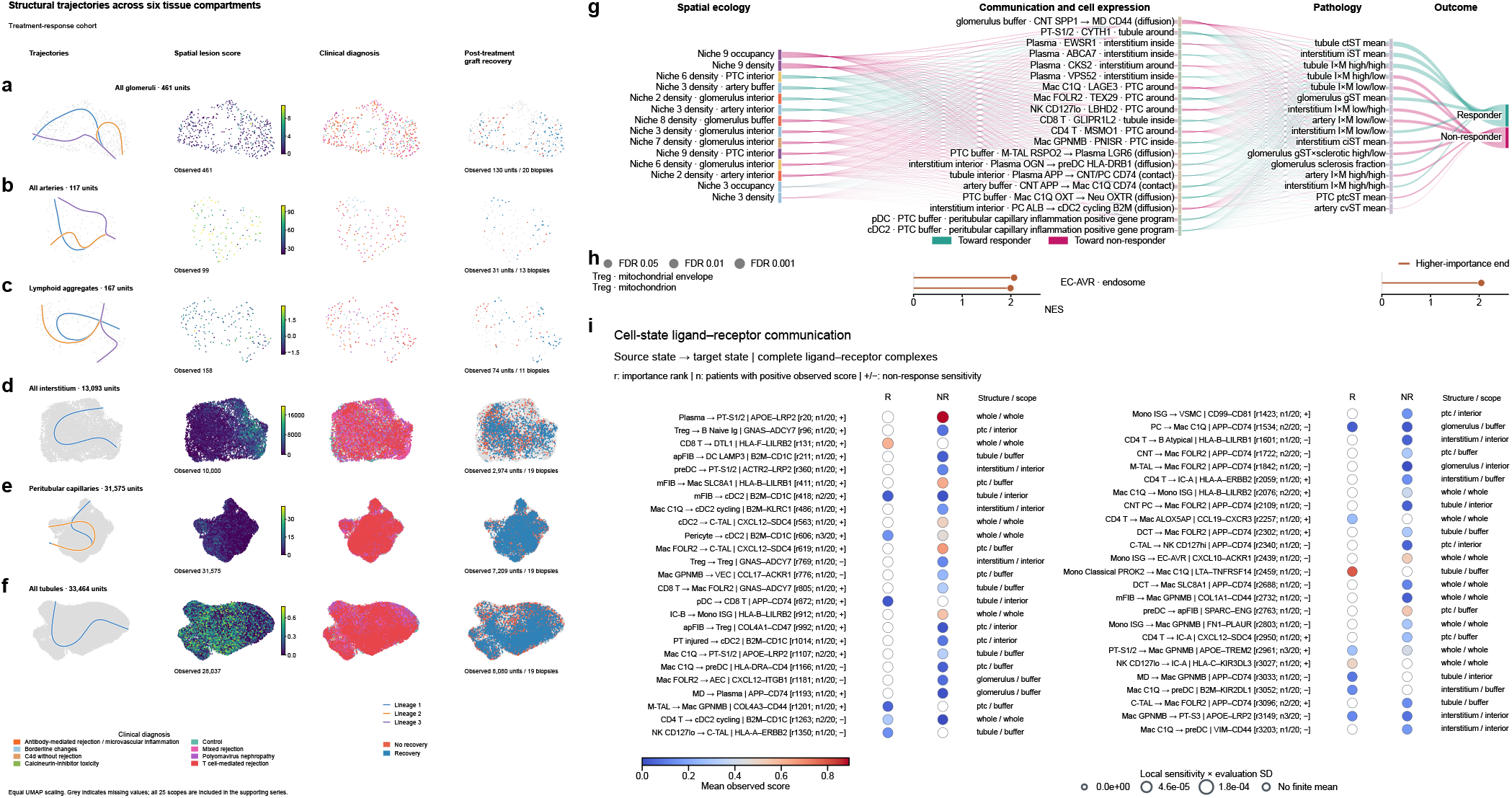
| Structural trajectories and molecular networks of graft response. Rows show all glomeruli (a, 461 units), all arteries (b, 117), lymphoid aggregates (c, 167), all interstitium (d, 13,093), peritubular capillaries (e, 31,575) and all tubules (f, 33,464). Columns show Slingshot curves, the primary observed spatial lesion score, clinical diagnosis and functional response. Each scope uses its expression-derived UMAP coordinates with equal x/y unit scaling. Curve colours identify lineage numbers separately within each compartment. Grey indicates an unobserved value. Response overlays contain 130 units from 20 biopsies (a), 31 from 13 (b), 74 from 11 (c), 2,974 from 19 (d), 7,209 from 19 (e) and 8,080 from 19 (f), drawn from the 20-patient response cohort. Blue indicates response and coral non-response. Repeated structures within a biopsy carry the same patient outcome; the outcome replication unit is the patient. The complete 25-scope structural series includes every lineage, lesion axis, gene module and observed coverage record. **g**, Local non-response-logit sensitivities in an alternative model that encoded communication as receiver-cell signal scores, fitted in the same 20 patients as Fig. 7 (11 R and nine NR). The network contains 14 ecological, 20 molecular, 16 pathology and two outcome nodes. Magenta and teal ribbons indicate positive and negative sensitivities, respectively; widths are normalized within each layer boundary. Niche colours follow Fig. 2. Labels specify cell identity, molecular feature, anatomy and spatial scope. The expanded network includes 85 molecular nodes and all 25 pathology features. **h**, All three significant enrichments at the high-importance end of the complete cell-state gene ranks of the alternative model: mitochondrial envelope and mitochondrion in Treg, and endosome in EC-AVR. Stem length shows NES and head area shows −log10(FDR). The two significant low-importance-end enrichments and all tests are documented in the study records for Supplementary Data 2. **i**, Fifty ligand–receptor interactions involving immune cells in the main model, with source and target cell states, complete molecular identity and structural scope. Colour shows the observed class mean before model imputation. Dot area shows absolute local sensitivity multiplied by the standard deviation at model evaluation. Labels give the complete-model importance rank (r), number of patients with positive observed scores (n, ranging from 1 to 3 of 20) and sign of the non-response sensitivity (+/−). The ring marks an undefined class mean. R, responder; NR, non-responder; PTC, peritubular capillary; TLS, tertiary lymphoid structure; t-IFTA, inflammation in atrophic tubules. Full scores, patient support and feature ranks are documented in the accompanying study records (Supplementary Data 2 and Source Data).

**Extended Data Fig. 10.**
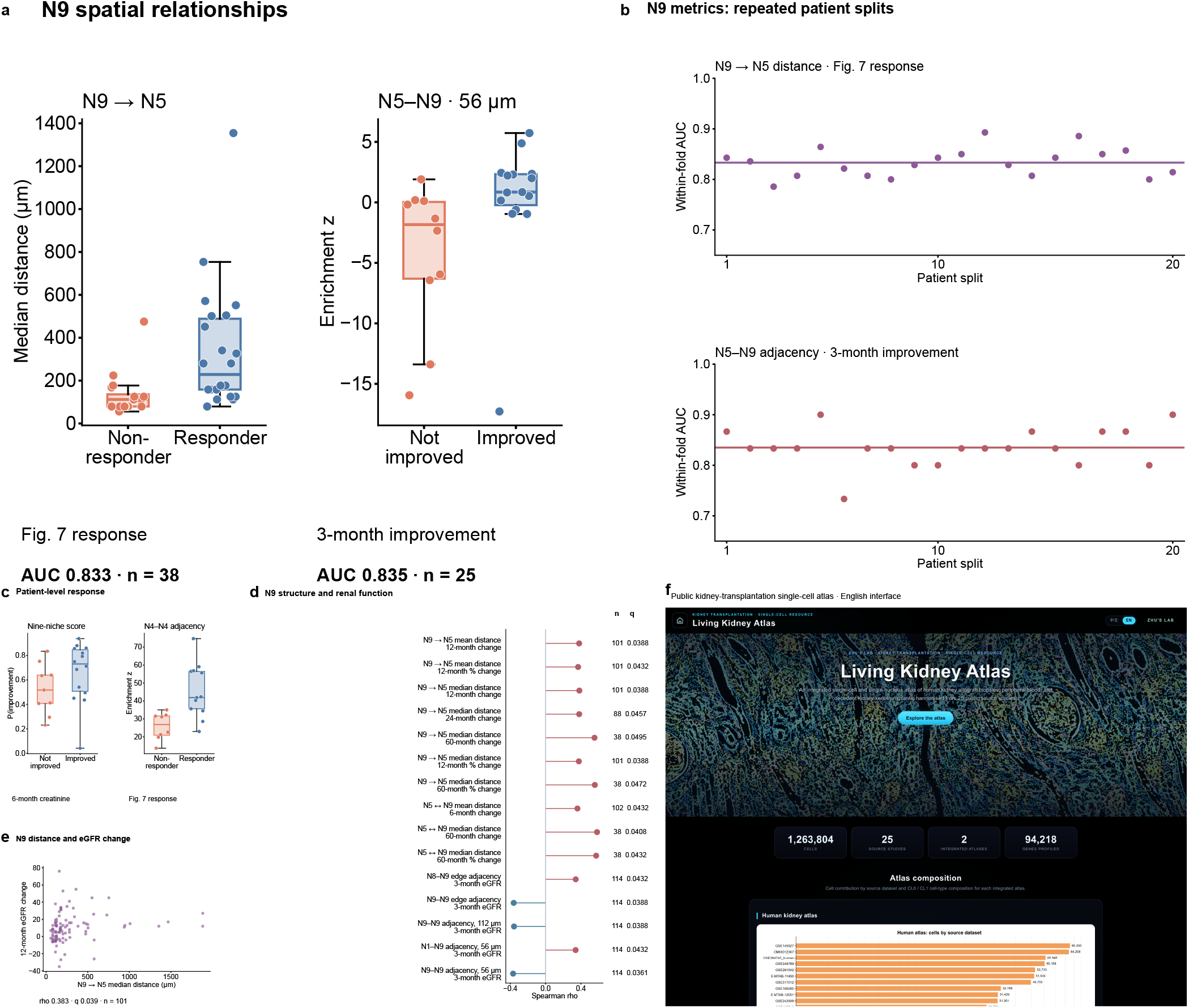
| Niche architecture, functional recovery and the public atlas. **a**, Median nearest N9-to-N5 distance and N5–N9 adjacency z, in 38 patients with the Fig. 7 response label and 25 patients with three-month improvement labels, respectively. Each dot represents a patient. **b**, Within-fold patient-pair-weighted AUC across 20 repeats of five patient folds for each metric; lines indicate repeat means. **c**, Patient-level held-out probabilities from the six-month nine-niche composite (23 patients) and N4–N4 adjacency z at 56 μm under the Fig. 7 response definition (21 patients). In a and c, boxes show medians and interquartile ranges; whiskers reach the most distant observations within 1.5 interquartile ranges. **d**, All 15 N9-related associations with reported eGFR or its change passing joint FDR<0.05. Rows give Spearman rho, patient count and q; correction spans the complete 920-feature and 23-endpoint family, with one selected biopsy per patient. Red and blue indicate positive and negative correlations. Models and features were selected after exploratory comparisons; predictions represent internal patient-fold evaluation. **e**, Patient-level N9-to-N5 median distance versus twelve-month absolute eGFR change (n=101; change in mL/min/1.73 m²). **f**, The interface (https://www.transplantation.com.cn/atlas/) provides access to integrated single-cell resources and a resource overview. Counts on the website describe the public resource; study cohort membership is defined in Methods. The page was accessed on 7 November 2026.

## Notes

### Competing Interest Statement

The authors have declared no competing interest.

### Funding

- National Natural Science Foundation of China: 82270789, 82241213
- Shanghai Municipal Education Commission: 24RGZNA02
- Science and Technology Commission of Shanghai Municipality: 09DZ2260300, S2025-001

